# Genomic forecasting links climate-associated variation to spring phenology in the European wild apple, a crop wild relative of the cultivated apple

**DOI:** 10.64898/2026.10.29.761786

**Authors:** Maxime Criado, Aurélie Mesnil, Somia Saidi, Anthony Venon, Komlan Avia, Johann Confais, Anamaria Roman, Tudor-Mihai Ursu, Dana Șuteu, Alexis Rodriguez, Thomas Kirisits, Mathieu Brisson, Frédérique Didelot, Arnaud Lemarquand, Stéphane Bazot, Gaëlle Vincent, Yann Bourgeois, Olivier François, Élodie Marchadier, Amandine Cornille

## Abstract

**Background:** Climate-associated genomic variation is often used as a proxy for adaptive potential, yet whether it reflects adaptation with phenotypic consequences, rather than demography alone, is rarely tested in long-lived species. Genomic offsets forecast climate maladaptation from genotype–environment models but are seldom validated against phenotypes. We combined whole-genome resequencing, SNP and transposable element insertion polymorphism (TIP) variation, landscape genomics, genome-wide association, genomic offset, and common-garden phenotyping in the European wild apple (*Malus sylvestris* Mill.), a crop wild relative with a complex demographic history, to test this.

**Results:** Of 148 resequenced accessions (135 wild, 13 cultivated), 89 wild individuals formed the core dataset. Population structure was associated with Last Glacial Maximum precipitation rather than contemporary climate, and genetic diversity increased from west to east. Genome–environment associations across 9.75 million SNPs identified climate-associated loci enriched in photoperiod-related flowering-time genes, with limited overlap with phenotype-associated loci. SNP-based genomic offset was associated with bud-break timing in the Saclay common garden (marginal R^2^ = 0.31), with higher offsets corresponding to earlier bud break, but not with growth or ecophysiological traits. This association depended on Austrian and Romanian populations and was clearest in non-admixed wild individuals. Offsets based on TIPs or glacial climate did not predict measured phenotypes.

**Conclusions:** Climate association, phenotypic relevance, and predictive power of genomic variation are separable. In a long-lived tree, genomic offset predicted spring phenology, a key adaptive trait, but this signal varied across provenances and ancestries and was absent for transposable elements. Genomic offsets can therefore forecast potential phenological maladaptation, provided they are validated against phenotypes such as bud break.

## Background

Climate change is rapidly altering temperature and precipitation regimes, creating a major evolutionary challenge for long-lived plant species whose generation times limit genetic turnover (1,2). Trees can buffer environmental change through phenotypic plasticity, but persistence over longer timescales also depends on standing genetic variation and evolutionary responses to local conditions (3). Identifying the genomic components associated with climate and testing whether they predict phenotypic responses under environmental change are therefore central goals of evolutionary and conservation genomics, particularly for understanding how perennial plants rapidly adapt to global change. Fruit trees exemplify this challenge: recurrent gene flow between wild and cultivated compartments means wild relatives have contributed substantially to cultivated genomes, making them valuable candidates for identifying climate-associated variation that may remain relevant to the cultivated gene pool. More broadly, crop wild relatives are not necessarily more genetically diverse overall than their cultivated counterparts, but domestication selects primarily on agronomic traits such as yield, taste, and storage rather than on climate resilience, so climate-relevant variation is not guaranteed to be retained in cultivated germplasm (4–6). Yet in perennial fruit trees, local climatic differentiation remains less well characterized than in annual plants, and the extent to which genomic predictions align with measured phenotypic responses remains uncertain (7).

Landscape genomics provides a framework for addressing this gap by linking genomic variation to environmental gradients. Genome–environment association (GEA) analyses identify variants whose frequencies covary with climate, while genomic offset summarizes the mismatch between genomic states predicted under different environmental conditions (8–10). Importantly, genomic offset is a predictive statistic rather than a direct measure of fitness loss; its biological relevance therefore depends on empirical validation. Moreover, interpreting genomic offset as a proxy for maladaptation assumes that contemporary genotype–environment associations reflect local adaptation, although demographic history and population structure can also shape these associations (11). This raises the question of whether genomic offset–phenotype relationships are consistent across ancestry backgrounds, particularly in individuals with admixed ancestry from gene pools associated with contrasting climatic conditions. Transposable element insertion polymorphisms (TIPs) represent a second, less-explored layer of structural variation that selection can also shape—including, potentially, climate-related selection—though their contribution to climate adaptation remains largely untested outside a handful of systems (12–14). A complementary approach to identify genotype–phenotype relationships directly is genome-wide association analysis (GWAS) (15), which tests for statistical association between genetic variants and measured trait variation without presupposing a link to climate. Combining GEA-based detection of climate-associated variants with GWAS-based detection of trait-associated variants allows us to assess the extent of overlap between these signals and ask whether climate-associated genomic variation is reflected in measured phenotypic differences.

The European wild apple, *Malus sylvestris* Mill., is well suited to test these ideas. It contributed to the domesticated apple genome, occurs across a broad European climatic range, and has a complex demographic history involving glacial refugia, postglacial recolonization, and recurrent introgression from cultivated apple (6,16,17). Gene flow between the two compartments is bidirectional, and adaptive variants originating in *M. sylvestris* have also been transferred into the cultivated gene pool (6), underscoring the breeding relevance of climate-associated variation segregating in the wild compartment. This demographic complexity raises questions about how demographic background, contemporary climate associations, and crop-to-wild gene flow shape genomic forecasts of climatic mismatch.

Here, we combined whole-genome sequencing, SNPs and TIPs, landscape genomics, genome-wide association analyses, and a common-garden network (Fig. 1). We first characterized population structure and its association with geographic, contemporary climatic, and Last Glacial Maximum variables. We then identified climate-associated variants and examined their functional and geographic distributions. Finally, we tested whether SNP-based genomic offset predicted phenotypic variation in the Saclay common garden and projected relative genomic mismatch under future climate scenarios. By combining SNP- and TIP-based signatures of climate-associated variation with independent phenotypic validation in a common garden and with historical climate legacy from the Last Glacial Maximum, we provide one of the first empirical tests of whether genomic offset generates phenotypically informative forecasts in a long-lived crop wild relative.

**Fig. 1.**
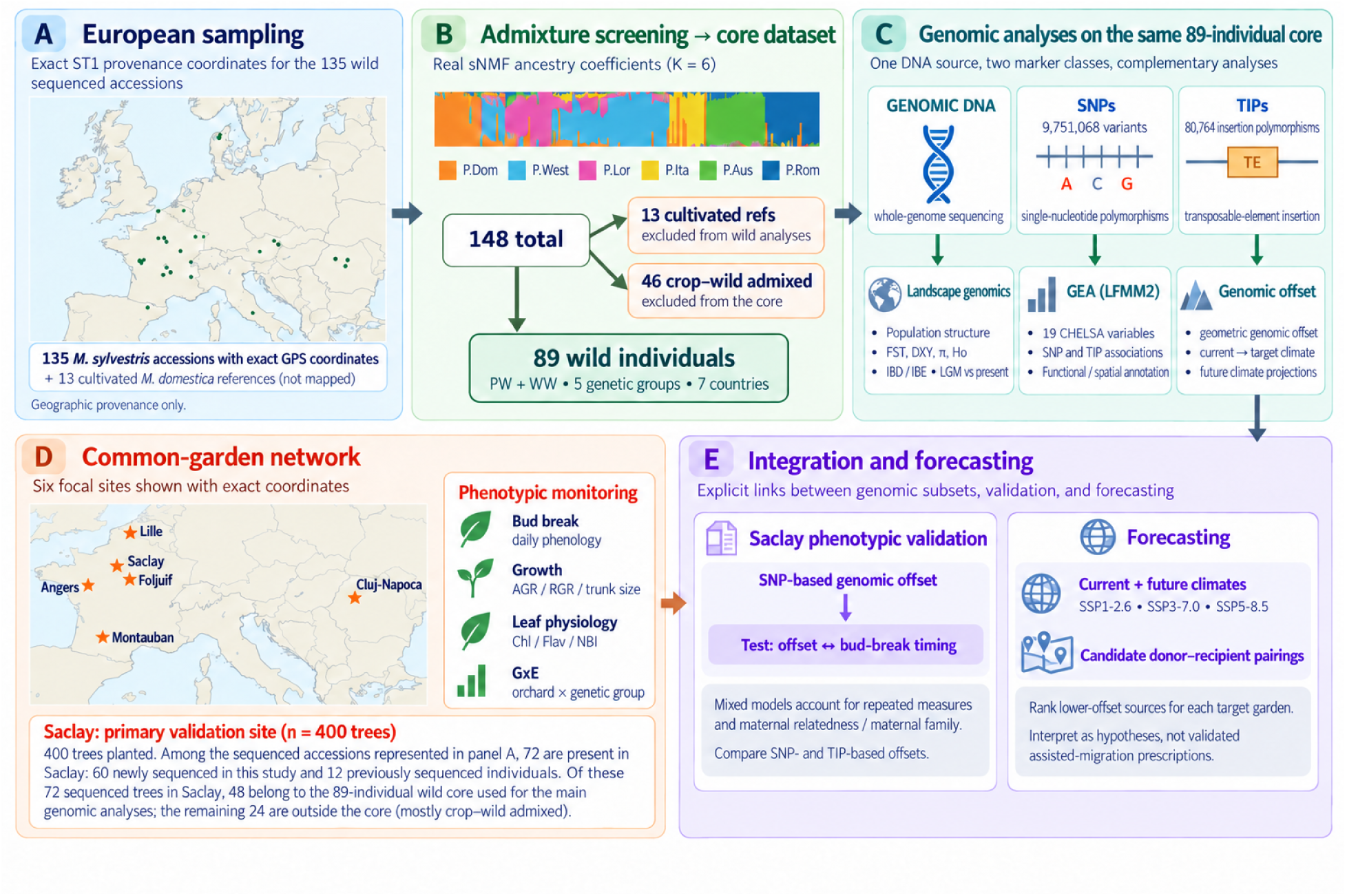
Study design and data flow, from European wild-apple sampling to genomic forecasting and common-garden validation. **(A)** Geographic provenance (exact coordinates, Suppl. Table 1) of the 135 resequenced *M. sylvestris* accessions; the 13 cultivated *M. domestica* references are not mapped. **(B)** Admixture screening of all 148 individuals with sNMF (K = 6; ancestry coefficients shown), separating the 13 cultivated references and 46 crop–wild admixed individuals from the 89-individual pure-wild/wild–wild-admixed core used for the core landscape-genomic analyses (5 genetic groups, 6 countries). **(C)** Genomic analyses performed on the same 89-individual core from a single whole-genome-sequencing DNA source: population genomics, genome–environment association (GEA, LFMM2) on 9,751,068 SNPs and 80,764 transposable-element insertion polymorphisms (TIPs), and genomic-offset modelling. **(D)** The common-garden network (eight orchards in total), with twelve phenotypic traits monitoring of bud break, growth, leaf physiology, and genotype-by-environment interactions. The Saclay orchard (n = 400 trees, primary validation site) includes 72 of the sequenced accessions from (A): 60 newly sequenced in this study and 12 previously sequenced (17). Of these 72 sequenced trees, 48 belong to the 89-individual wild core used for the main genomic analyses; the remaining 24 fall outside the core, mostly crop–wild admixed (Suppl. Table 1). **(E)** Integration and forecasting: SNP-based genomic offset against bud-break timing at Saclay in mixed models accounting for repeated measures and maternal relatedness, and use current and future genomic offset (SSP1-2.6, SSP3-7.0, SSP5-8.5) to rank candidate donor–recipient pairings, interpreted as testable hypotheses rather than validated assisted-migration prescriptions.

## Results

### Historical population structure provides the genomic background for climate-associated variation

We inferred population structure and admixture among 148 individuals, including 135 *M. sylvestris* accessions and 13 cultivated *M. domestica* references (Suppl. Figs. 1-2); Suppl. Tables 1–2). We identified six ancestry components using both fastSTRUCTURE and sNMF, including a cultivated cluster (*Pdom;* Suppl. Figs. 3-4). We classified individuals with >10% *Pdom* membership (see cut-off choice in Suppl. Fig. 5) as crop–wild admixed (CW) (Suppl. Fig. 5). Removing 46 CW individuals and the 13 cultivated references yielded 89 wild individuals for the core population-genomic and landscape-genomic analyses (Suppl. Fig. 6). Reanalysis of population structure for the 89 wild individuals supported five major genetic groups distributed across western Europe, Lorraine, Italy, Austria, and Romania (Fig. 2; Suppl. Figs. 7,8,9,10; Suppl. Table 3). This structure was concordant across fastSTRUCTURE, sNMF, PCA, Neighbor-Net, and kinship analyses (Fig. 2). For the 89 wild individuals retained for downstream analyses, sNMF cross-entropy was minimized at K=5 and increased at K=6 (Suppl. Fig. 9 B), while the fastSTRUCTURE cross-validation error plateaued from K=5 onward with broadly overlapping confidence intervals across K=5–K=10 (Suppl. Fig. 9 A); we therefore retained K=5 as the most parsimonious solution supported by both methods.

**Fig. 2.**
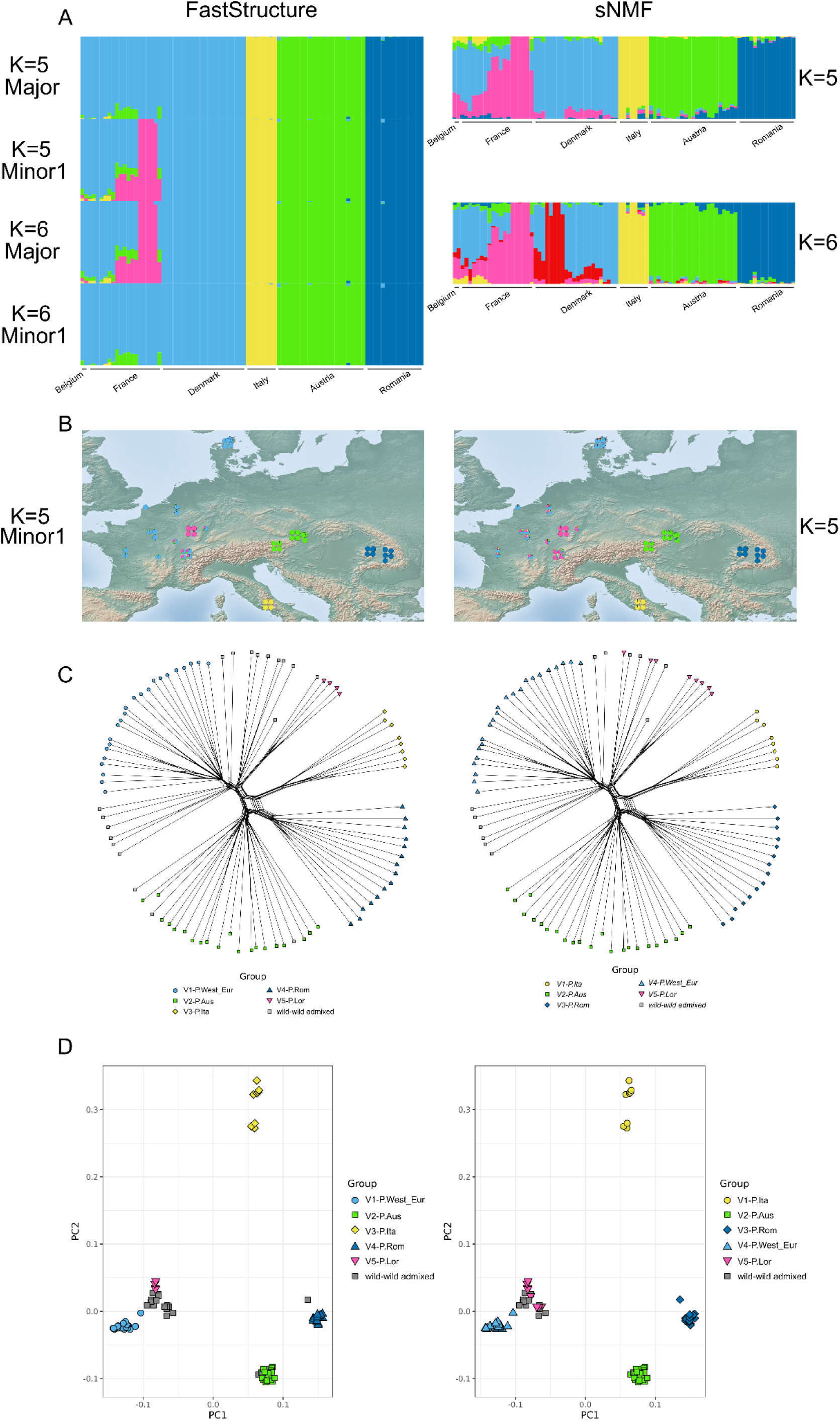
Population structure analysis of pure core European wild apple individuals (n = 89) inferred using sNMF and fastSTRUCTURE. **(A)** Ancestry bar plots obtained with fastSTRUCTURE (left; K = 5 and K = 6, Major and Minor clusters) and sNMF (right; K = 5 and K = 6). Colours correspond to the five inferred wild clusters: P.West_Eur (cyan), P.Lor (pink), P.Ita (yellow), P.Aus (green), and P.Rom (blue). **(B)** Geographic distribution of ancestry coefficients shown as pie charts for fastSTRUCTURE (K = 5, MinorCluster; left) and sNMF (K = 5; right). **(C)** SplitsTree representation of genetic relationships among individuals, with clusters inferred by fastSTRUCTURE (left) and sNMF (right) at K = 5. **(D)** Principal component analysis (PCA) based on fastSTRUCTURE (left) and sNMF (right) cluster assignments at K = 5.

Genetic differentiation was the weakest among western (France and Denmark) wild apple populations and the highest in comparisons involving Italian and Romanian populations (Fig. 3A). The maximum pairwise *F_ST_* was 0.211 between Italy and Denmark, and *D_XY_* showed the same broad geographic pattern.

**Fig. 3.**
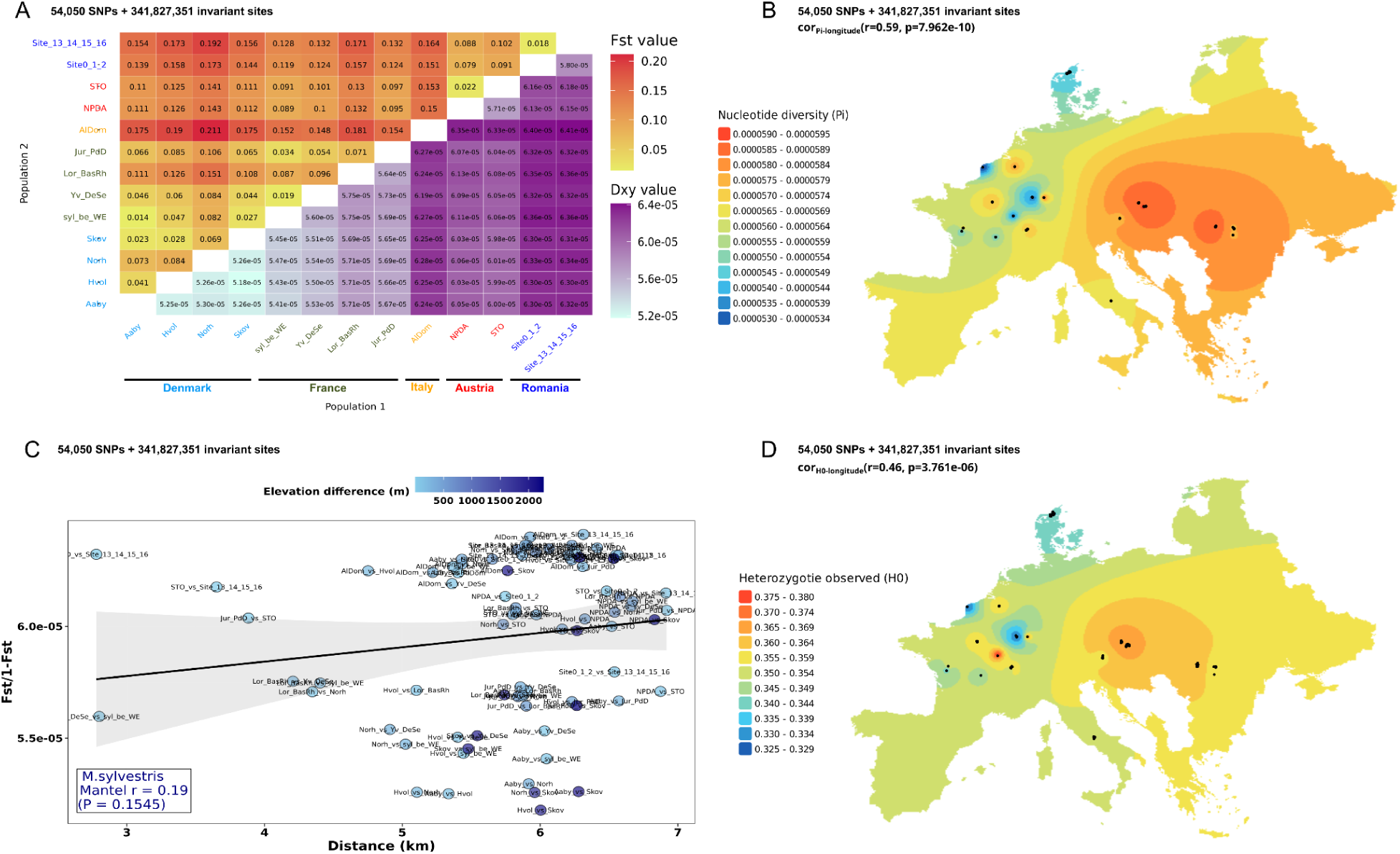
Landscape genomics analyses conducted on pure wild individuals (n = 89). **(A)** Pairwise ***F_ST_*** (top) and ***D_XY_*** (bottom) matrices estimated with *Pixy* using grouped sampling sites (**ST3**), aggregating individuals by site. **(B)** Nucleotide diversity (π) estimated with *Pixy* at the individual level. **(C)** Isolation-by-distance analysis among grouped sampling sites. **(D)** Observed heterozygosity (H_O_) estimated with *Stacks* at the individual level.

Patterns of nucleotide diversity and observed heterozygosity increased from western toward central and eastern populations, with the highest values in Austria and Romania (Fig. 3B–D; Suppl. Fig. 11). Because nucleotide diversity was estimated from a polymorphic, LD-pruned SNP set, π estimates are interpreted as relative differences among populations rather than conventional genome-wide estimates.

Geographic distance explained little of the observed differentiation: pairwise F_ST_/(1−F_ST_) was weakly and non-significantly correlated with geographic distance (Mantel r = 0.19, P = 0.1545; Fig. 3C), indicating no significant isolation-by-distance. We next tested whether spatial and 19 CHELSA bioclimatic variables (1981–2010) (18) or LGM climatic variables better accounted for population differentiation. Distance-based redundancy analyses including PCNM spatial predictors and CHELSA climate variables did not retain contemporary climate variables as significant predictors but identified Last Glacial Maximum precipitation during the driest month (Bio14-LGM) as the only significant climatic predictor (Suppl. Table 5).

### Contemporary climate-associated variation converges on seasonal phenology

We performed GEA analyses with LFMM2 (19) on 89 wild individuals using 9,751,068 high-quality SNPs (filtering to retain SNPs with MAF ≥ 0.05 and per-site missingness < 0.20) and 19 CHELSA bioclimatic variables. We included latent factors to account for population structure and unobserved confounding effects. Across individual climatic predictors, candidate SNPs clustered predominantly around precipitation of the coldest quarter (Bio19; 902,792 SNPs), mean temperature of the wettest quarter (Bio8; 57,460 SNPs), and annual precipitation (Bio12; 10,911 SNPs), whereas all other bioclimatic variables each yielded fewer than 6,000 significant SNPs (Suppl. Fig. 12 A-B). To isolate the most robust signals and control for potential redundancy among correlated predictors (Suppl. Fig. 13), we intersected loci detected for Bio8, Bio12, and Bio19 with candidate genes identified by a global model that simultaneously integrates all 19 environmental variables (All_Bio; Suppl. Fig. 12 B). Among these intersected loci associated with Bio8, Bio12, or Bio19, 341 non-redundant candidates fell within flowering-time gene models, corresponding to 100 candidate genes and 129 physical loci (Suppl. Table 8). Most of these genes were associated with a single bioclimatic variable, whereas 28% were shared across two or more variables.

Functional analyses did not identify significant Gene Ontology enrichment in the complete set of climate-associated candidate genes after multiple-testing correction. Cross-referencing climate-associated candidates with curated flowering-time resources recovered genes involved in photoperiod, circadian regulation, vernalization, sugar signaling, and related developmental pathways (Suppl. Figs.12 & 14; Suppl. Tables 6–9). Against the Plant Flowering-time Gene Database background (PFGD) (20), candidate flowering genes were distributed non-randomly across functional pathways (χ^2^ goodness-of-fit *P* = 9. 999×10^-5^), with the strongest enrichment in the photoperiod category (65 observed genes versus 28.81 expected; FDR = 8. 38×10^-19^). Single-variable candidate examples included VERNALIZATION1 (*VRN1)* for Bio19, CRYPTOCHROME 1 (*CRY1)* for Bio8, and FLOWERING LOCUS T (*FT)* for Bio12, while candidates shared across climatic variables included *RAP2.7/TOE1*, *UBP12*, *CDC73/PHP*, and *FPA* (Suppl. Table 9). Allele-frequency maps of these representative candidates showed spatially structured variation across Europe, including marked west–east or peripheral frequency shifts (Suppl. Fig. 15).

### Phenotypic variation across common gardens and genetic association mapping

We tested genotype-by-environment (*G*×*E*) variation across the common-garden network using DAPC genetic groups (Suppl. Fig. 16; Suppl. Table 10). Most growth and ecophysiological traits showed no significant *G*×*E* interaction (Suppl. Table 11). Bud break was the only measured trait with a significant orchard × genetic-group interaction (*P* < 0. 05), with the strongest contrasts observed between Saclay and Angers (Suppl. Fig. 17).

GWAS of common-garden phenotypes (Fig. 1) highlighted 327 SNPs (296 significant and 31 suggestive; LOD ≥ 3) and 330 haplotypes (325 significant and 5 suggestive) across the twelve evaluated traits (Suppl. Tables 12 & 13). Most associated variants were intergenic (>5 kb from annotated genes; 61.3%, 403/657; Suppl. Table 14). Although the distribution across structural categories differed from baseline genome expectations (χ^2^ = 37. 62, *df* = 3, *p* = 3. 41×10^−8^), the frequency of distant intergenic variants did not significantly deviate from its physical genome coverage (*p* = 0. 213, binomial test) and remained uniform across all evaluated traits (χ^2^ = 38. 47, *df* = 44, *p* = 0. 707). Functional annotation of intragenic GWAS candidates recovered several genes with known roles in chromatin regulation, sugar signaling, and stress responses, including *SU(VAR)3-9-RELATED PROTEIN 5* (*SUVR5*; AT2G21740), *SUCROSE SYNTHASE 4 (SUS4*; AT3G43190), and *TREHALOSE-6-PHOSPHATE SYNTHASE 1 (TPS1*; AT1G78580) (21). However, overlap with the curated flowering-time database was no greater than expected by chance (*P* = 0. 242).

At the SNP level, 26 SNPs were shared between GEA and GWAS signals across seven traits, compared to 46.28 expected under the reported hypergeometric null (Suppl. Table 15). Consistent with overall GWAS results, most of these overlapping loci were in intergenic regions (>5 kb). At the haplotype level, 196 GEA-associated SNPs fell within 87 GWAS haplotype intervals, which was also lower than expected under the null distribution (233.27 expected; *P* = 0. 00395; Suppl. Table 16).

### Genomic offset predicts bud-break variation in the Saclay common garden

We estimated genomic offset between the climatic conditions at each source population and those at the Saclay common garden using the genetic.offset function in LEA (22), with SNPs identified as climate-associated by LFMM2 (Fig. 4; Suppl. Table 17). Here, we treated genomic offset as a predictor of genomic mismatch rather than a direct measure of realized fitness loss. Genomic offset increased broadly with environmental distance from Saclay (Suppl. Fig. 18; r = 0.615, p-value = 1.321e-10). French populations near Saclay showed the lowest values (0.054–0.056), whereas Romanian, Austrian, and Italian populations showed higher values (0.76–0.89). Northern Danish populations remained relatively low despite their geographic distance. We then tested whether offset was associated with phenotypic traits measured at Saclay, the common garden with the highest density of sequenced individuals (Suppl. Table 1) and the most complete phenotypic records (Fig. 4A).

**Fig. 4:**
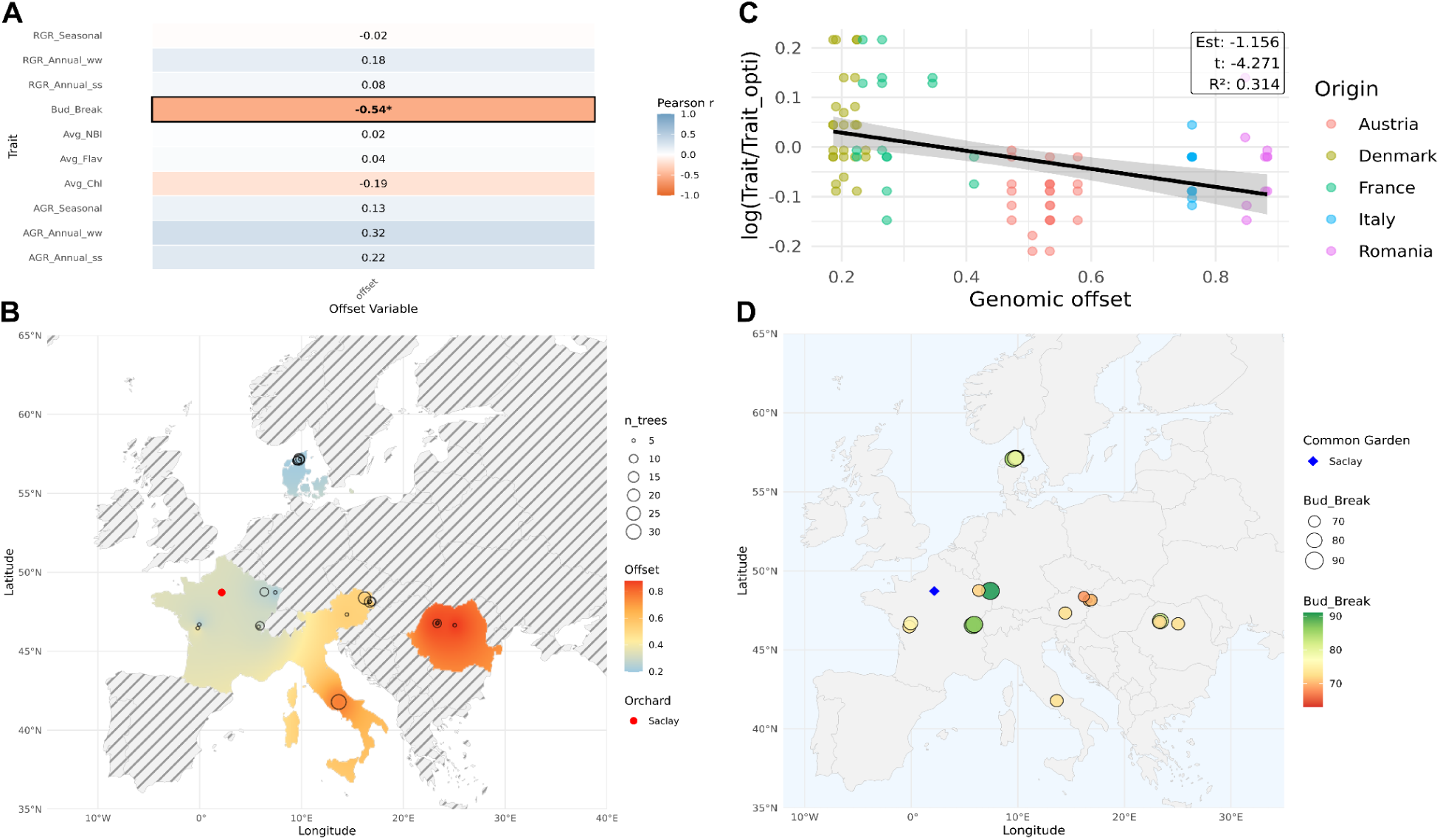
Phenotypic validation and spatial distribution of genomic offset predictions. **(A) Heatmap of correlation coefficients (r) between growth traits, fitness proxies, and predicted genomic.** The genomic offset predictions were performed using Saclay orchard as the target site. Statistically significant correlations are highlighted in black with an asterisk (*). **(B) Spatial distribution and interpolation of raw predicted genomic offset across Europe.** The red point highlights the target site (Saclay orchard). Open circles denote sampled populations, with circle size proportional to the number of sequenced individuals. Colors indicate raw predicted genomic offset values, calculated by comparing each population’s origin environment with the target environment at the Saclay orchard, ranging from blue (lowest offset) to red (highest offset). Spatial interpolation is restricted to regions containing sampled populations; unsampled areas are shown with grey hatching (see Methods for details on sampling coverage). **(C) Relationship between genomic offset and adjusted bud break timing.** Linear mixed-effects model regression demonstrating a significant negative association between genomic offset and adjusted bud break timing [log(bud_break/bud_break_optimum_); *r* =− 0. 54, marginal *R*^2^ = 0. 314)]. Points are color-coded by population or country of origin, and the shaded region represents the 95% confidence interval of the fixed effect. **(D) Geographic distribution of raw bud break phenology across European sampling sites.** Circle size and color gradient represent observed bud break timing measured for pure wild (PW) and wild-wild (WW) individuals at the Saclay orchard (marked by the blue diamond). Individuals are plotted at their original sampling sites, highlighting a distinct clinal gradient in spring developmental phenology.

Among the two statistical modeling approaches evaluated (Materials and Methods), the relative fitness formulation, log(Trait/Trait_optimum_), was retained based on superior goodness-of-fit parameters, exhibiting higher marginal/conditional *R*^2^ values and lower AIC scores compared to the raw squared trait distance models across all traits (Suppl. Table 17). Under this log-linear framework, most ecophysiological (chlorophyll, flavonoid content, and NBI) and growth traits (annual and seasonal AGR or RGR) showed weak, non-significant relationships with genomic offset (P>0.05; and negligible explanatory power R^2^_m_, Fig. 4A; Suppl. Table 17). In stark contrast, spring phenology (Bud_Break) exhibited a strong, highly significant linear relationship with genomic offset (R_m_^2^=0.314; Fig. 4C). Both linear (β=-1.16, p=0.00015) and quadratic (β=0.95, P=0.00087) terms were highly significant, indicating that higher genomic offsets were associated with earlier spring development at Saclay (Suppl. Table 17). By contrast, offset predictions evaluated using annual climate conditions rather than long-term 1981–2010 estimates failed to yield significant validation results (Suppl. Fig. 19).

We tested whether this relationship between relative fitness and genomic offset persisted after controlling for neutral population structure and environmental distance. We selected models across nested covariates. Bud_Break was the only trait for which the optimal model (lowest AIC = -180.56) retained genomic offset alongside environmental distance and population structure PCs (PCA model; β=0.3035, P=0.04782). Furthermore, genomic offset remained a significant predictor of bud breaks in the baseline model (p < 0.001), the environmental control model (p < 0.001), and the full PCA-adjusted model (p = 0.0478; Suppl. Table 18).

Because the positive effect size (β=0.3035) was sensitive to spatial structure, we conducted a leave-one-group-out (jackknife) sensitivity analysis across the five geographic origins. The association proved sensitive to group composition: excluding Austria (β=-0.088, P=0.7195) or Romania (β=-0.1185, P=0.5703) eliminated statistical significance, whereas excluding France preserved a strong positive effect (β=0.4651, P=0.0122; Suppl. Table 19).

Mapping genomic offset and phenological variation across Europe revealed a strong geographic pattern (Fig. 4B–D), where populations from Romania, Austria, and Italy exhibited the highest genomic offsets and earliest bud-break phenotypes. This phenological correspondence provides empirical support for the biological relevance of the genomic predictions.

While phenotypic validation relied on sequenced accessions maintained mostly at the Saclay orchard (Fig. 1), evaluating genomic offsets across all orchards (Suppl. Fig. 16) allowed us to identify candidate donor–recipient pairings, with donor selection stability confirmed via non-parametric bootstrapping (B = 1,000) and random-donor permutations (10,000 draws). Targeted translocations significantly reduced climate mismatch compared to unguided donor selection (global permutation P = 9.999 × 10⁻^5^; per-orchard P_perm_ ≤ 0.033). Suitability varied considerably across candidate donor sources. Orchards such as Saclay, Foljuif, Sovata, Angers, Lille, and Montauban paired effectively with candidate donors, yielding low residual genomic offsets (GOmin = 0-0.75). Translocation provided clear suitability gains over local baselines for Lille (ΔGO = 0.252, achieving a minimum offset of 0.326 [95% CI: 0.326, 0.451]), Foljuif (ΔGO = 0.052), and Montauban (ΔGO = 0, without a local candidate). Conversely, Saclay and Sovata showed zero offset reduction (ΔGO = 0.000), confirming that their local genotype (A038, P41C3_L1P4, Table S1) remains optimal without intervention (Suppl. Table 20).

In contrast, southern and coastal locations like Maison Estuaire in Brittany (GOmin = 1.98; 95% CI: [1.98, 2.056]) and Avignon (GOmin = 2.73; 95% CI: [2.73, 3.03]) retained high residual offsets post-optimization, highlighting donor pool limitations in extreme environments. However, these sites achieved the largest net offset reductions (ΔGO = 0.75 for Maison Estuaire; ΔGO = 0.48 for Avignon). Bootstrapping confirmed moderate-to-high rank stability for primary donor designations (P_top_ = 0.658 for Avignon donor C06; P_top_ = 0.0.653 for Maison Estuaire donor D08; Suppl. Table 20). These metrics show that while assisted migration substantially mitigates climate mismatch across most sites, extreme climate non-analogs remain vulnerable (Suppl. Fig. 20 A).

### Ancestry-dependent genomic offsets

Genomic offset models generally assume that contemporary genotype–environment associations reflect local adaptation, although demographic history and population structure can also confound these associations (11). To control for background population structure—including cultivated gene flow—GEA and genomic offset estimation were conducted using *K* = 6 latent genetic clusters across 135 individuals, explicitly accounting for pure wild (PW; n=76), wild–wild admixed (WW; n=13), and crop–wild admixed (CW; n=46) lineages (9,875,620 SNPs; Suppl. Fig. 21, Fig. 1). We then tested whether relationships between genomic offset and phenotypic traits differed among these ancestry groups. Across chlorophyll, NBI, and flavonoid traits, neither ancestry main effects nor offset × ancestry interactions were statistically significant (*P* > 0. 05), despite moderate within-group correlations. Bud breaks provided the clearest ancestry-dependent response. PW individuals exhibited a strong negative correlation with genomic offset (*r* =− 0. 59), a significantly higher baseline relative log-distance from the optimum (β = 0. 140, *P* = 0. 0086), and significant linear (β =− 1. 07, *P* = 0. 0196) and quadratic (β = 1. 37, *P* = 0. 0251) offset × status interaction terms. Corresponding significant interaction effects were absent for CW (*r* =− 0. 18) and WW (*r* =− 0. 46) individuals (Suppl. Table 21).

Growth traits also revealed significant ancestry-dependent dynamics under the log-linear LMER models. Baseline log-deviations from optimal growth were significantly lower in PW trees compared to CW hybrids across several growth parameters (AGR_Annual_ww: β =− 0. 700, *P* = 0. 0453; AGR_Seasonal: β =− 1. 75, *P* = 0. 0043; RGR_Seasonal: β =− 1. 30, *P* = 0. 0110). Crucially, seasonal absolute growth rate (AGR_Seasonal) exhibited a significant positive linear interaction with genomic offset specifically in PW individuals (β = 10. 3, *P* = 0. 0422; within-group *r* = 0. 24), indicating that seasonal growth accelerates under greater climate mismatch (Suppl. Fig. 21; Suppl. Table 21).

In contrast, large raw (simple, unadjusted bivariate; as opposed to the LMER-based interaction estimates) correlations observed within the WW subgroup for annual growth metrics (e.g., RGR_Annual_ss: *r* =− 0. 97; AGR_Annual_ss: *r* =− 0. 78; Suppl. Table 21) lacked significant LMER interaction terms; given the limited subgroup sample size, we treat these correlations as descriptive.

### Historical climate associations persist, but palaeoclimatic offset does not predict contemporary phenotypes

To assess whether climate-associated variation detected using Last Glacial Maximum conditions overlaps contemporary signals, we compared LGM and present-day GEA results. For Bio14-LGM, 679 SNPs were significant, 294 of which also appeared in the contemporary global environmental set (hypergeometric P < 10−16). At the genome-wide level, 328,123 SNPs were shared between the LGM and present global models, corresponding to 74–76% of the respective candidate sets.

### TIP climate-associated variation, but no improvement of genomic forecasting

GEA analysis across all environmental variables identified 2,984 climate-associated TIPs out of 80,764 genotyped markers. However, TIP-based genomic offsets did not correlate with measured common-garden phenotypes, contrasting with the SNP-based offset–bud break association.

Variable-specific GEA analyses revealed distinct association patterns for TIPs compared to SNPs. Across individual bioclimatic variables (Bio01–Bio19) and the combined environmental dataset (All_bio), a total of 5,368 TIPs and 1,451,380 SNPs were identified as climate-associated. Proportionally, TIPs were significantly less enriched for climate associations than SNPs (6.65% vs. 14.70%; Odds Ratio *OR* = 0. 41, *p* < 2. 2 × 10^−16^). Furthermore, the distribution of candidate TIPs across genomic structural categories differed significantly from that of SNPs (χ^2^ = 94. 22, *df* = 3, *p* < 2. 2 × 10^−16^; Suppl. Fig. 22), exhibiting depletion within gene bodies and enrichment in gene-proximal intergenic regions (< 5*kb*)—likely reflecting selective constraints against disruptive coding insertions.

Neither primary genome annotations nor broader BLAST homology searches against PFGD (20) revealed significant pathway enrichment among gene-proximal climate-associated TIPs (hypergeometric *P* = 0. 981; pathway distribution χ^2^ = 9. 9151, simulated *P* = 0. 1671). Only nominal, non-significant tendencies were observed in the sugar (3 observed vs. 0.98 expected; *P* = 0. 0609, FDR = 0.4869) and photoperiod pathways (3 observed vs. 1.44 expected; *P* = 0. 1565, FDR = 0.6260).

Complementary GWAS using Fast3VmrMLM (23) across single-site (100 TIPs), haplotype (129 TIPs), and mQTL (62 TIPs) frameworks identified only a single variant overlapping with GEA candidate regions: *MdoRA.svim_asm.INS.2031* (chr2:11,288,317; hypergeometric depletion *P* = 0. 722). This 183-bp insertion served as a left-flanking marker in a haplotype-level GWAS signal and showed strong climate association, confirming that TIPs driving environmental adaptation are largely distinct from those underlying measured traits.

Flanking alignments (4-kb) confirmed that *MdoRA.svim_asm.INS.2031* lacks the miRNA172 coding region, establishing it as an independent MITE insertion with high homology to pri-miRNA172p (24) (terminal inverted repeats 11–12 bp; Suppl. Figs. 23–24). Across pangenome assemblies (25), the locus was resolved in 41 of 42 *Malus* genomes across three configurations: unoccupied (14 bp; *n* = 26), MITE insertion (215–216 bp; *n* = 13), and intermediate intervals (57 and 79 bp; *n* = 2). A duplicated 9-bp target site duplication in MITE-containing alleles supports an ancestral unoccupied state. Phylogenetically, the insertion occurred in all *M. orientalis* (3/3) and *M. sieversii* (2/2) assemblies, was polymorphic in *M. domestica* (8/23; 34.8%), and was absent in *M. sylvestris* (0/10; Suppl. Fig. 25), indicating a pre-domestication origin in wild Central Asian and Caucasian lineages followed by introgression or incomplete lineage sorting.

### Future genomic offset is spatially heterogeneous but sensitive to projection uncertainty

Future climate projections revealed substantial heterogeneity in predicted vulnerability across the European range of *M. sylvestris* (Fig. 5): they exhibited relatively low genomic offsets under high-emission scenarios but emerged as among the most vulnerable groups under climate stabilization pathways. We also evaluated donor–recipient combinations minimizing genomic offset for common-garden sites under late-century SSP5-8.5 climates (Suppl. Fig. 20 B). Permutation tests confirmed that targeted seed transfers significantly reduce genomic offset relative to random selection (p < 0.05). Bootstrap iterations showed strong convergence toward key donor lineages: population D15 proved optimal for six of eight locations (Rank 1 probability 61.6% – 65.3%), while donor D11 / E10 / E13 yielded the greatest offset mitigation at Sovata (ΔGO = 1.96). In contrast, Montauban derived negligible benefit (ΔGO = 0.013). Crucially, even after optimization, residual genomic offsets across sites remain high (e.g., MinGO = 8.87 at Sovata, 7.88 at Avignon, and 7.5 at Montauban; Suppl. Table 23).

**Fig. 5:**
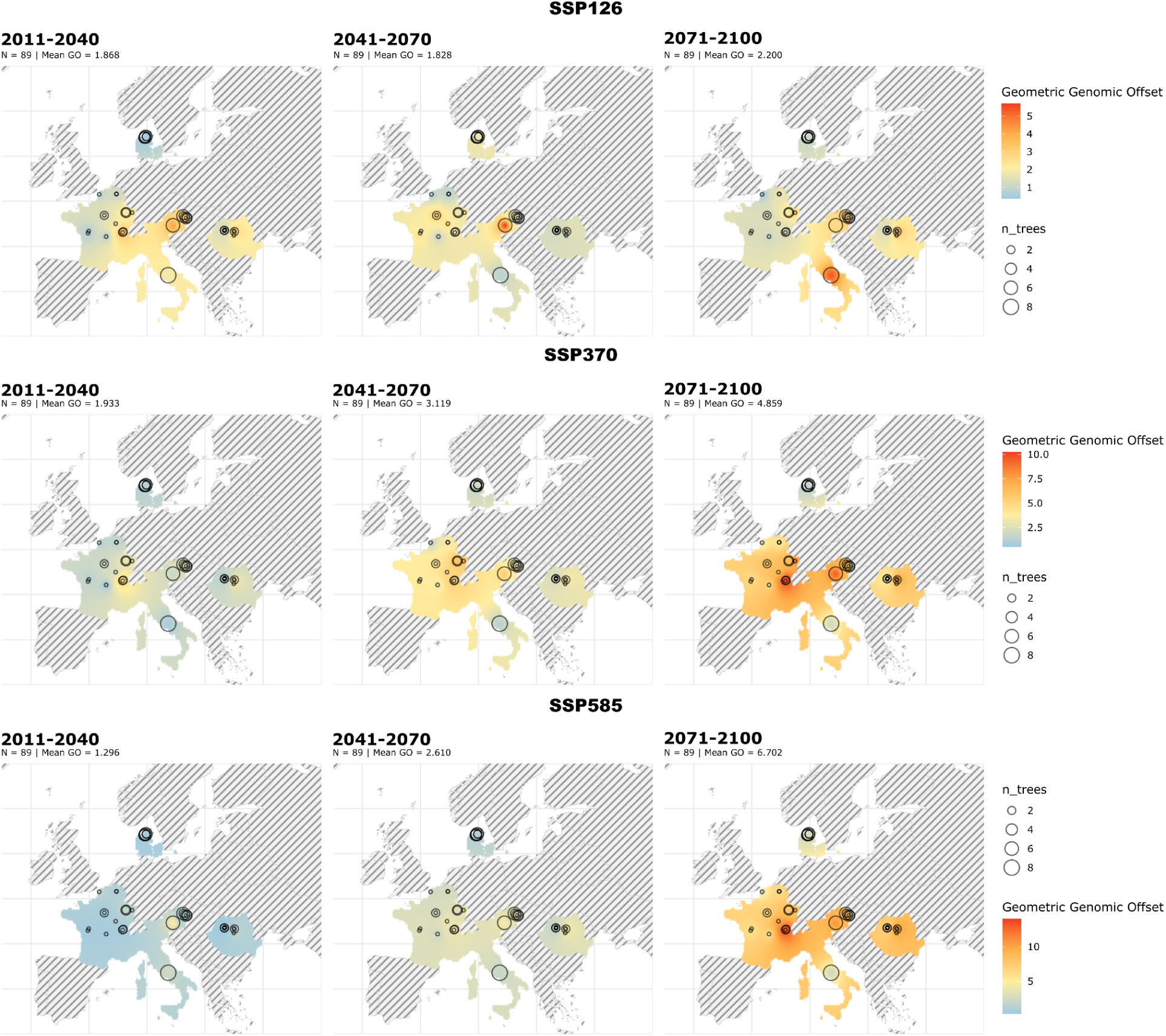
Projected genomic offset of *Malus sylvestris* across its native range under future climate scenarios. Genomic offsets for the European wild apple (*Malus sylvestris*) were estimated across three Shared Socioeconomic Pathways (SSP1-2.6, SSP3-7.0, and SSP5-8.5) and three future timescales (2011–2040, 2041–2070, and 2071–2100). Climate projections were extracted from the CHELSA database using the IPSL-CM6A-LR Earth System Model (83). Predicted offset values are indicated by a color gradient, ranging from blue (lowest predicted genomic mismatch) to red (highest predicted genomic mismatch). Circle sizes represent the number of individuals sampled at each coordinate.

## Discussion

By combining landscape genomics, SNP and transposable element variation, historical and contemporary climatic associations, and common-garden phenotyping, this study asks how different lines of genomic evidence relate to climate adaptation in a long-lived crop wild relative, *M. sylvestris*. Our results distinguish among genomic variation associated with climate, variation that is phenotypically relevant, and variation that predicts across contexts. These three properties need not coincide, and conflating them risks overstating what genomic forecasts can tell us about future maladaptation. This study provides a rare empirical test, in a wild, long-lived, outcrossing fruit tree with a complex demographic history, of how far genomic signatures of climate association translate into measured phenotypic responses — a question addressed so far mostly in herbaceous or short-generation systems. The same distinction likely matters for other perennial crop wild relatives, where direct selection experiments are impractical and genomic forecasting is often the only tractable way to anticipate climate vulnerability.

### Seasonal development as a recurrent axis of climate association

Climate-associated candidates included regulators of seasonal development, consistent with bud break being the only measured trait showing a significant genotype-by-environment response across the common gardens. The strongest functional enrichment was in photoperiod-related genes within the curated flowering-time candidate set, whereas the full set of climate-associated genes showed no significant Gene Ontology enrichment after multiple-testing correction. This convergence at the level of a broad developmental process is consistent with evidence from other forest trees, where developmental timing is a recurrent axis of climatic differentiation (26–28), including association mapping in Sitka spruce linking bud-set timing and cold hardiness to climatic variables (29).

Candidate annotations suggest several routes through which environmental conditions could influence developmental timing. The CRY1-annotated locus associated with mean temperature of the wettest quarter (Bio8) is particularly informative: in Arabidopsis, the CRY1 blue-light receptor also modulates temperature-induced growth through PIF4 (30). The UBP12 candidate offers a second possible link between seasonal development and abiotic responses: Arabidopsis UBP12/UBP13 regulate circadian rhythms and photoperiodic flowering and also influence abscisic acid signaling and drought responses through protein deubiquitinating (31,32). These functions nominate regulators that could integrate developmental and environmental signals, rather than genes whose relevance is restricted to flowering.

Other candidates point to control of developmental transitions at transcriptional and post-transcriptional levels. The VRN1-annotated locus associated with precipitation of the coldest quarter (Bio19) is a candidate for cold-related developmental regulation: Arabidopsis VRN1 participates in maintaining vernalization-induced repression of *FLC* (33). The FT-annotated locus associated with annual precipitation (Bio12) and the RAP2.7/TOE1 candidate further implicate regulation of the floral transition; in Arabidopsis, TOE1 represses *FT* expression (34). CDC73/PHP and FPA extend these hypotheses to chromatin-associated transcription and RNA 3′ end processing, respectively (35,36). Together, these annotations suggest that climatic differentiation could involve differences in when developmental programs are activated or maintained. However, association with a precipitation variable does not establish direct water sensing, nor does it identify the physiological cue acting on a locus.

To assess the relevance of these annotations to spring phenology, distinguish the floral transition from vegetative bud break. FT-like genes have experimentally supported roles in cultivated apples: *MdFT1* and *MdFT2* differ in their expression patterns, and *MdFT1* overexpression promotes precocious flowering (37). This supports the developmental relevance of the FT family in *Malus*, but does not establish which paralogue our candidate represents or whether its alleles control bud break. Moreover, controlled experiments in apple found that low temperature, rather than photoperiod, regulated growth cessation, dormancy induction, and release (38). Annotations assigned to photoperiodic flowering pathways therefore motivate tests of broader seasonal regulation, rather than implying that environmental cue responses are identical in Arabidopsis and apple. Expression profiling and allele-specific expression analyses during chilling and subsequent bud growth would help prioritize these candidates for functional validation.

### Climate association, trait association and prediction do not coincide

Apart from seasonal development—which underlies both climate and trait signatures—climate association, trait association, and predictive value are largely separable properties of genomic variation. Overlap between climate-associated (GEA) and phenotype-associated (GWAS) loci was limited at the level of individual variants. GWAS nevertheless recovered candidates annotated as TPS1 and SUVR5. In Arabidopsis, TPS1 links trehalose-6-phosphate signaling to flowering, whereas SUVR5 participates in chromatin-mediated repression of stimulus-responsive genes (39,40). These are phenotype-associated candidates, not evidence that the same loci mediate our climate associations.

The limited overlap may reflect the different questions these methods ask — GEA identifies variants covarying with environmental gradients against a background of population structure (41), whereas GWAS detects variants associated with measured phenotypic variation in a given environment — combined with the expected consequences of a polygenic architecture dominated by small-effect loci (42) and limited power to detect the same loci twice independently. This limited locus-level overlap does not contradict the phenological convergence described above; rather, it suggests that convergence is better supported at the level of biological processes than at the level of specific causal variants, which should be treated as hypothesis-generating candidates rather than demonstrated mechanisms. A polygenic explanation remains a hypothesis, and geographic structuring of allele frequencies may reflect demography as much as selection.

TIPs illustrate the same distinction. TIPs provided a complementary layer of climate-associated genomic variation, with a genomic distribution distinct from that of SNP-based candidates, yet TIP-based genomic offset did not predict any of the common-garden phenotypes measured here. In *Arabidopsis thaliana*, natural variation in TE insertions has been associated with climatic conditions, supporting the potential relevance of this variation to environmental responses (12). TE insertions can also modify gene expression and stress-responsive regulatory networks, providing mechanisms through which their effects could depend on the traits and environmental conditions examined. Such mechanisms remain untested in our study, and the absence of offset–phenotype associations cannot distinguish limited phenotypic effects from effects outside the scope of our experimental design. Our results therefore broaden the characterization of climate-associated variation in *M. sylvestris*, but do not support the predictive value of TIP-based offsets for the traits measured at Saclay.

### Genomic offset tracks bud break, but not uniformly across provenances and ancestries

Among the traits evaluated, SNP-based genomic offset was clearly associated with bud-break timing, whereas growth and ecophysiological traits showed little or no relationship with offset. This specificity suggests a trait-specific biological signal rather than a general offset-to-fitness relationship. Because we did not measure frost damage, survival, or reproductive output, we interpret this association as a phenological shift with potential fitness consequences rather than as demonstrated fitness loss — a distinction central to the broader debate on whether genomic offset can be treated as a proxy for maladaptation (43,44). The association was also not uniform across the sampled range: it did not hold when specific source regions were excluded, and should therefore not be read as a range-wide signal. We conducted phenotypic validation at a single common-garden site, limiting conclusions about spatial transferability; broader validation would require testing across multiple sites or using spatially explicit hold-out designs (10,44).

Another caveat concerns the temporal mismatch between climatic and phenotypic data: we estimated climate associations and genomic offsets from 1981–2010 climate normals, whereas phenotypes were measured in 2023–2026. Indeed, when we evaluated genomic offsets calculated using year-specific annual climate data for the measurement years, these metrics failed to yield significant phenotypic validation results. This lack of predictive power highlights that single-year weather events do not capture the primary selective drivers of local adaptation. Because we scored traits in a common garden, phenotypic differences primarily reflect genetic differentiation among source populations, and historical climate normals describe the selective environment that shaped these long-lived trees over multiple generations far better than recent inter-annual conditions do. If natural populations lag behind recent warming (27), this temporal mismatch should weaken, not inflate, genotype–climate–phenotype associations. However, garden conditions during the measurement years departed from long-term site norms, which could affect offset estimates for Saclay and modulate bud-break expression—a trait sensitive to inter-annual variation in winter chilling and spring temperature—although measurement year was accounted for in our main mixed models.

The offset–phenotype relationship also differed by ancestry status: it was clearest among pure wild individuals, weaker or absent among crop–wild admixed individuals, and inconsistent among wild–wild admixed individuals, for which apparent subgroup correlations were not supported by the fitted interaction models. We treat the crop–wild pattern as a hypothesis — that cultivated introgression may alter the relationship between climate-associated genomic variation and phenotype (45,46) — rather than as a demonstrated buffering mechanism, because our design does not isolate the physiological or regulatory pathway involved. This ancestry dependence implies that genomic offset estimates derived from admixed genetic backgrounds should not be assumed to carry the same phenotypic meaning as those from non-admixed individuals. This directly limits the use of genomic offset in systems where wild and cultivated gene pools are not cleanly separated, as in most crop wild relatives.

Taken together, these results indicate that genomic offset can forecast potential phenological maladaptation: offset predicted deviation in bud-break timing from the local optimum, a fitness-relevant component of climate adaptation in temperate trees. This predictive signal is, however, trait-specific, regionally contingent, and not yet linked to measured fitness loss, so bud-break phenotyping provides a tractable and informative validation step for genomic forecasts rather than a substitute for fitness measurements.

### Glacial history shapes genomic background, not phenotypic predictability

Broad population structure in European wild apples was more strongly associated with a historical (LGM) climate variable than with contemporary climate or geography, consistent with a lasting influence of glacial refugia and postglacial recolonization on present-day genomic background (17,47–50). Because our analysis is correlative and does not explicitly compare demographic models, this pattern should be read as a historical genomic correlate rather than causal evidence of a specific recolonization history.

Historical structure and contemporary climate association operate at different temporal scales: glacial history provides the broad genomic background, whereas contemporary GEA signals identify a narrower subset of loci covarying with recent climate after accounting for that background. Consistent with this distinction, LGM-based genomic offsets did not predict the common-garden phenotypes measured here. Historical climate legacy thus provides genomic context for interpreting present-day structure, but no additional forecasting power for contemporary phenotypic responses. Explicit demographic modeling would be needed to formally test specific recolonization scenarios.

### Future offsets and donor sources: hypotheses for assisted gene flow

Projected future genomic offset varied across climate scenarios and across the species’ range, indicating that different climate trajectories could reorder the relative genomic mismatch of populations rather than affect all populations uniformly. Regional patterns that diverged from the overall trend are potentially informative, but currently rest on a single climate model and projection pipeline; before being interpreted as evidence of pre-adaptation or refugial status, they would need to be tested for sensitivity to the choice of Earth System Model, emission scenario, and extrapolation beyond the sampled climatic envelope. In addition, future genomic offsets could be estimated for given sites to provide a more spatially explicit assessment of local genomic mismatch under projected climatic conditions and to identify sites that may be particularly vulnerable or retain relatively low levels of mismatch. We also obtained our offset estimates using a single modeling framework, and offsets can differ depending on the chosen framework (11,51). These projections should therefore be treated as one plausible estimate of relative mismatch rather than a uniquely robust one, and cross-method comparison would be needed to establish their sensitivity to this choice.

These projections are further constrained by the genomic resources and sampling design that underlie them. Variant detection relied on a congeneric reference genome, which may bias detection in structurally divergent regions, and geographic sampling was uneven across countries, with some regions contributing less to both population-structure inference and genotype–environment associations. More balanced sampling would strengthen confidence in the geographic patterns reported here.

Translating these projections into conservation guidance illustrates both the potential and the current limits of assisted gene flow under late-century warming. Candidate source populations reduced projected genomic offset relative to random selection, with the greatest apparent benefit for northern and north-western recipient sites and comparatively little benefit at the species’ southern range limit, suggesting that late-century warming there may exceed the adaptive buffer captured within the sampled gene pool. These donor–recipient pairings minimize relative genomic mismatch within the sampled gene pool; they do not, by themselves, demonstrate establishment success, ecophysiological tolerance or long-term persistence, and should be read as testable hypotheses for assisted gene flow rather than validated prescriptions.

## Conclusions

By integrating population genomics, genome–environment associations, GWAS, transposable element insertion polymorphisms, genomic offset and common-garden phenotyping, we show that the genomic background of European wild apple retains a signature of glacial climate, whereas contemporary (1981–2010) climate-associated variation converges on seasonal phenology. These results separate three often-conflated properties of genomic variation: association with climate, relevance to phenotype, and predictive power. In M. sylvestris, these properties align for SNPs and spring phenology: genomic offset predicted deviation in bud-break timing from the local optimum, indicating that offsets can forecast potential phenological maladaptation. This signal was, however, trait-specific, driven largely by Austrian and Romanian provenances, weaker in crop–wild admixed individuals, and absent for offsets based on TIPs or glacial climate. Genomic forecasts of climate adaptation therefore become informative when anchored to phenotypes such as bud break, and their generality now needs to be tested across sites, source populations, and genetic backgrounds (52). With these safeguards, genomic forecasting can help rank relative climatic mismatch among wild populations and inform conservation hypotheses, without treating genomic offset itself as a direct measure of maladaptation.

## Materials and Methods

### Plant material and genomic data acquisition

This study used 148 individuals, comprising 135 *M. sylvestris* accessions and 13 dessert cultivars as non-admixed cultivated reference genotypes (*M. domestica*) (6). Of the wild accessions, 60 individuals were newly sequenced, derived from flash-frozen leaves collected from trees planted in a wild apple orchard at Saclay (Université Paris-Saclay, France); the remaining wild individuals correspond to material described in (6,17), together representing seven European countries (Suppl. Table 1, Fig. 1). All individuals were sequenced using Illumina (6,17) and DNBSEQ™ technology (MGI) short-read technology (25× paired-end coverage).

For the 60 newly sequenced individuals, genomic DNA was extracted using 400 µL of Carlson lysis buffer (100 mM Tris-HCl, pH 9.5; 2% w/v CTAB; 1.4 M NaCl; 1% w/v PEG 8000; 20 mM EDTA). RNase A (10 µL; MN NucleoSpin® Plant II kit, Macherey-Nagel) was added to each sample, followed by incubation at 65°C for 10 minutes. An equal volume of chloroform was then added, and samples were vortexed and centrifuged at 11,000 × g for 10 minutes. The aqueous phase was transferred to a new tube, mixed with 450 µL of PC buffer, and loaded onto the NucleoSpin® Plant II column for purification according to the manufacturer’s protocol. DNA concentration and purity were assessed with a NanoDrop 2000 spectrophotometer (Thermo Fisher Scientific), retaining samples with A260/A280 ratios between 1.8 and 2.0 and A260/A230 ratios between 2.0 and 2.2. A total of 1.0 µg of DNA per sample was used for library preparation.

Sequencing was performed using DNBSEQ™ technology (MGI), based on DNA nanoball (DNB) sequencing. Fragmented double-stranded DNA was ligated to adapters and circularized into single-stranded circular DNA molecules, which were amplified by rolling circle replication to generate DNA nanoballs, minimizing amplification bias. DNBs were loaded onto patterned flow cells and sequenced using combinatorial probe-anchor synthesis (cPAS) chemistry with fluorescently labeled, reversibly terminated nucleotides.

All plant material used in this study was collected and used in compliance with applicable international regulations, including the Nagoya Protocol and the International Treaty on Plant Genetic Resources for Food and Agriculture (ITPGRFA). Appropriate permits and material transfer agreements were obtained for the collection, use, and analysis of both cultivated and wild apple genetic resources.

### SNP calling and filtering

Single-nucleotide polymorphisms (SNPs) were identified using a variant-calling pipeline based on GATK v3.8 (53), bcftools (54), and custom scripts. Individual gVCF files were generated using *GATK HaplotypeCaller* (55) and split by chromosome (chr01–chr17). For each chromosome, gVCFs were merged across individuals using *CombineGVCFs* and jointly genotyped with *GenotypeGVCFs* using the ‘--all-sites’ option to retain both variant and invariant positions. Initial hard filtering removed sites with fully missing genotypes (F_MISSING = 1) or invalid mapping quality annotations (MQ = NaN or Inf) using bcftools. Variant-level quality filters were then applied with GATK *VariantFiltration*, excluding sites with low mapping quality (MQ < 40), strand bias (Fisher Strand > 60 for SNPs or 200 for indels; Strand Odds Ratio > 3 for SNPs or 10 for indels), low quality-by-depth (QD < 2), or extreme ReadPosRankSum values (< –8 for SNPs or < –20 for indels). Only sites passing all criteria were retained using *SelectVariants*.

Filtered chromosome-level VCFs were concatenated using *bcftools concat* (54) to generate a genome-wide dataset. SNPs with more than 20% missing data (F_MISSING > 0.2) were excluded, and genotype-level filters removed genotypes with low depth (DP < 5), excessively high depth (DP > 100), or low genotype quality (GQ < 20). Depth and missingness thresholds were assessed on a random 1% subset of SNPs using custom R scripts. No individuals exceeded the 30% missing-data exclusion threshold, and all were retained.

Invariant and variant sites were processed separately to generate datasets suitable for diversity and differentiation analyses. Invariant sites were retained if they passed depth and missingness filters (DP 5–100, RGQ ≥ 20, F_MISSING ≤ 0.2). Variant sites were restricted to biallelic SNPs, filtered to remove SNPs within 10 bp of one another (--SnpGap 10), and subjected to the same depth, genotype quality, and missingness thresholds. Invariant and variant sites were concatenated to produce chromosome-level datasets that retained both site types, and an additional variant-only dataset, excluding monomorphic sites (AC = 0 or AC = AN), was generated for downstream analyses.

Genome-wide variant datasets were used to estimate genomic relatedness among individuals with PLINK2 (--make-king) (56). Duplicate or clonally related individuals were identified using the KING-robust estimator (kinship > 0.354) (57) and removed prior to population genomic analyses.

For population genomic analyses requiring neutral, independent markers, variants were further filtered for missingness (F_MISSING ≤ 0.2) and minor allele frequency (MAF > 0.05), and functionally annotated using SnpEff v5.0 (58) against the GDDH13 reference genome. Synonymous SNPs were extracted, and linkage disequilibrium (LD) pruning was performed using PLINK v1.9 (--indep-pairwise 50 5 0.2), yielding a final dataset of approximately independent synonymous SNPs used for population structure, genetic variation, and differentiation analyses.

Distinct SNP subsets were used according to downstream analytical objectives. Population structure analyses relied exclusively on LD-pruned synonymous SNPs to minimize the influence of selection and physical linkage. The same SNPs, merged with invariant sites, were used to estimate nucleotide diversity (π) at the individual level, genetic differentiation (*F_ST_*, DXY), and isolation by distance (IBD) at the sampling-site level (Suppl. Table 4). Isolation by environment (IBE), GEA, and genomic offset analyses were conducted using high-quality biallelic SNPs filtered for MAF > 0.05 and F_MISSING ≤ 0.2, excluding indels (Suppl. Fig. 26).

### Crop-wild admixture and population structure inference

Population structure was inferred from a dataset of synonymous, LD-pruned SNPs (N = 54,092) using two complementary clustering approaches. Ancestry proportions were first estimated using sNMF (59), a sparse non-negative matrix factorization method implemented in the R package LEA (22), and second using fastSTRUCTURE (60), a Bayesian model-based clustering algorithm. For both methods, the number of ancestral populations (K) was evaluated from K = 2 to K = 10, with 20 independent replicates per K to ensure robustness and reproducibility (Suppl. Figs. 3-4).

To assess crop–wild introgression, the 13 cultivated *M. domestica* reference genotypes (6) were included in all population structure analyses. Wild individuals showing a cumulative membership coefficient greater than 0.1 in the cultivated apple genetic cluster (*P.Dom*; Suppl. Fig. 1) were classified as crop–wild admixed (CW); individuals below this threshold were considered wild, including both pure wild (PW) and wild–wild admixed (WW) individuals. Individuals identified as CW by either method were considered CW and removed together with the 13 cultivated references (Suppl. Fig. 6). Population structure analyses were then repeated independently on the remaining wild individuals (n = 89), using only PW and WW SNPs (N = 54,050; Suppl. Fig. 26).

Because sNMF and fastSTRUCTURE differ in their underlying statistical frameworks, they also differ in the distribution of individual ancestry coefficients they produce, and method-specific membership thresholds were therefore applied, in this second clustering round, to assign individuals to genetic clusters: 0.99 for fastSTRUCTURE, whose maximum membership coefficients were strongly concentrated near 1 for non-admixed individuals, and 0.70 for sNMF, whose maximum membership coefficients showed a more continuous distribution extending well below 1 (Suppl. Fig. 9 C–D). Individuals at or above these thresholds were classified as pure wild (PW); those below were designated as wild–wild admixed (WW). The optimal number of clusters (K) was determined by integrating cross-validation and cross-entropy criteria, concordance between methods, and prior knowledge of population structure from previously published studies.

Phylogenetic relationships and patterns of genetic differentiation among individuals were explored using SplitsTree (61) and principal component analysis (PCA), based on the same set of synonymous, LD-pruned SNPs used for population structure analyses. In SplitsTree, uncorrected p-distances were used to generate a Neighbor-Net network depicting reticulate relationships and potential admixture among individuals (Fig. 2; Suppl. Fig. 28). PCA results were colored according to the population clusters inferred by sNMF and fastSTRUCTURE, allowing direct comparison between multivariate genetic patterns and model-based clustering results.

### Genetic diversity estimates

Genome-wide patterns of genetic variation were quantified using complementary summary statistics. Nucleotide diversity (π) (62), absolute genetic distance (*D_XY_*), and population differentiation (Weir & Cockerham’s *F_ST_*) were estimated with *Pixy* (63), using VCF files containing the LD-pruned population-structure dataset (synonymous sites) merged with invariant sites. Observed heterozygosity (H_O_) was calculated from the same dataset using the *populations* module of *Stacks*.

Before analysis, the VCF dataset was split by chromosome to reduce computational time. For each chromosome, diversity and differentiation metrics were calculated in non-overlapping 50-kb genomic windows and subsequently combined in accordance with *Pixy* guidelines. *D_XY_* and *F_ST_* were computed for each sampling site after regrouping individuals to standardize site sizes (Suppl. Table 4), and comparison matrices were generated to visualize pairwise estimates. Individual-level π and HO were spatially interpolated to visualize genome-wide patterns of genetic diversity across the landscape.

### Landscape genomics: isolation by distance and isolation by environment

Isolation by distance (IBD) was assessed by correlating pairwise genetic differentiation among sampling sites, computed as F_ST_/(1−F_ST_) from the population-structure dataset merged with invariant sites, against the corresponding geographic distances. The significance of this relationship was evaluated using a Mantel test with 10,000 random permutations.

Isolation by environment (IBE) was investigated using distance-based redundancy analysis (db-RDA) to model the relationship between genetic differentiation (*F_ST_*) and environmental and spatial predictors. Geographic distances underlying IBD were represented as vectors using positive eigenvalues of principal coordinates of neighbor matrices (PCNM) (64). Environmental variation was characterized using 19 bioclimatic variables from the CHELSA database (30-arc-second resolution) (18), representing annual and seasonal trends and extremes, averaged over 1981–2010, together with reconstructed values for the Last Glacial Maximum (∼21,000 years ago) (65).

A full db-RDA model, implemented with the *capscale* function in the vegan R package (64), included all spatial (PCNM) and environmental variables. Model selection was performed using the *ordistep* function in vegan to identify the subset of variables that best explains the observed genetic structure. The relative contribution of spatial versus environmental factors was quantified using variance partitioning with the *varpart* function (64).

### Genome–environment association analysis

To characterize the genomic basis of local climate adaptation in *M. sylvestris*, GEA analyses were conducted using the *lfmm2* function within the LEA R package. Two genomic datasets were evaluated: one containing only PW and WW individuals (n = 89; 9,751,068 SNPs) and one additionally including CW individuals (n = 135; 9,875,620 SNPs). Both datasets were filtered to retain only variants with MAF ≥ 0.05 and per-site missingness (F_MISSING) below 0.20.

Because missing data can bias association tests, genotype imputation was performed using the *impute* function from LEA, using ancestry and genotype frequency estimates derived from an initial sNMF analysis. The number of ancestral populations (K) was set to 5 for the PW/WW dataset and 6 for the PW/WW/CW dataset, selecting the run with the lowest cross-entropy across 20 repetitions; the most likely genotype was then assigned using the ‘mode’ method. Imputation was performed independently for each chromosome to maintain computational efficiency.

GEA was performed using latent factor mixed models (LFMM) to account for unobserved environmental confounders and population structure. To eliminate redundancy from linkage disequilibrium (LD) during latent factor estimation, the dataset was pruned using PLINK v1.9 (--indep-pairwise 50 5 0.2; 50-SNP sliding window, 5-SNP step size, r^2^ > 0.2 threshold), yielding 940,231 independent loci for the PW/WW dataset, and 890,788 for the CW/PW/WW dataset. Latent factors were computed on this pruned set using the lfmm2 least-squares algorithm, setting K = 5 based on population structure analyses excluding CW individuals. These latent factors were then incorporated into the lfmm2.test function and applied to the full, non-pruned datasets (full = TRUE) to compute joint p-values across all 19 CHELSA bioclimatic variables (1981–2010) simultaneously. This full dataset—comprising 9,751,068 (n = 89) and 9,875,620 (n = 135) SNPs—was filtered solely for minor allele frequency and missingness, retaining all variants for downstream genomic offset calculations. Because the global LFMM2 test showed an excess of p-values close to 1, we excluded p-values > 0.9 before q-value estimation as an empirical filtering step intended to reduce the influence of strongly over-conservative test statistics, enabling accurate false discovery rate (FDR) control. Climate-associated SNPs were subsequently identified using the q-value package at a significance threshold of q < 0.05.

### Genomic characterization and functional annotation of environment-associated loci

While *lfmm2.test* with full = TRUE provides the most robust identification of climate-associated loci, the analysis was also run with full = FALSE to characterize functional implications, calculating p-values separately for each environmental variable and thereby mapping specific adaptive regions to distinct climate factors. Both sets of significant SNPs were mapped to the *M. domestica* GDDH13 reference genome (GDDH13 v1.1) (66). The genomic position of each SNP was categorized as intragenic (CDS, mRNA, 3’UTR, or 5’UTR) or intergenic, with intergenic SNPs further partitioned by proximity to the nearest gene (within 2 kb, 2–5 kb, or beyond 5 kb).

Gene density was quantified for each bioclimatic predictor to identify the variables harboring the most candidate genes. To prioritize the most robust adaptive signals, variable-specific gene sets were intersected with genes identified in the global LFMM2 estimation (full = TRUE, All_Bio). This conservative filtering retained only SNPs repeatedly identified across independent runs, minimizing false positives and highlighting genes mediating broad climatic adaptation rather than isolated variable-specific noise. For these overlapping gene sets, gene ontology enrichment analysis was performed using the clusterProfiler R package and a custom ontology framework (67). A custom organism database (OrgDb) was built with AnnotationForge using the complete GDDH13 gene set as background, integrating Phytozome GDDH13 genome annotations with *Arabidopsis thaliana* (TAIR10) GO terms transferred via BLASTP homology (best BLAST hits with ≥ 50% identity and ≥ 70% alignment coverage).

To investigate the specific phenological drivers of adaptation, candidate genes were cross-referenced against an annotated flowering-time gene database (6) and against the PFGD database (20) via BLAST search (*E*-value ≤ 10⁻^5^, identity ≥ 80%, top hit), to identify potential orthologs involved in circadian rhythm, vernalization, or other flowering-time-related processes.

### Genomic offset estimation under current environmental conditions and phenotypic validation

To estimate the genomic mismatch of European wild apple populations under shifted climatic conditions, genomic offset was calculated using the genetic.offset function in the R package LEA. This metric quantifies the mismatch between an individual’s current genetic composition in its local environment and the genomic state predicted under the target environment—here, the climatic conditions of the experimental common gardens. We used the geometric genomic offset — a proxy for the squared difference between a fictive adaptive trait value and its local optimum in a given environment — computed across genomic loci discovered in the GEA study. Only SNPs significant (430,743 of 9,751,068) after FDR correction were used to predict genomic offset. By focusing exclusively on environmentally associated loci, we aimed to quantify the adaptive mismatch between historical and novel climatic environments rather than genome-wide neutral divergence.

To evaluate the predictive power of genomic offset, correlation analyses were performed between calculated genetic offsets and observed fitness traits, focusing on the Saclay experimental orchard, which contained the highest density of sequenced individuals (Suppl. Table 1). To increase statistical power, offset values of sequenced individuals were generalized to non-sequenced relatives and siblings at the same site, an approach justified by preliminary analyses showing that highly related individuals from the same environment exhibit near-identical offset values; where multiple siblings from the same mother were sequenced at the same site, the mean offset value was used for that family. Offset values were correlated with several fitness proxies, including summer and winter trunk diameter, ecophysiological traits (chlorophyll, flavonoids, NBI), bud break and flowering time, and absolute (AGR) and relative (RGR) growth rates across seasons (see ‘Eco-physiological trait measurement’ below).

To evaluate the capacity of geometric genomic offset (GO) to predict local adaptation, we modeled its relationship with phenotypic performance using two alternative conceptual formulations of trait mismatch relative to the local optimum (*trait_optimum_*), defined as the population mean observed at the reference site (*Y_v_*). Because theoretical models make different assumptions about how maladaptation scales with trait variation, we formally evaluated two competing metrics:

Log-transformed relative fitness ratio, 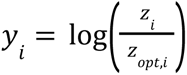 (68,69): Measures proportional relative mismatch between the observed trait (*z_i_*) and the local phenotypic optimum (*z_opt_*_,*i*_). Operating on a multiplicative scale, this formulation renders fitness sensitivity scale-invariant across baseline trait values while stabilizing variance for skewed biological metrics..

Direct quadratic mismatch, (*z_i_* – *z_opt_*_,*i*_)^2^ (10,70): Measures absolute physical mismatch between *z_i_* and *z_opt_*_,*i*_. Following traditional Gaussian selection landscapes on the raw measurement scale, this formulation tests whether absolute phenotypic deviations scale linearly or quadratically with genomic distance.

To account for repeated measures and shared ancestry, both metrics were evaluated using linear mixed-effects models (LMMs) incorporating fixed effects for linear (*g_i_*) and quadratic 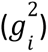 genomic offset, as well as random intercepts for measurement year 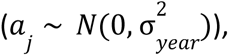 individual identity 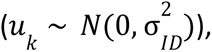 and maternal lineage 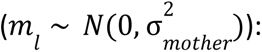

- Model A (Log-fitness ratio):

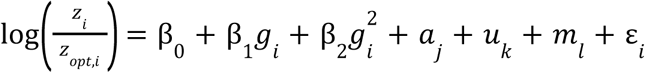
- Model B (Quadratic mismatch):

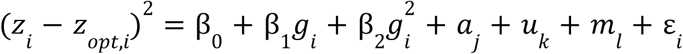

where 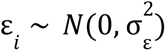 represents the residual error term.

Both mixed-effects models were fitted via Restricted Maximum Likelihood (REML) in R using lme4. Based on superior model fit (consistently lower AIC) and effective residual variance stabilization across populations, the log-linearized fitness ratio log(*W*) was selected as the optimal response variable for all downstream modeling.

To disentangle the predictive power of genomic offset (*g_i_*) from neutral demographic history (*p_i_*_,1_, *p_i_*_,2_) and broad environmental gradients 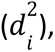, we evaluated five nested fixed-effects linear regression models predicting log-transformed fitness (*y_i_* = log(*W_i_*)):

1. 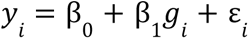
2. 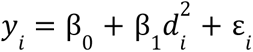
3. 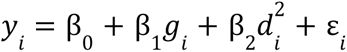
4. 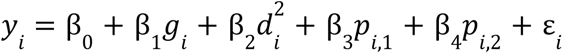
5. 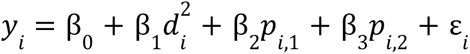

where 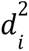 denotes squared environmental distance, and *p_i,_*_1_ and *p_i_*_,2_ represent the first two genome-wide SNP principal components controlling for population structure. Candidate specifications were compared using Akaike Information Criterion (AIC) to identify the optimal predictive model per trait.

For traits where AIC-selected models yielded unexpected positive parameter estimates ( β*_GO_* > 0) despite a biologically expected negative relationship with fitness loss (e.g., earlier, frost-sensitive Bud_Break), we performed a Leave-One-Group-Out (Jackknife) sensitivity analysis (71,72). Accessions were iteratively excluded by geographic origin cluster (e.g., Austria, Denmark, France, Italy, Romania), and all candidate models were refitted. Parameter variance, statistical significance (*p* < 0. 05), and coefficient sign stability across iterations were tracked to assess cluster leverage and potential Simpson’s paradox driven by population structure.

### Annual climate estimation, genomic offset prediction, and phenotypic validation

To capture local climatic conditions at the common garden site, daily meteorological measurements—including daily precipitation (RR), minimum temperature (TN), and maximum temperature (TX)—were obtained from the INRAE weather station at Moulon/Saclay (Station ID: 91272001). Daily records were first aggregated to monthly values by calculating total monthly precipitation along with monthly mean minimum and maximum temperatures. For each calendar year containing complete 12-month records, the 19 standard bioclimatic variables (Bio1–Bio19) were computed using the biovars function from the dismo R package (73). Incomplete years lacking full 12-month coverage were omitted from downstream analysis. To ensure baseline consistency with the CHELSA climate dataset conventions, isothermality (Bio3) was scaled by dividing standard percentage values by 100 (*Bio*3*_scaled_* = *Bio*3/100).

Using these annual bioclimatic variables, genotype–environment associations (GEA) and annual genomic offsets were calculated following the same analytical workflow implemented for the long-term (1981–2010) climate baseline. Climate-associated SNPs were identified using LFMM2 (19), and genomic offsets between each source population and the Saclay common garden were calculated for each specific measurement year using the genetic.offset function in the LEA R package (22).

To evaluate whether annual genomic offsets predicted phenotypic performance at Saclay, we modeled trait variation as a function of annual offset using linear mixed-effects models (LMMs). Because phenotypic measurements were directly paired with year-specific genomic offset estimates, random intercepts for measurement year and individual identity were omitted from this formulation. To control for shared ancestry, a random intercept for maternal lineage 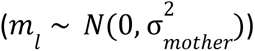 was retained:

- Model A (Log-fitness ratio):

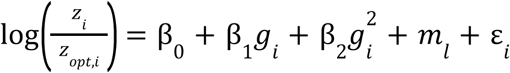
- Model B (Quadratic mismatch):

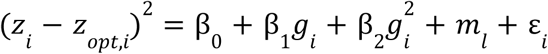

where *g_i_* and 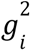 represent the linear and quadratic fixed-effect terms for annual genomic offset, *z_i_* is the observed phenotypic trait value for individual *i*, *z_opt_*_,*i*_ is the trait optimum, and ε*_i_* ∼ *N*(0, σ^2^) represents the residual error term.

### Transposable element insertion polymorphisms

To investigate the potential role of TEs in climatic adaptation, TIPs were identified across a diverse set of genomes. First, a comprehensive panTE library was constructed to capture TE diversity—including recent or line-specific TE families absent from the primary reference genome—by performing independent *de novo* annotations across seven high-quality assemblies: four newly generated *M. sylvestris* assemblies representing wild populations (Denmark, Austria, France, Romania), two previously published *M. sylvestris* assemblies (74), and the *M. domestica* reference genome (GDDH13 v1.1; (66). For each assembly, the TEdenovo pipeline (75) from the REPET v3.0 package (https://urgi.versailles.inrae.fr/Tools/REPET) was run to generate a new transposon library, followed by two rounds of curation based on TEannot (76), where any consensus not supported by at least one full-length copy in the assembly was discarded (77). Resulting annotations were merged into a single panTE library and refined using the remove-redundancy step of PASTEC (78) to retain a single representative for each transposon definition sharing ≥98% of length and ≥95% of sequence identity.

To detect structural variants (SVs) across a broader sample of genetic diversity, GraffiTE (13) was used with the GDDH13 genome as the reference genome. GraffiTE constructed a pangenome graph built from 41 long-read whole-genome assemblies, enabling the identification of non-reference SVs and mitigating single-reference bias. Long-read assemblies included the seven *M. sylvestris* and *M. domestica* genomes used in the panTE library alongside 34 additional assemblies capturing widespread *Malus* diversity. The identified SVs were annotated as TEs using the refined panTE library. The resulting graph-defined TIPs were subsequently genotyped across the short-read dataset (89 PW and WW individuals) using PanGenie (79), which leverages k-mer counts and haplotype-resolved pangenome graphs to accurately assign genotypes from short reads.

Because TIPs represent presence/absence variations of structural sequences, genotypes were converted into a 0/1/2 dosage matrix: homozygous reference (0), heterozygous (1), or homozygous alternate (2). For insertion variants (INS), a dosage of 2 indicated homozygous presence of the TE relative to the reference. For deletion variants (DEL)—where the reference genome harbors the insertion—a dosage of 2 indicated homozygous absence of the element in the query individual. SplitsTree and PCA were conducted to verify that the population structure (*K* = 5) observed with SNPs was consistent when evaluated with TIPs. Genomic offset prediction, validation, and candidate selection (using lfmm2 with p-value inflation correction and *FDR* < 0. 05) were performed following the SNP workflow described above, yielding a final filtered dataset of 80,764 TIPs across the 89 individuals.

### Orchard experimental design

Wild apple (*M. sylvestris*) seeds were collected across Europe between 2016 and 2018 (Suppl. Table 1). Seeds were stored at 5°C and sown in 2019 in a nursery at I2BC and Université Paris–Saclay, then transferred between 2021 and 2024 to a network of eight common gardens (Suppl. Fig. 16). During the nursery phase, seedlings received a preventive fungicide treatment (Captan®, 480 g/L) to protect against seedborne disease. Common gardens followed a fully randomized block design; most orchards comprised approximately 100 trees, organized into four replicates (lines) of 25 trees each, with five individuals per genetic group as defined by (17). The Saclay (France) orchard served as a larger focal site, comprising 400 trees across eight replicates. Where seed availability permitted, individuals from Italy and Spain were also included; these provenances were first established at Saclay before being introduced to the remaining network sites.

### Eco-physiological trait measurement

Trunk growth was monitored annually during winter and summer across all orchards. To account for non-circular trunk morphology, diameter was recorded as the mean of two perpendicular measurements (0° and 90°). These data were used to calculate absolute growth rate (AGR) and relative growth rate (RGR):

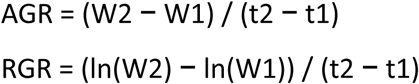

where W1 and W2 represent trunk diameter at the beginning (t1) and end (t2) of the growth period, respectively. Growth rates were analyzed across three timelines: winter-to-winter, summer-to-summer, and seasonal (summer-to-winter). Ecophysiological traits—chlorophyll (Chl), flavonoids (Flav), and the Nitrogen Balance Index (NBI)—were assessed using a Dualex^®^ leaf clip (Force-A, France), which uses fluorescence and absorbance properties to evaluate foliage physiological status. Data were collected between May and mid-July, each value representing the average of 10 leaves per tree.

Phenological traits—timing of bud break (BB) and flowering (FT)—were recorded daily from March until observation of the final tree (BB) or final flower bud (FT) in the orchard, limited to the Angers and Saclay sites for 2024 and 2025. Because flowering was significantly reduced in younger, non-grafted trees, FT data for these individuals were excluded from the final analysis.

### Genotype-by-environment variation across common gardens

To evaluate genotype-by-environment interactions, orchard datasets were standardized to approximately 100 individuals per site after filtering out mortality caused by insect or animal damage. Very young orchards (Sovata) and those heavily compromised by wildlife (Foljuif) were excluded entirely. At Saclay, which originally comprised nearly 400 trees, 100 individuals across four rows were selected via stratified random sampling to ensure representation of all genetic groups present at the site.

Genetic group assignments were based on a Discriminant Analysis of Principal Components (DAPC) performed on SSR genotyping data (17) obtained for a subset of individuals (∼30% of the trees). To assign the remaining ungenotyped individuals, we applied a hierarchical decision pipeline: unsequenced trees were assigned first based on maternal lineage (matching known genotyped siblings) or, if maternal records were unavailable, to the dominant genetic cluster observed at their respective site (Suppl. Table 10).

Genotype-by-environment variation was assessed using linear mixed-effects models fitted with lmer (80). For each trait, orchard × genetic-group interaction was used to test whether genetic groups differed in their responses among common gardens. Year, orchard, and genetic group nested within origin were treated as fixed effects, while block, mother, and individual identity were included as random effects. Model selection used AIC via a stepwise procedure (the step function); the final selected models for each trait are summarized in Suppl. Table 11. Because a significant G×E interaction does not by itself demonstrate home-site fitness advantage, these analyses are interpreted as tests of differential reaction norms rather than as direct tests of local adaptation.

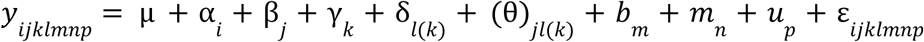

where *y_ijklmnp_* represents the phenotypic measurement recorded in year *i* within orchard *j* for individual *p* in block *m*, belonging to mother *n*, classified under genetic group *l* nested within origin *k*, with 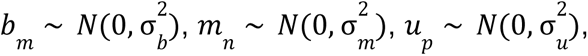 and 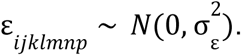.

### Genome-wide association studies and multi-locus mapping

To perform genome-wide association studies (GWAS) on individuals sequenced from the Saclay orchard, association mapping was conducted across the seven phenotypic traits measured: annual and seasonal growth rates (AGR_Annual_ss, RGR_Annual_ss, AGR_Annual_ww, RGR_Annual_ww, AGR_Seasonal, RGR_Seasonal), physiological indices (Avg_Chl, Avg_Flav, Avg_NBI), phenology (Bud_Break), and structural growth (Winter_Trunk_Measurement, and Summer_Trunk_Measurement).

Prior to association mapping, Best Linear Unbiased Predictions (BLUPs) were calculated for each trait using linear mixed models to remove environmental and block noise across measurement periods. Pairwise genetic relatedness among individuals was controlled using a genomic kinship matrix (*K*) calculated via the centered identity-by-state method on high-density SNPs, and population stratification was controlled using principal components derived from the genotypic matrix. Analyses were conducted using the Fast3VmrMLM R package (23), which integrates single-variant genome-wide scanning with expectation-maximization (EM) empirical Bayes machine learning within a compressed three-variance-component multi-locus random-effects mixed linear model.

The genotypic datasets comprised 9,875,136 SNPs and 80,764 TIPs for 72 individuals present on Saclay orchard. For the SNP dataset, GWAS was conducted at two analytical levels: (1) single-locus variant level using the standard Fast3VmrMLM module (genoType = “SNP”), and (2) linkage disequilibrium (LD)-based bin-haplotype level using the Fast3VmrMLM-Hap module (genoType = “Hap”). For the TIPs dataset, three analytical levels were evaluated: single-locus, haplotype, and molecular QTL, using the Fast3VmrMLM-mQTL module (genoType = “molecular”) to account for structural variants.

In all analyses, population structure and polygenic genetic background were controlled using integrated principal components and kinship matrices. Variants were initially selected through single-locus genome-wide scanning under a relaxed significance threshold, after which non-zero marker effects were estimated using multi-locus machine learning. Candidate quantitative trait nucleotides/loci (QTNs/QTLs) were identified using a likelihood ratio test, with a threshold of LOD ≥ 3.0 for suggested loci and a Bonferroni-corrected threshold of *P* < 0.05/*m* (where *m* is the total number of tested variants) for highly significant loci.

### Identification of candidate genes with concordant TIP and SNP climate associations

To evaluate whether climate-associated TIPs and SNPs converge on the same candidate genes, variants identified through the GEA analyses described above were further filtered and annotated. Both *lfmm2* approaches, with full = FALSE, providing a p-value for each bioclimatic variable, and with full = TRUE, providing an F-test for overall significance and a unique p-value computed across all variables, were used to detect candidate TIPs and SNPs. All detected variants were structurally annotated according to their genomic location, classified as intragenic (gene body, mRNA, ncRNA, 3’UTR, 5’UTR, CDS or exon) or intergenic; intergenic variants were further stratified by distance to the nearest gene (within 2 kb, 2–5 kb, or beyond 5 kb).

To isolate variants located within regulatory and coding regions of interest, TIPs were retained if located within the immediate flanking regulatory regions of a gene (∼2 kb upstream or downstream), while SNPs were restricted to those located within the corresponding gene body. Downstream analyses focused on genes for which a TIP in the flanking regulatory region and a SNP in the coding region were both significantly associated with the same bioclimatic variable, ensuring strict environmental and functional concordance between the two variant types (Suppl. Table 9).

To retrieve functional annotations for these candidate genes, BLAST searches were performed against the PFGD database (20) to target flowering-time-related functions in *M. sylvestris*. Gene ontology (GO) enrichment analysis was conducted on the final candidate gene list using the clusterProfiler R package and the custom ontology framework described above (67). The candidate gene list was additionally cross-referenced with the flowering-time gene annotations of (6).

Finally, to evaluate potential convergence between environmental adaptation and phenotypic variation, candidate gene-proximal TIPs identified through GEA were cross-referenced with phenotype-associated TIPs identified in the GWAS (Suppl. Table 22). A hypergeometric test was performed against the background set of genotyped TIPs to assess whether the observed variant overlap differed significantly from random expectation.

### Validation and characterization of the MdoRA.svim_asm.INS.2031 insertion across *Malus* genomes

The insertion MdoRA.svim_asm.INS.2031, located at chr2:11,288,317 in the GDDH13 reference genome, was investigated across 41 *Malus* genome assemblies (25). Two 1-kb sequences flanking the insertion site were independently mapped to each assembly using minimap2 v2.28 (-x asm5). Alignments covering ≥80% of each flank were retained. Loci were considered resolved when both flanks mapped to the same contig in a consistent orientation and order. The intervening distance was used to classify loci as *no_large_insertion*, *INS_like_183bp*, or *other_structure*.

The insertion sequence was queried against the NCBI nucleotide database using BLASTN and matched a previously characterized miniature inverted-repeat transposable element (MITE) associated with the apple *pri-miRNA172p* locus (24); GenBank KM521513.1), with 99.24% identity across 131 bp (E = 1 × 10⁻^64^). The 9-bp target site duplications (TSDs) and 18-bp terminal inverted repeats (TIRs) were delineated by comparison with Figure S8 of (24). MITE sequences recovered from 13 insertion-carrying assemblies were aligned with KM521513.1 using MAFFT v7.525 (--localpair --maxiterate 1000) to characterize nucleotide substitutions and indels.

To assess the potential of the MITE terminal inverted repeats (TIRs) to form a hairpin structure through intramolecular base pairing (24,81), the left TIR (positions 10–27) and right TIR (positions 147–164) were extracted from each MITE sequence based on their alignment to the 173-bp KM521513.1 reference. The right TIR was reverse-complemented, and both sequences were globally aligned using Biopython PairwiseAligner (match = +2, mismatch = −1, gap opening = −3, gap extension = −0.5). Complementarity was assessed by identifying canonical Watson–Crick base pairs (A–T and G–C), which can stabilize the stem of a DNA hairpin through intramolecular pairing. Because the right TIR was reverse-complemented, identical nucleotides in the alignment represented canonical pairing between the original inverted repeats. Mismatches and gap-containing columns were also counted, and complementarity was calculated as the percentage of aligned columns showing canonical pairing, including gaps in the denominator. Genomic context was characterized by positioning the mapped insertion and its flanking sequences relative to annotated genes within ±10 kb of the locus across assemblies.

### Offset and phenotypic trait comparison among crop–wild, pure-wild and wild–wild admixed individuals

To evaluate how historical cultivar admixture or natural intraspecific mixing modulates climate vulnerability and phenotypic response, we analyzed the complete wild and admixed dataset (*n* = 135), comprising pure wild (PW), wild–wild admixed (WW), and crop–wild admixed (CW) individuals. Genomic offsets relative to the common garden site were predicted following the workflow described above.

Linear mixed-effects models were fitted using the lme4 and lmerTest R packages to determine whether genetic status significantly influences the relationship between climatic mismatch and tree performance. Building upon prior model evaluations in separate PW and WW analyses—which established superior distributional stability, residual homoscedasticity, and direct percentage-based interpretability for log-transformed relative trait deviations compared to quadratic distance formulations—we focused on log-linear model specifications.

Phenotypic fitness proxies (*z_i_*), including annual and seasonal growth rates (AGR, RGR), bud break phenology, chlorophyll content, NBI, and flavonoid levels, were modeled as relative log-deviations from site-specific trait optima (*y_i_* = log(*z_i_* /*z_opt_*_,*i*_)):

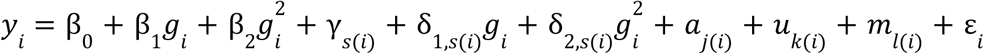

Fixed effects included linear (*g_i_*) and quadratic 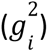 genomic offset terms, genetic status (γ*_s_*_(*i*)_ where *s* ∈ {*CW*, *PW*, *WW*}), and their respective interaction terms (δ_1,*s*(*i*)_ and δ_2,*s*(*i*)_). This parameterization allowed us to test whether crop- or wild-derived gene flow alters either the linear slope or the non-linear curvature of phenotypic responses along the genomic offset gradient. Random intercepts for measurement year 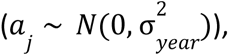 individual tree identity 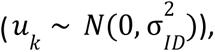 and maternal line 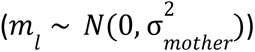 were included to account for inter-annual microclimatic variation, repeated temporal measurements, and maternal genetic structure, with residual variance 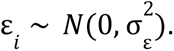.

### Past climate role in current response to climate

To evaluate whether historical climate selection shaped contemporary adaptive variation, Genotype–Environment Associations (GEA) were conducted using latent factor mixed models (LFMM2) on palaeoclimatic conditions corresponding to the Last Glacial Maximum (LGM, ∼21,000 BP). GEA was performed for precipitation of the driest month (Bio14-LGM) as well as for the full multivariate LGM bioclimatic variable set. Significant candidate SNPs were identified after False Discovery Rate correction (*q* < 0. 05).

To assess historical persistence of climate-adaptive loci, candidate SNPs identified under LGM models were intersected with those derived from contemporary global GEA models. Candidate loci were mapped to the reference genome annotation using priority-based hierarchical assignment (CDS, UTRs, exon, mRNA/gene, followed by intergenic distance tiers: < 2*kb*, 2 − 5*kb*, and > 5*kb*). Genes harboring overlapping intra-genic candidates were cross-referenced against a curated database of phenology and flowering-time genes. The statistical significance of candidate SNP enrichment between LGM and present-day GEA sets was tested using a one-tailed hypergeometric test (phyper in R) (82) against a total genome-wide background pool of *N* = 9, 875, 136 SNPs.

To test whether historical climate mismatch predicts contemporary phenotypic performance, palaeoclimatic genomic offsets were calculated following the same workflow as in contemporary analyses. In this implementation, the local reference environment was defined as the historical LGM climate at each population’s geographical origin, and the target environment was set to the present-day climate at the common garden orchard site (Saclay). Linear mixed-effects models (lme4 and lmerTest packages) were fitted to evaluate whether palaeoclimatic genomic offset predicted log-transformed relative trait deviations (*y_i_* = log(*z_i_* /*z_opt_*_,*i*_)) from site-specific optima across measured growth and phenological traits:

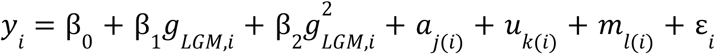

Linear (β_1_) and quadratic (β_2_) terms for palaeoclimatic genomic offset (*g_LGM_*_,*i*_) were included as fixed effects. Random intercepts for measurement year 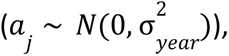 individual tree identity 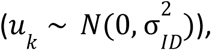 and maternal family 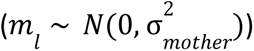 were incorporated to control for inter-annual environmental variability, repeated measurements, and maternal genetic structure, with residual error 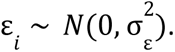

### Climate projections under future scenarios

To assess potential future maladaptation, genomic offsets were projected across three future time horizons (2011–2040, 2041–2070, 2071–2100) using the 19 CHELSA bioclimatic variables derived from three Shared Socioeconomic Pathways (SSPs) of the Coupled Model Intercomparison Project Phase 6 (CMIP6): SSP1-2.6, SSP3-7.0, and SSP5-8.5. These scenarios represent a gradient of anthropogenic radiative forcing ranging from a sustainable, low-emission pathway (SSP1-2.6; +2.6 W/m^2^) to a high-emission pathway (SSP5-8.5; +8.5 W/m^2^). Climate projections were extracted from the CHELSA database using the IPSL-CM6A-LR Earth System Model (83).

### Modeling donor–recipient translocations and climate mismatch

To model assisted migration, directional translocation vectors from candidate donor populations to target recipient orchards were identified under current and future (SSP5-8.5, 2071–2100) climate conditions. Baseline climate mismatch (Baseline_GO) was defined using the geographically nearest donor (Local_Donor), while optimal candidate donors (Optimal_Donor) were selected by minimizing residual genomic offset (Min_Offset), quantifying net adaptation gain as ΔGO=Baseline_GO−Min_Offset. Donor selection stability and 95% confidence intervals (CI95%) were evaluated using B=1,000 non-parametric bootstrap iterations, ranking candidates (Ranks 1–5) by their selection probability (Boot_Prob). Empirical permutation tests (N=10,000) assessed whether targeted donor selection significantly reduced genomic offset relative to unguided random translocation. Spatial translocation vectors were mapped in R (ggplot2, sf, ggrepel) (84–86) with vector colors indicating genomic offset. Selection uncertainty was visually encoded by displaying Rank 1 (primary) and Rank 2 (alternative) donors as solid and dashed arrows, respectively, with line width (0.6–1.4 pt), alpha transparency (α=0.40–0.95), and donor point opacity scaling continuously with Boot_Prob, alongside explicit text labels reporting individual donor IDs and selection support percentages.

### Statistical analyses and software

All statistical analyses were performed in R (v4.4.3) (82). Linear and linear mixed-effects models were fitted using the lme4 and lmerTest R packages, and model selection was based on the Akaike Information Criterion (AIC). Spatial interpolation maps were generated in R using the packages sf (84), rnaturalearth (87), gstat (88), ggpattern (89), and sp (90). Figures were produced using ggplot2 (85) and custom R scripts (https://github.com/CornilleEclecticLab/Apple-Genomic_offset).

## Abbreviations

AGR: Absolute Growth Rate
AIC: Akaike Information Criterion
CW: Crop-Wild admixed
CHELSA: Climatologies at High resolution for the Earth’s Land Surface Areas
CMIP6: Coupled Model Intercomparison Project Phase 6
CDS: Coding DNA Sequence
DAPC: Discriminant Analysis of Principal Components
db-RDA: distance-based Redundancy Analysis
DEL: Deletion
DXY: Absolute genetic divergence
FDR: False Discovery Rate
FST: Fixation index
GEA: Genome–Environment Association
GxE: Genotype-by-Environment interaction
Ho: Observed Heterozygosity
IBD: Isolation By Distance
IBE: Isolation By Environment
INS: Insertion
LD: Linkage Disequilibrium
LFMM: Latent Factor Mixed Model
LGM: Last Glacial Maximum
LMM: Linear Mixed-effects Model
MAF: Minor Allele Frequency
NBI: Nitrogen Balance Index
PanTE library: Transposable element library at the pangenomic level
PCA: Principal Component Analysis
PCNM: Principal Coordinates of Neighbour Matrices
PFGD: Plant Flowering-time Gene Database
π: Nucleotide Diversity
PW: Pure Wild
RDA: Redundancy Analysis
RGR: Relative Growth Rate
RONA: Risk Of Non-Adaptation
SNP: Single Nucleotide Polymorphism
sNMF: Sparse Non-negative Matrix Factorization
SSP: Shared Socioeconomic Pathway
TIP: Transposable element Insertion Polymorphism
WW: Wild–Wild admixed.

## Acknowledgements

This research was carried out on the High-Performance Computing resources at New York University Abu Dhabi, the Core Cluster of the Institut Français de Bioinformatique (IFB) (ANR-11-INBS-0013), and the URGI facilities (https://doi.org/10.15454/1.5572414581735654E12). We thank Adrien Falce, Olivier Langella, and Benoit Johannet for their help and support at the INRAE Génétique Quantitative et Évolution - Le Moulon lab. We thank all partners who contributed to the design, installation and monitoring of the common-garden network: Vincent Tournier, Elodie Cornille, Herve Cornille et Faustine Ducoin, and Jean-Baptiste Rey and ENRx (Espaces naturels régionaux) at Lille; the CEREEP-Ecotron Île-de-France (CNRS, ENS) at Foljuif; Unité Expérimentale Horticole, UE 0449, Centre d’Angers-Nantes, Beaucouzé, France, at Angers; Dominique Beauvais at Maison de l’Estuaire.

## Data Availability

Raw whole-genome sequencing reads for the 60 newly sequenced *M. sylvestris i*ndividuals have been deposited in NCBI SRA under BioProject [PRJNA1552270]. Sequencing and microsatellite data for previously published individuals are available in (6,17). The four new *M. sylvestris* genome assemblies are available under (25). Filtered SNP (9,751,068 and 9,875,620 SNPs) and TIP (80,764 TIPs) genotype matrices, phenotypic data from the common-garden network, and genomic offset estimates have been deposited in Zenodo 10.5281/zenodo.23665740. The panTE library is available at https://doi.org/10.57745/WFOTRJ. We obtained contemporary, Last Glacial Maximum, and future climate data from CHELSA (https://chelsa-climate.org). All scripts are available at https://github.com/CornilleEclecticLab/Apple-Genomic_offset.

## Ethics Approval and Consent to Participate

Non applicable

## Consent for Publication

All authors consent to publication.

## Author Contributions

AC conceived the study and acquired funding. AC and EM supervised the study. AC designed the common-garden network. AV, ARom, TMU, DS, ARod, TK, FD, AL, SB, and AC collected plant material and established and maintained the orchards. MC, AV, ARom, ARod, MB, FD, AL, SB, GV, AC performed phenotypic measurements. MC and AV extracted DNA and performed sequencing. MC and AM performed SNP calling and filtering. SS and JC built the panTE library, the pangenome graph, and TIP genotyping. MC, AM, SS, AC, performed population genomic and landscape genomic analyses. XC and MC performed genome–environment association and genomic offset analyses. MC and KA performed GWAS. MC performed G×E and mixed-model analyses. SS performed the MITE characterization. OF, YB contributed analytical methods (genomic offset and TE analyses). MC, SS, EM and AC wrote the manuscript with input from all co-authors. All authors read and approved the final manuscript.

## References

1. Aitken SN, Yeaman S, Holliday JA, Wang T, Curtis-McLane S. Adaptation, migration or extirpation: climate change outcomes for tree populations. Evol Appl. févr 2008;1(1):95-111. doi:10.1111/j.1752-4571.2007.00013.x

2. Duputié A, Rutschmann A, Ronce O, Chuine I. Phenological plasticity will not help all species adapt to climate change. Glob Change Biol. août 2015;21(8):3062-73. doi:10.1111/gcb.12914

3. Petit RJ, Hampe A. Some Evolutionary Consequences of Being a Tree. Annu Rev Ecol Evol Syst. 1 déc 2006;37(1):187-214. doi:10.1146/annurev.ecolsys.37.091305.110215

4. Warschefsky E, Penmetsa RV, Cook DR, Von Wettberg EJB. Back to the wilds: Tapping evolutionary adaptations for resilient crops through systematic hybridization with crop wild relatives. Am J Bot. oct 2014;101(10):1791-800. doi:10.3732/ajb.1400116

5. Brozynska M, Furtado A, Henry RJ. Genomics of crop wild relatives: expanding the gene pool for crop improvement. Plant Biotechnol J. avr 2016;14(4):1070-85. doi:10.1111/pbi.12454

6. Chen X, Dadole R, Avia K, Venon A, Brisson M, Remoué C, et al. Gene flow from the European wild apple and selection shaped the domesticated apple genome. Curr Biol. avr 2026;36(8):2104–2118.e7. doi:10.1016/j.cub.2026.03.048

7. Criado M, Brisson M, Alix K, Bourgeois YXC, François O, Marchadier É, et al. Genomic forecasting for climate-resilient fruit trees. New Phytol. août 2026;251(4):1640-53. doi:10.1111/nph.71352

8. Fitzpatrick MC, Keller SR. Ecological genomics meets community-level modelling of biodiversity: mapping the genomic landscape of current and future environmental adaptation. Vellend M, éditeur. Ecol Lett. janv 2015;18(1):1-16. doi:10.1111/ele.12376

9. Capblancq T, Fitzpatrick MC, Bay RA, Exposito-Alonso M, Keller SR. Genomic Prediction of (Mal)Adaptation Across Current and Future Climatic Landscapes. Annu Rev Ecol Evol Syst. 2 nov 2020;51(1):245-69. doi:10.1146/annurev-ecolsys-020720-042553

10. Gain C, Rhoné B, Cubry P, Salazar I, Forbes F, Vigouroux Y, et al. A Quantitative Theory for Genomic Offset Statistics. Rosenberg M, éditeur. Mol Biol Evol. 1 juin 2023;40(6):msad140. doi:10.1093/molbev/msad140

11. Rellstab C, Dauphin B, Exposito-Alonso M. Prospects and limitations of genomic offset in conservation management. Evol Appl. mai 2021;14(5):1202-12. doi:10.1111/eva.13205

12. Baduel P, Leduque B, Ignace A, Gy I, Gil J, Loudet O, et al. Genetic and environmental modulation of transposition shapes the evolutionary potential of Arabidopsis thaliana. Genome Biol. déc 2021;22(1):138. doi:10.1186/s13059-021-02348-5

13. Groza C, Chen X, Wheeler TJ, Bourque G, Goubert C. A unified framework to analyze transposable element insertion polymorphisms using graph genomes. Nat Commun. 16 oct 2024;15(1):8915. doi:10.1038/s41467-024-53294-2

14. Ma Z, Xu X, Peng W, Zhang T, Chen Z, Cao S, et al. Population genomics reveals association of transposable elements variants with climatic adaptation in wild Amur grape. Nat Commun. 26 févr 2026;17(1):3213. doi:10.1038/s41467-026-70026-w

15. Anderson JT, Song B. Plant adaptation to climate change—Where are we? J Syst Evol. sept 2020;58(5):533-45. doi:10.1111/jse.12649

16. Cornille A, Giraud T, Smulders MJM, Roldán-Ruiz I, Gladieux P. The domestication and evolutionary ecology of apples. Trends Genet. févr 2014;30(2):57-65. doi:10.1016/j.tig.2013.10.002

17. Chen X, Avia K, Forler A, Remoué C, Venon A, Rousselet A, et al. Ecological and evolutionary drivers of phenotypic and genetic variation in the European crabapple [Malus sylvestris (L.) Mill.], a wild relative of the cultivated apple. Ann Bot. 2023.

18. Brun P, Zimmermann NE, Hari C, Pellissier L, Karger DN. Global climate-related predictors at kilometer resolution for the past and future. Earth Syst Sci Data. 16 déc 2022;14(12):5573-603. doi:10.5194/essd-14-5573-2022

19. Caye K, Jumentier B, Lepeule J, François O. LFMM 2: Fast and Accurate Inference of Gene-Environment Associations in Genome-Wide Studies. Kelley J, éditeur. Mol Biol Evol. 1 avr 2019;36(4):852-60. doi:10.1093/molbev/msz008

20. Wu T, Liu Z, Yu T, Zhou R, Yang Q, Cao R, et al. Flowering genes identification, network analysis, and database construction for 837 plants. Hortic Res. 4 avr 2024;11(4):uhae013. doi:10.1093/hr/uhae013

21. Lamesch P, Berardini TZ, Li D, Swarbreck D, Wilks C, Sasidharan R, et al. The Arabidopsis Information Resource (TAIR): improved gene annotation and new tools. Nucleic Acids Res. janv 2012;40(D1):D1202-10. doi:10.1093/nar/gkr1090

22. Frichot E, François O. LEA: An R package for landscape and ecological association studies. O’Meara B, éditeur. Methods Ecol Evol. août 2015;6(8):925-9. doi:10.1111/2041-210X.12382

23. Wang J, Chen Y, Shu G, Zhao M, Zheng A, Chang X, et al. Fast3VmrMLM: A fast algorithm that integrates genome-wide scanning with machine learning to accelerate gene mining and breeding by design for polygenic traits in large-scale GWAS datasets. Plant Commun. juill 2025;6(7):101385. doi:10.1016/j.xplc.2025.101385

24. Yao J, Xu J, Cornille A, Tomes S, Karunairetnam S, Luo Z, et al. A *micro RNA* allele that emerged prior to apple domestication may underlie fruit size evolution. Plant J. oct 2015;84(2):417-27. doi:10.1111/tpj.13021

25. Mesnil A, Castric V, Lombardi G, Blein T, Yao JL, Ampomah-Dwamena C, et al. Innovation within stability: balancing selection preserves multigenic S-haplotype architec-ture in apple [Internet]. 2025 [cité 21 oct 2026]. Disponible sur: http://biorxiv.org/lookup/doi/10.1101/2025.10.28.683340 doi:10.1101/2025.10.28.683340

26. Rohde A, Bhalerao RP. Plant dormancy in the perennial context. Trends Plant Sci. mai 2007;12(5):217-23. doi:10.1016/j.tplants.2007.03.012

27. Savolainen O, Lascoux M, Merilä J. Ecological genomics of local adaptation. Nat Rev Genet. nov 2013;14(11):807-20. doi:10.1038/nrg3522

28. Aitken SN, Bemmels JB. Time to get moving: assisted gene flow of forest trees. Evol Appl. janv 2016;9(1):271-90. doi:10.1111/eva.12293

29. Holliday JA, Ritland K, Aitken SN. Widespread, ecologically relevant genetic markers developed from association mapping of climate-related traits in Sitka spruce ( *Picea sitchensis* ). New Phytol. oct 2010;188(2):501-14. doi:10.1111/j.1469-8137.2010.03380.x

30. Ma D, Li X, Guo Y, Chu J, Fang S, Yan C, et al. Cryptochrome 1 interacts with PIF4 to regulate high temperature-mediated hypocotyl elongation in response to blue light. Proc Natl Acad Sci. 5 janv 2016;113(1):224-9. doi:10.1073/pnas.1511437113

31. Cui X, Lu F, Li Y, Xue Y, Kang Y, Zhang S, et al. Ubiquitin-Specific Proteases UBP12 and UBP13 Act in Circadian Clock and Photoperiodic Flowering Regulation in Arabidopsis. Plant Physiol. 31 mai 2013;162(2):897-906. doi:10.1104/pp.112.213009

32. Liu G, Liang J, Lou L, Tian M, Zhang X, Liu L, et al. The deubiquitinases UBP12 and UBP13 integrate with the E3 ubiquitin ligase XBAT35.2 to modulate VPS23A stability in ABA signaling. Sci Adv. 8 avr 2022;8(14):eabl5765. doi:10.1126/sciadv.abl5765

33. Levy YY, Mesnage S, Mylne JS, Gendall AR, Dean C. Multiple Roles of *Arabidopsis VRN1* in Vernalization and Flowering Time Control. Science. 12 juill 2002;297(5579):243-6. doi:10.1126/science.1072147

34. Du SS, Li L, Li L, Wei X, Xu F, Xu P, et al. Photoexcited Cryptochrome2 Interacts Directly with TOE1 and TOE2 in Flowering Regulation. Plant Physiol. sept 2020;184(1):487-505. doi:10.1104/pp.20.00486

35. Yu X, Michaels SD. The Arabidopsis Paf1c Complex Component CDC73 Participates in the Modification of *FLOWERING LOCUS C* Chromatin. Plant Physiol. 1 juill 2010;153(3):1074-84. doi:10.1104/pp.110.158386

36. Duc C, Sherstnev A, Cole C, Barton GJ, Simpson GG. Transcription Termination and Chimeric RNA Formation Controlled by Arabidopsis thaliana FPA. Baulcombe DC, éditeur. PLoS Genet. 31 oct 2013;9(10):e1003867. doi:10.1371/journal.pgen.1003867

37. Kotoda N, Hayashi H, Suzuki M, Igarashi M, Hatsuyama Y, Kidou S ichiro, et al. Molecular Characterization of FLOWERING LOCUS T-Like Genes of Apple (Malus × domestica Borkh.). Plant Cell Physiol. avr 2010;51(4):561-75. doi:10.1093/pcp/pcq021

38. Heide OM, Prestrud AK. Low temperature, but not photoperiod, controls growth cessation and dormancy induction and release in apple and pear. Tree Physiol. 1 janv 2005;25(1):109-14. doi:10.1093/treephys/25.1.109

39. Wahl V, Ponnu J, Schlereth A, Arrivault S, Langenecker T, Franke A, et al. Regulation of Flowering by Trehalose-6-Phosphate Signaling in *Arabidopsis thaliana*. Science. 8 févr 2013;339(6120):704-7. doi:10.1126/science.1230406

40. Caro E, Stroud H, Greenberg MVC, Bernatavichute YV, Feng S, Groth M, et al. The SET-Domain Protein SUVR5 Mediates H3K9me2 Deposition and Silencing at Stimulus Response Genes in a DNA Methylation–Independent Manner. Grewal S, éditeur. PLoS Genet. 11 oct 2012;8(10):e1002995. doi:10.1371/journal.pgen.1002995

41. Rellstab C, Gugerli F, Eckert AJ, Hancock AM, Holderegger R. A practical guide to environmental association analysis in landscape genomics. Mol Ecol. sept 2015;24(17):4348-70. doi:10.1111/mec.13322

42. Lind BM, Menon M, Bolte CE, Faske TM, Eckert AJ. The genomics of local adaptation in trees: are we out of the woods yet? Tree Genet Genomes. avr 2018;14(2):29. doi:10.1007/s11295-017-1224-y

43. Bay RA, Harrigan RJ, Underwood VL, Gibbs HL, Smith TB, Ruegg K. Genomic signals of selection predict climate-driven population declines in a migratory bird. Science. 5 janv 2018;359(6371):83-6. doi:10.1126/science.aan4380

44. Lotterhos KE. Interpretation issues with “genomic vulnerability” arise from conceptual issues in local adaptation and maladaptation. Evol Lett. 29 mai 2024;8(3):331-9. doi:10.1093/evlett/qrae004

45. Rius M, Darling JA. How important is intraspecific genetic admixture to the success of colonising populations? Trends Ecol Evol. avr 2014;29(4):233-42. doi:10.1016/j.tree.2014.02.003

46. Todesco M, Pascual MA, Owens GL, Ostevik KL, Moyers BT, Hübner S, et al. Hybridization and extinction. Evol Appl. août 2016;9(7):892-908. doi:10.1111/eva.12367

47. Petit RJ, Aguinagalde I, De Beaulieu JL, Bittkau C, Brewer S, Cheddadi R, et al. Glacial Refugia: Hotspots But Not Melting Pots of Genetic Diversity. Science. 6 juin 2003;300(5625):1563-5. doi:10.1126/science.1083264

48. Hewitt GM. Genetic consequences of climatic oscillations in the Quaternary. Willis KJ, Bennett KD, Walker D, éditeurs. Philos Trans R Soc Lond B Biol Sci. 29 févr 2004;359(1442):183-95. doi:10.1098/rstb.2003.1388

49. Cornille A, Giraud T, Bellard C, Tellier A, Le Cam B, Smulders MJM, et al. Postglacial recolonization history of the E uropean crabapple ( *Malus sylvestris* M ill.), a wild contributor to the domesticated apple. Mol Ecol. avr 2013;22(8):2249-63. doi:10.1111/mec.12231

50. Cornille A, Feurtey A, Gélin U, Ropars J, Misvanderbrugge K, Gladieux P, et al. Anthropogenic and natural drivers of gene flow in a temperate wild fruit tree: a basis for conservation and breeding programs in apples. Evol Appl. avr 2015;8(4):373-84. doi:10.1111/eva.12250

51. Lind BM, Lotterhos KE. The accuracy of predicting maladaptation to new environments with genomic data. Mol Ecol Resour. mai 2025;25(4):e14008. doi:10.1111/1755-0998.14008

52. Ortiz-Barrientos D, Cooper M. Additive Channels in Curved Fitness Landscapes [Internet]. 2026 [cité 21 oct 2026]. Disponible sur: http://biorxiv.org/lookup/doi/10.64898/2026.03.21.713332 doi:10.64898/2026.03.21.713332

53. Auwera G van der, O’Connor BD. Genomics in the cloud: using Docker, GATK, and WDL in Terra. First edition. Sebastopol, CA: O’Reilly Media; 2020. 467 p.

54. Danecek P, Bonfield JK, Liddle J, Marshall J, Ohan V, Pollard MO, et al. Twelve years of SAMtools and BCFtools. GigaScience. 29 janv 2021;10(2):giab008. doi:10.1093/gigascience/giab008

55. Poplin R, Ruano-Rubio V, DePristo MA, Fennell TJ, Carneiro MO, Van Der Auwera GA, et al. Scaling accurate genetic variant discovery to tens of thousands of samples [Internet]. 2017 [cité 19 oct 2026]. Disponible sur: http://biorxiv.org/lookup/doi/10.1101/201178 doi:10.1101/201178

56. Chang CC, Chow CC, Tellier LC, Vattikuti S, Purcell SM, Lee JJ. Second-generation PLINK: rising to the challenge of larger and richer datasets. Gigascience. 1 déc 2015;4(1):s13742-015-0047-8. doi:10.1186/s13742-015-0047-8

57. Manichaikul A, Mychaleckyj JC, Rich SS, Daly K, Sale M, Chen WM. Robust relationship inference in genome-wide association studies. Bioinformatics. 15 nov 2010;26(22):2867-73. doi:10.1093/bioinformatics/btq559

58. Cingolani P, Platts A, Wang LL, Coon M, Nguyen T, Wang L, et al. A program for annotating and predicting the effects of single nucleotide polymorphisms, SnpEff: SNPs in the genome of Drosophila melanogaster strain w^1118^ ; iso-2; iso-3. Fly (Austin). avr 2012;6(2):80-92. doi:10.4161/fly.19695

59. Frichot E, Mathieu F, Trouillon T, Bouchard G, François O. Fast and Efficient Estimation of Individual Ancestry Coefficients. Genetics. 1 avr 2014;196(4):973-83. doi:10.1534/genetics.113.160572

60. Raj A, Stephens M, Pritchard JK. fastSTRUCTURE: Variational Inference of Population Structure in Large SNP Data Sets. Genetics. 1 juin 2014;197(2):573-89. doi:10.1534/genetics.114.164350

61. Huson DH. SplitsTree: analyzing and visualizing evolutionary data. Bioinformatics. 1 janv 1998;14(1):68-73. doi:10.1093/bioinformatics/14.1.68

62. Nei M. ESTIMATION OF AVERAGE HETEROZYGOSITY AND GENETIC DISTANCE FROM A SMALL NUMBER OF INDIVIDUALS. Genetics. 20 juill 1978;89(3):583-90. doi:10.1093/genetics/89.3.583

63. Korunes KL, Samuk K. _PIXY_ : Unbiased estimation of nucleotide diversity and divergence in the presence of missing data. Mol Ecol Resour. mai 2021;21(4):1359-68. doi:10.1111/1755-0998.13326

64. Oksanen J, Simpson GL, Blanchet FG, Kindt R, Legendre P, Minchin PR, et al. vegan: Community Ecology Package [Internet]. 2001 [cité 19 oct 2026]. p. 2.7-6. Disponible sur: https://CRAN.R-project.org/package=vegan doi:10.32614/CRAN.package.vegan

65. Karger DN, Lange S, Hari C, Reyer CPO, Conrad O, Zimmermann NE, et al. CHELSA-W5E5: daily 1 km meteorological forcing data for climate impact studies. Earth Syst Sci Data. 12 juin 2023;15(6):2445-64. doi:10.5194/essd-15-2445-2023

66. Daccord N, Celton JM, Linsmith G, Becker C, Choisne N, Schijlen E, et al. High-quality de novo assembly of the apple genome and methylome dynamics of early fruit development. Nat Genet. juill 2017;49(7):1099-106. doi:10.1038/ng.3886

67. Dadole R, Chen X, Sandoval J, Venon A, Criado M, Delpouve N, et al. Conserved transcriptional plasticity, not local adaptation, dominates early climate responses across wild and cultivated apples [Internet]. 2026 [cité 19 oct 2026]. Disponible sur: http://biorxiv.org/lookup/doi/10.64898/2026.01.15.698562 doi:10.64898/2026.01.15.698562

68. Fisher RA. The genetical theory of natural selection [Internet]. Oxford: Clarendon Press; 1930 [cité 19 oct 2026]. Disponible sur: https://www.biodiversitylibrary.org/bibliography/27468 doi:10.5962/bhl.title.27468

69. Lande R. QUANTITATIVE GENETIC ANALYSIS OF MULTIVARIATE EVOLUTION, APPLIED TO BRAIN:BODY SIZE ALLOMETRY. Evolution. mars 1979;33(1Part2):402-16. doi:10.1111/j.1558-5646.1979.tb04694.x

70. Lande R, Arnold SJ. THE MEASUREMENT OF SELECTION ON CORRELATED CHARACTERS. Evolution. nov 1983;37(6):1210-26. doi:10.1111/j.1558-5646.1983.tb00236.x

71. Quenouille MH. NOTES ON BIAS IN ESTIMATION. Biometrika. 1956;43(3-4):353-60. doi:10.1093/biomet/43.3-4.353

72. Blackman RB, Tukey JW. The measurement of power spectra from the point of view of communications engineering — Part I. Bell Syst Tech J. janv 1958;37(1):185-282. doi:10.1002/j.1538-7305.1958.tb03874.x

73. Hijmans RJ, Phillips S, Leathwick J, Elith J. dismo: Species Distribution Modeling [Internet]. 2010 [cité 23 oct 2026]. p. 1.3-16. Disponible sur: https://CRAN.R-project.org/package=dismo doi:10.32614/CRAN.package.dismo

74. Sun X, Jiao C, Schwaninger H, Chao CT, Ma Y, Duan N, et al. Phased diploid genome assemblies and pan-genomes provide insights into the genetic history of apple domestication. Nat Genet. déc 2020;52(12):1423-32. doi:10.1038/s41588-020-00723-9

75. Flutre T, Duprat E, Feuillet C, Quesneville H. Considering Transposable Element Diversification in De Novo Annotation Approaches. PLOS ONE. 31 janv 2011;6(1):e16526. doi:10.1371/journal.pone.0016526

76. Quesneville H, Bergman CM, Andrieu O, Autard D, Nouaud D, Ashburner M, et al. Combined evidence annotation of transposable elements in genome sequences. PLoS Comput Biol. juill 2005;1(2):166-75. doi:10.1371/journal.pcbi.0010022 PubMed PMID: 16110336; PubMed Central PMCID: PMC1185648.

77. Jamilloux V, Daron J, Choulet F, Quesneville H. De novo annotation of transposable elements : tackling the fat genome issue. Proc IEEE. 2017;105(3):474. doi:10.1109/JPROC.2016.2590833

78. Hoede C, Arnoux S, Moisset M, Chaumier T, Inizan O, Jamilloux V, et al. PASTEC: an automatic transposable element classification tool. PloS One. 2014;9(5):e91929. doi:10.1371/journal.pone.0091929 PubMed PMID: 24786468; PubMed Central PMCID: PMC4008368.

79. Ebler J, Ebert P, Clarke WE, Rausch T, Audano PA, Houwaart T, et al. Pangenome-based genome inference allows efficient and accurate genotyping across a wide spectrum of variant classes. Nat Genet. avr 2022;54(4):518-25. doi:10.1038/s41588-022-01043-w

80. Bates D, Maechler M, Bolker B, Walker S, Ly A, Jagan M. lme4: Linear Mixed-Effects Models using « Eigen » and S4 [Internet]. 2003 [cité 19 oct 2026]. p. 2.0-6. Disponible sur: https://CRAN.R-project.org/package=lme4 doi:10.32614/CRAN.package.lme4

81. Piriyapongsa J, Jordan IK. Dual coding of siRNAs and miRNAs by plant transposable elements. RNA. mai 2008;14(5):814-21. doi:10.1261/rna.916708 PubMed PMID: 18367716; PubMed Central PMCID: PMC2327354.

82. R Core Team. R: A Language and Environment for Statistical Computing [Internet]. Vienna, Austria: R Foundation for Statistical Computing; 2025. Disponible sur: https://www.R-project.org/

83. Boucher O, Servonnat J, Albright AL, Aumont O, Balkanski Y, Bastrikov V, et al. Presentation and Evaluation of the IPSL-CM6A-LR Climate Model. J Adv Model Earth Syst. juill 2020;12(7):e2019MS002010. doi:10.1029/2019MS002010

84. Pebesma E. sf: Simple Features for R [Internet]. 2016 [cité 19 oct 2026]. p. 1.1-3. Disponible sur: https://CRAN.R-project.org/package=sf doi:10.32614/CRAN.package.sf

85. Wickham H, Chang W, Henry L, Pedersen TL, Takahashi K, Wilke C, et al. ggplot2: Create Elegant Data Visualisations Using the Grammar of Graphics [Internet]. 2007 [cité 6 oct 2026]. p. 4.0.3. Disponible sur: https://CRAN.R-project.org/package=ggplot2 doi:10.32614/CRAN.package.ggplot2

86. Slowikowski K, Brand T van den, Schep A, Hughes S, Dang TK, Lukauskas S, et al. ggrepel: Automatically Position Non-Overlapping Text Labels with « ggplot2 » [Internet]. 2026 [cité 21 oct 2026]. Disponible sur: https://cran.r-project.org/web/packages/ggrepel/index.html

87. Massicotte P, South A. rnaturalearth: World Map Data from Natural Earth [Internet]. 2017 [cité 19 oct 2026]. p. 1.2.0. Disponible sur: https://CRAN.R-project.org/package=rnaturalearth doi:10.32614/CRAN.package.rnaturalearth

88. Pebesma E, Graeler B. gstat: Spatial and Spatio-Temporal Geostatistical Modelling, Prediction and Simulation [Internet]. 2003 [cité 19 oct 2026]. p. 2.1-6. Disponible sur: https://CRAN.R-project.org/package=gstat doi:10.32614/CRAN.package.gstat

89. Fc M, Davis TL, ggplot2 authors. ggpattern: « ggplot2 » Pattern Geoms [Internet]. 2022 [cité 19 oct 2026]. p. 1.3.1. Disponible sur: https://CRAN.R-project.org/package=ggpattern doi:10.32614/CRAN.package.ggpattern

90. Pebesma E, Bivand R. sp: Classes and Methods for Spatial Data [Internet]. 2005 [cité 21 oct 2026]. p. 2.2-4. Disponible sur: https://CRAN.R-project.org/package=sp doi:10.32614/CRAN.package.sp

