## Supplementary figures for "Genomic forecasting links climate-associated variation to spring phenology in the European wild apple, a crop wild relative of the cultivated apple": Supplementary_Figures.docx

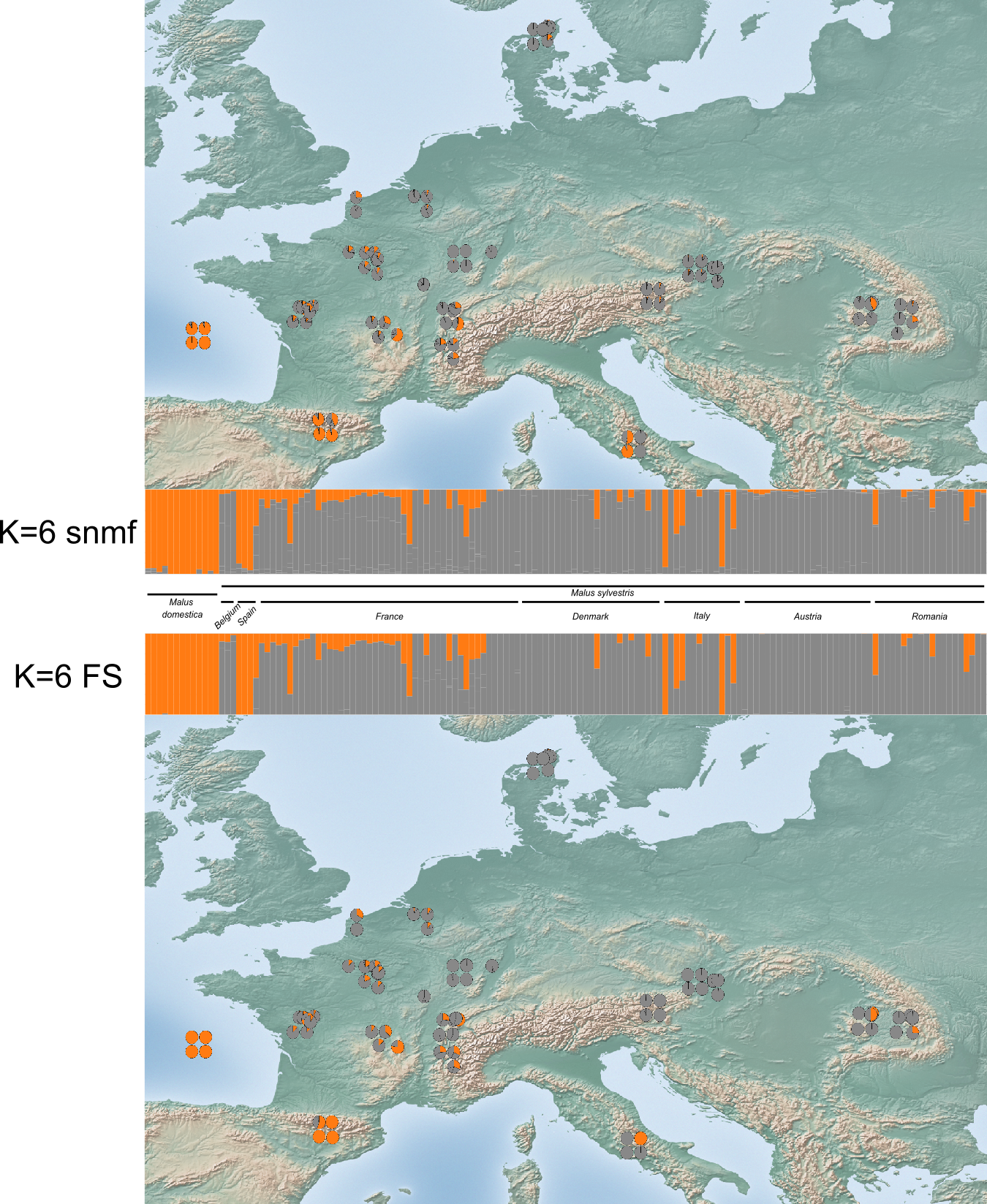

**Suppl. Fig. 1: Detection of crop introgression in wild populations using 13 cultivated reference individuals and population structure analyses performed with sNMF and fastSTRUCTURE at K = 6 (see Suppl. Fig. 19).** Cultivated individuals and membership in the P.Dom cluster are shown in orange, whereas membership in all other clusters is shown in grey. We classified individuals with a P.Dom membership coefficient ≥ 10% as crop–wild admixed.

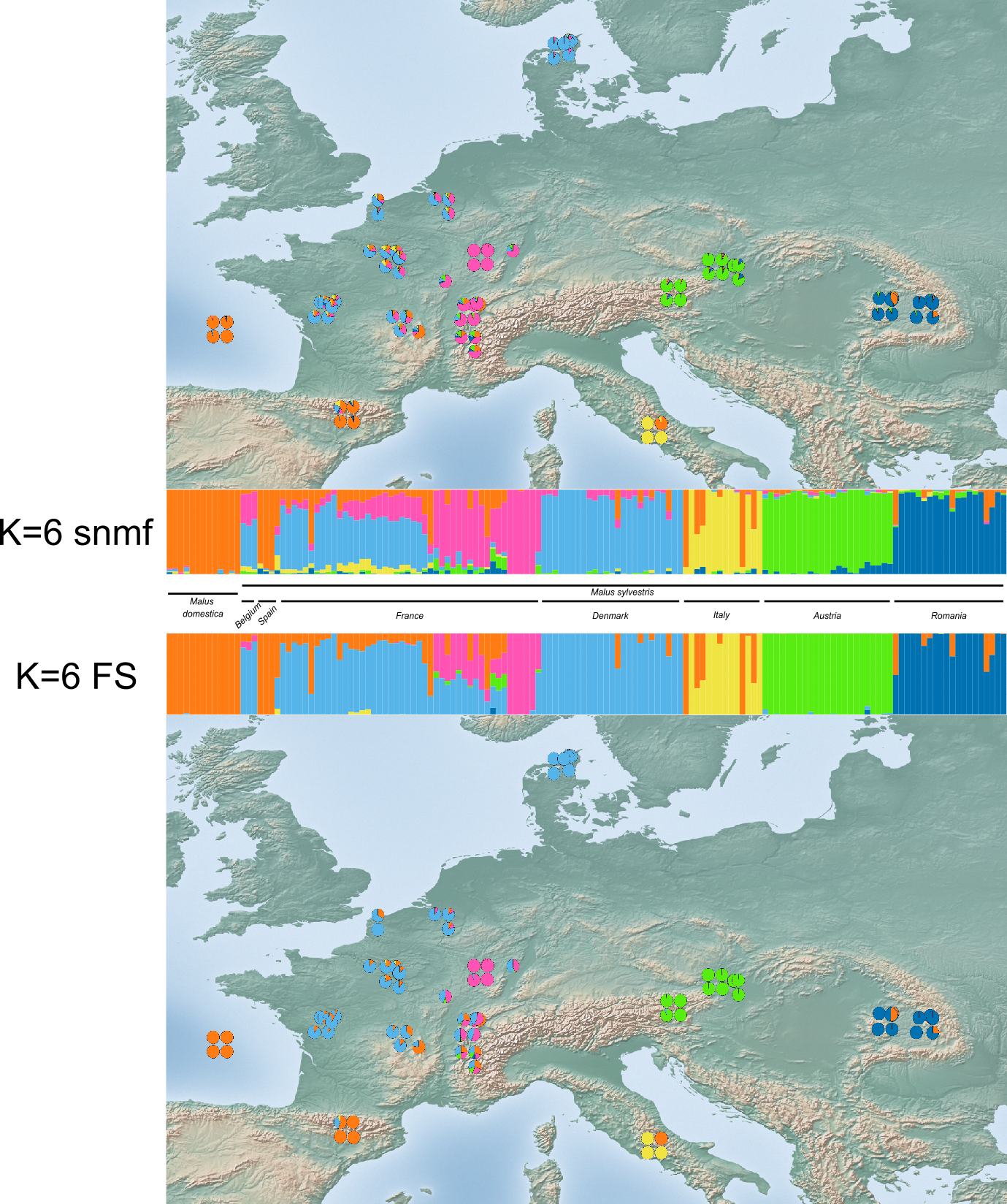

**Suppl. Fig. 2: Population structure inferred using sNMF (K = 6) and fastSTRUCTURE (K = 6, MajorCluster).** The map displays pie charts representing individual ancestry coefficients for each of the 148 sampled individuals, with colors indicating cluster membership. The corresponding ancestry bar plots are shown alongside. Six genetic clusters were identified: P.Dom (orange), P.West_Eur (cyan), P.Lor (pink), P.Ita (yellow), P.Aus (green), and P.Rom (blue).

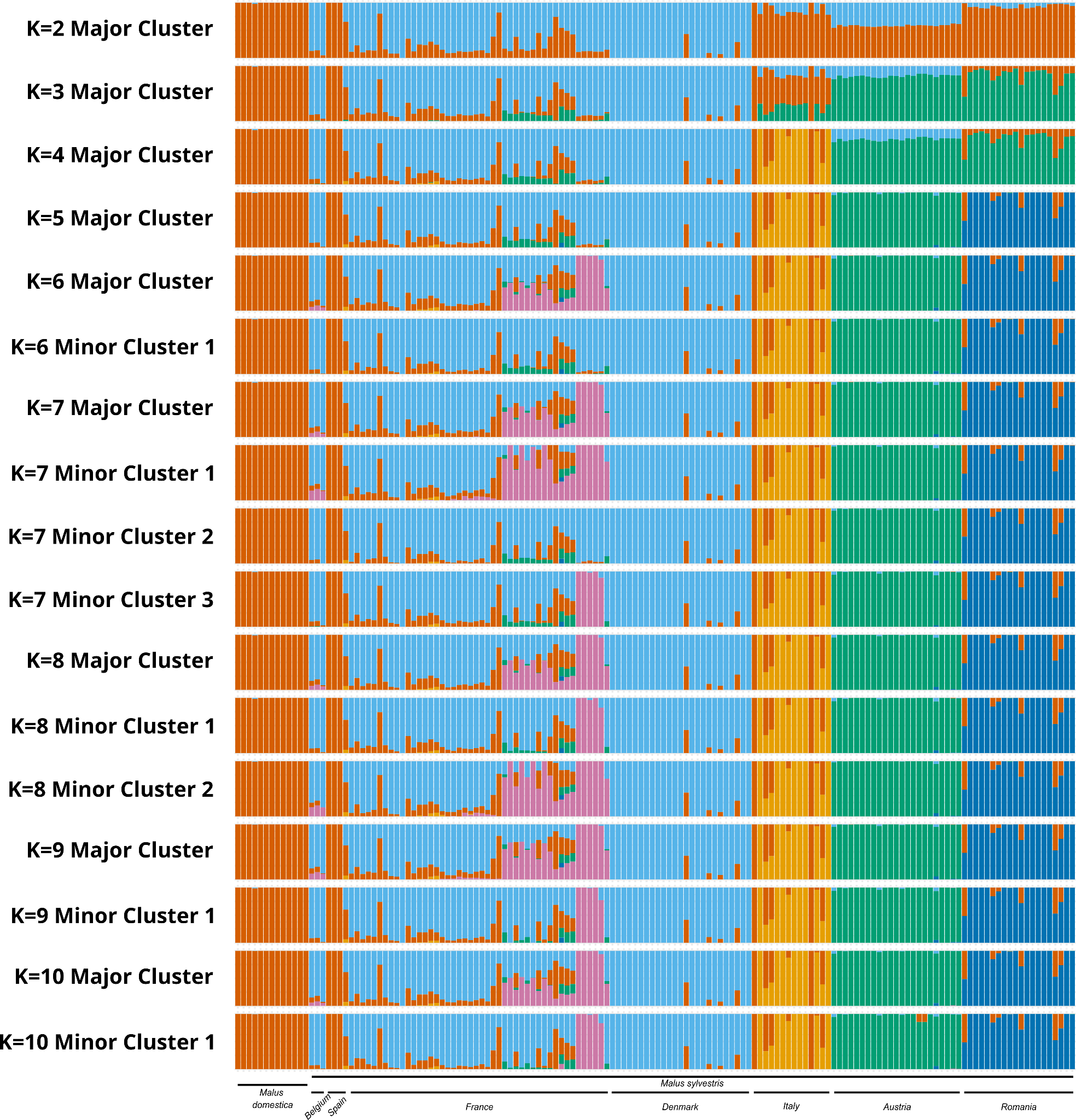

**Suppl. Fig. 3: Population structure and individual ancestry proportions from K=2 to K=10.** Individual ancestry barplots generated using fastSTRUCTURE (20 repetitions per $K$) across genetic cluster numbers ranging from $K=2$ to $K=10$. The dataset comprises 148 individuals ($13$ *Malus domestica* and $135$ *Malus sylvestris*) genotyped at 54,092 unlinked, synonymous SNPs (see Suppl. Fig. 19). Each vertical bar represents a single individual, with color segments corresponding to the proportional membership coefficient in each inferred ancestral cluster.

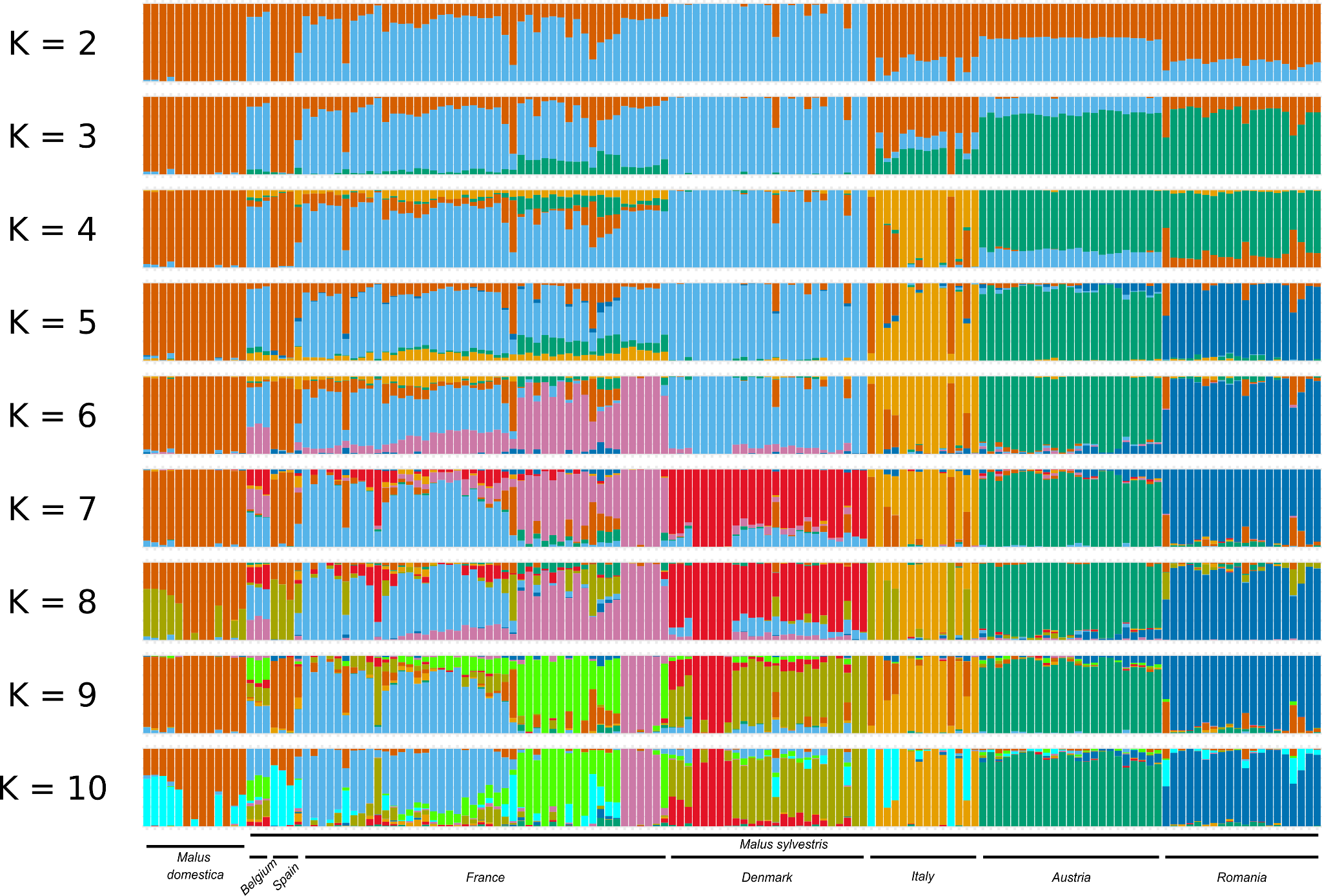

**Suppl. Fig. 4: Population structure and individual ancestry proportions from K=2 to K=10.** Individual ancestry barplots generated using sNMF (20 repetitions per $K$) across genetic cluster numbers ranging from $K=2$ to $K=10$. The dataset comprises 148 individuals ($13$ *Malus domestica* and $135$ *Malus sylvestris*) genotyped at 54,092 unlinked, synonymous SNPs (see Suppl. Fig. 19). Each vertical bar represents a single individual, with color segments corresponding to the proportional membership coefficient in each inferred ancestral cluster.

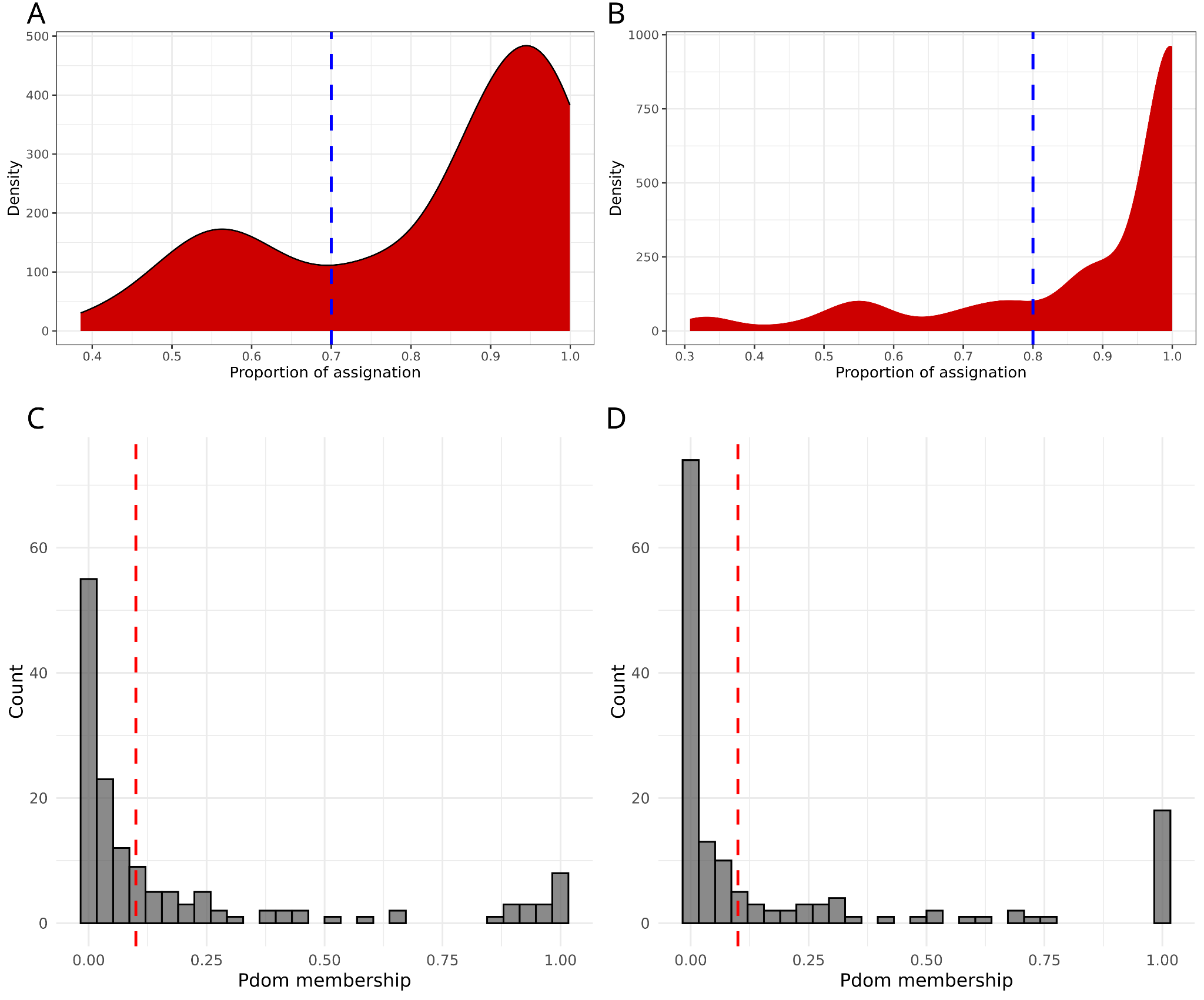

**Suppl. Fig. 5**: **Max membership coefficient at K=6 (n=148; A & B) and membership distribution (C & D) of the Pdom (*M. domestica)* cluster with sNMF (A & C) and fastSTRUCTURE (B & D).** The blue dotted line marks the threshold used at K=6 (0.70 for fastSTRUCTURE and 0.80 for sNMF) to assign each individual to population groups, and the red dotted line marks the 10% threshold used to separate pure wild (PW) from crop-wild (CW) individuals.

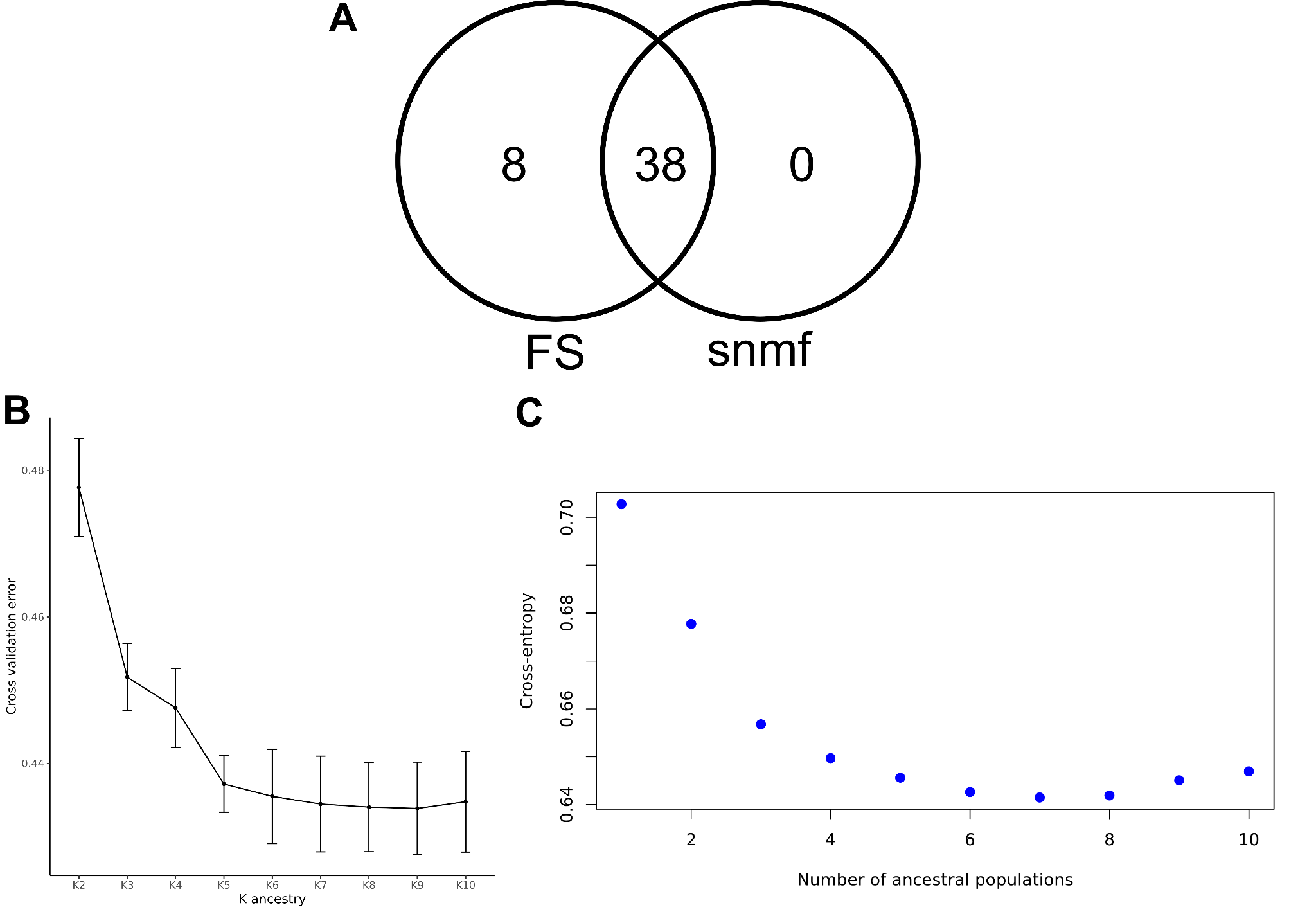

**Suppl. Fig. 6: Comparison of crop–wild admixture detection and model evaluation metrics.** **(A)** Venn diagram showing shared and method-specific crop–wild (CW) admixed individuals identified by fastSTRUCTURE (FS) and sNMF at K = 6. **(B)** Cross-validation results for fastSTRUCTURE tested for K=2 to K=10, with 20 repetitions. **(C)** Cross-entropy results for SNMF tested for K=2 to K=10, with 20 repetitions.

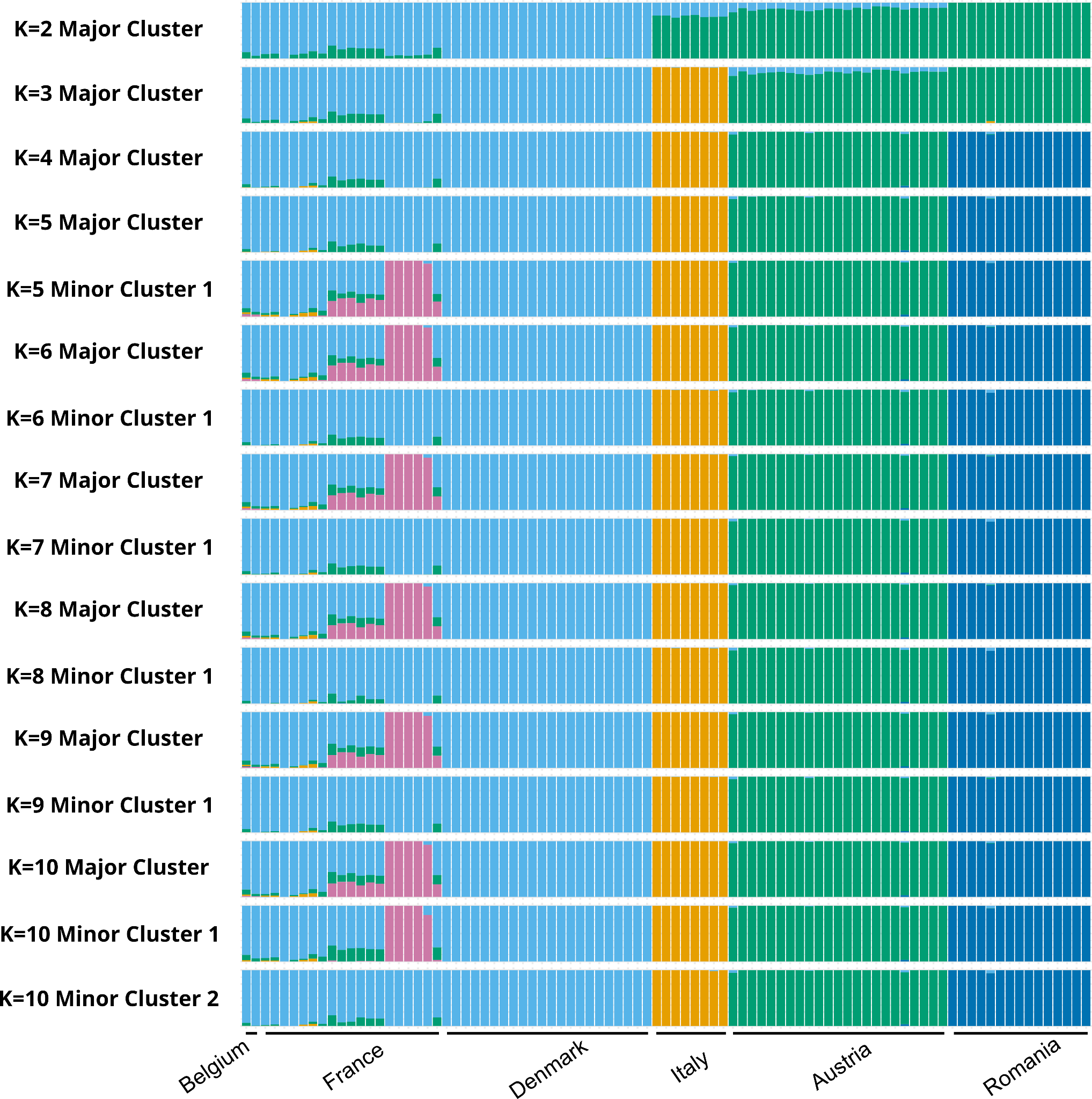

**Suppl. Fig. 7: Population structure and individual ancestry proportions from K=2 to K=10.** Individual ancestry barplots generated using fastSTRUCTURE (20 repetitions per $K$) across genetic cluster numbers ranging from $K=2$ to $K=10$. The dataset comprises 89 individuals (filtered to exclude crop-wild admixed individuals) genotyped at 54,050 unlinked, synonymous SNPs (see Suppl. Fig. 19). Each vertical bar represents a single individual, with color segments corresponding to the proportional membership coefficient in each inferred ancestral cluster.

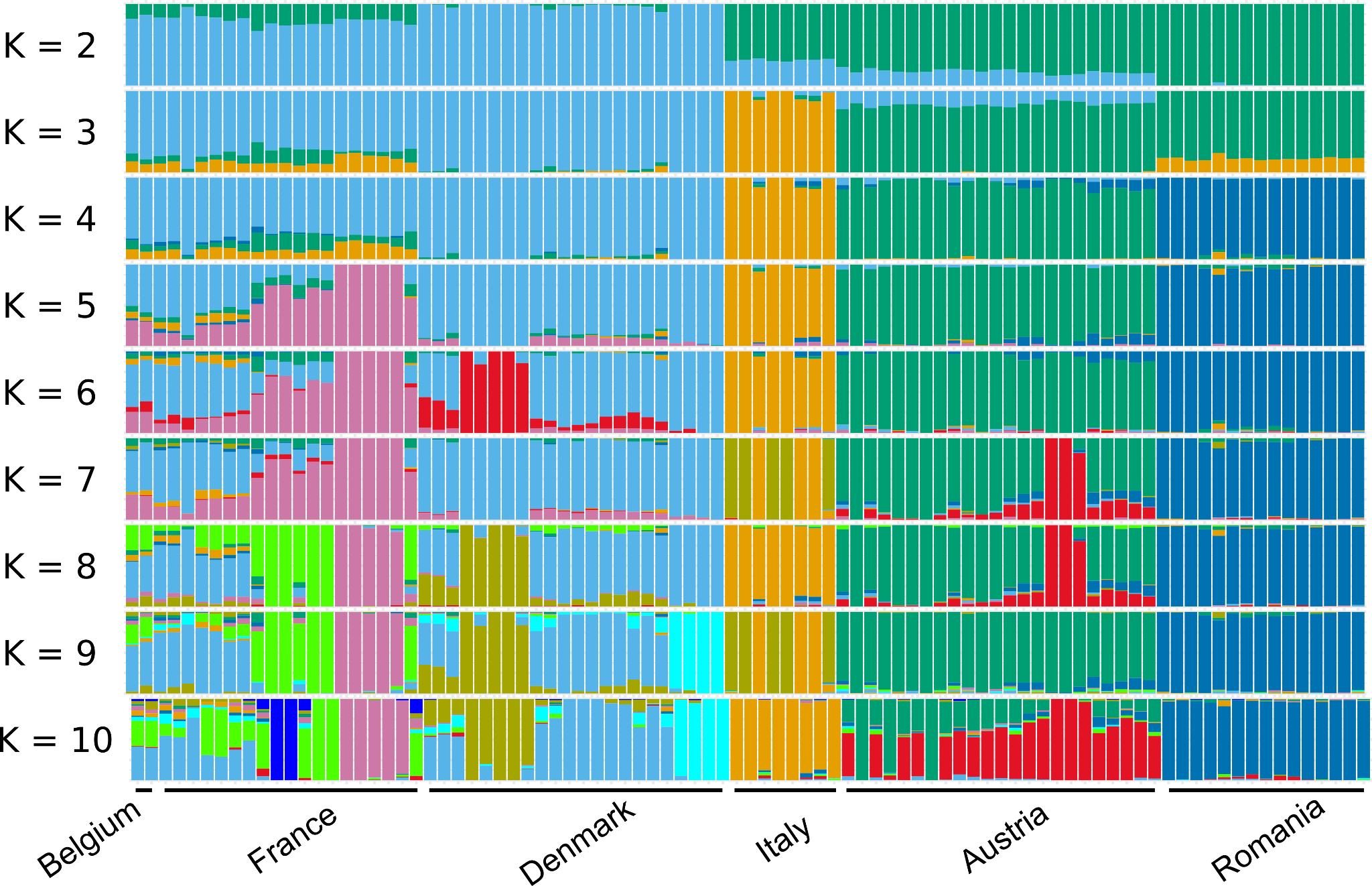

**Suppl. Fig. 8: Population structure and individual ancestry proportions from K=2 to K=10.** Individual ancestry barplots generated using sNMF (20 repetitions per $K$) across genetic cluster numbers ranging from $K=2$ to $K=10$. The dataset comprises 89 individuals (filtered to exclude crop-wild admixed individuals) genotyped at 54,050 unlinked, synonymous SNPs (see Suppl. Fig. 19). Each vertical bar represents a single individual, with color segments corresponding to the proportional membership coefficient in each inferred ancestral cluster.

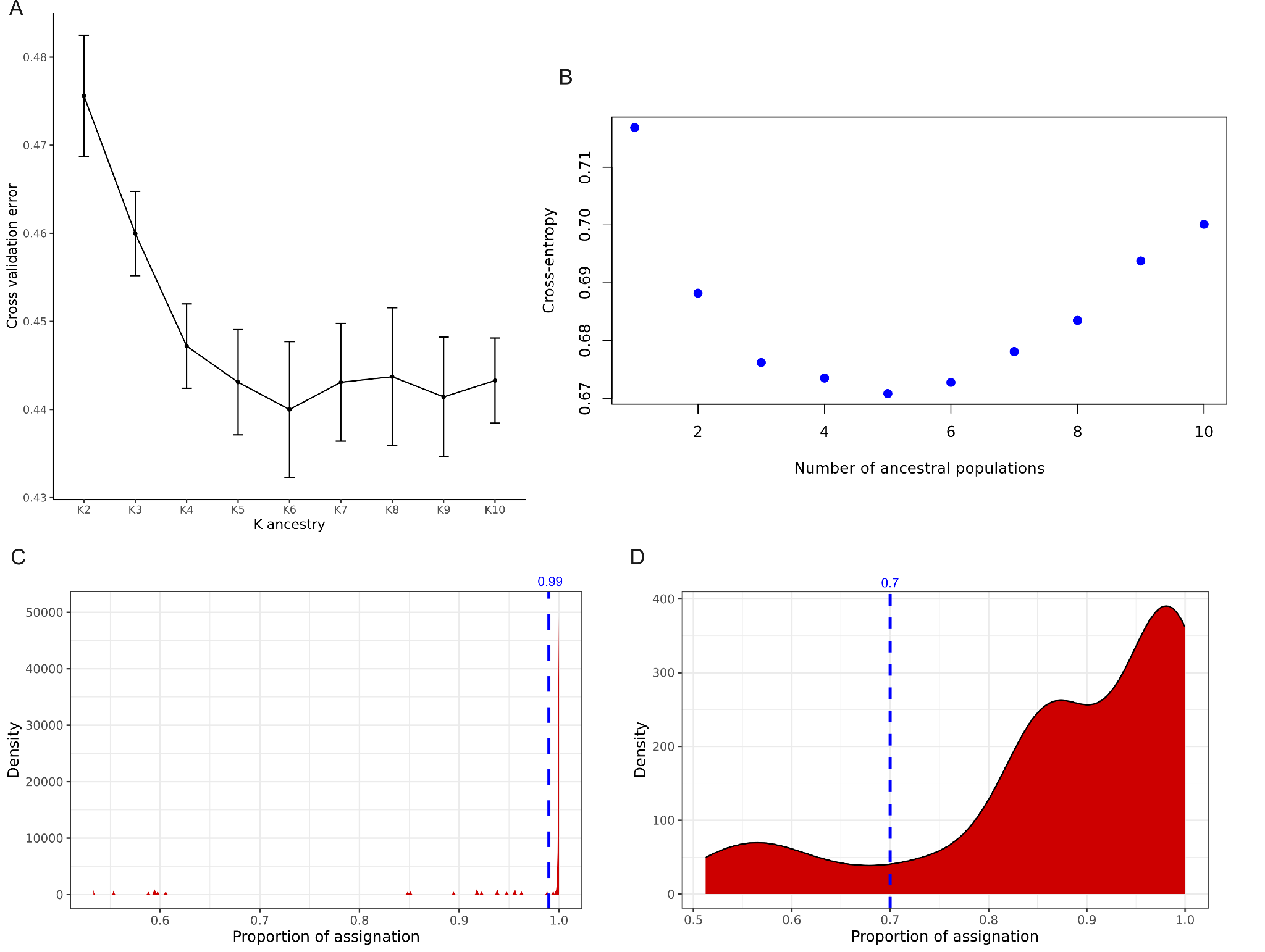

**Suppl. Fig. 9**: **Model evaluation metrics and maximum ancestry membership distributions for wild individuals (*n* = 89, excluding crop–wild hybrids).** (A) Cross-validation metrics for fastSTRUCTURE tested across *K* = 2 to *K* = 10 (20 replicates per *K*). (B) Cross-entropy scores for sNMF tested across *K* = 2 to *K* = 10 (20 replicates per *K*). (C) Distribution of maximum ancestry membership coefficients from fastSTRUCTURE at *K* = 5 (MinorCluster criterion) applying an ancestry threshold of 0.99. (D) Distribution of maximum ancestry membership coefficients from sNMF at *K* = 5 applying an ancestry threshold of 0.70.

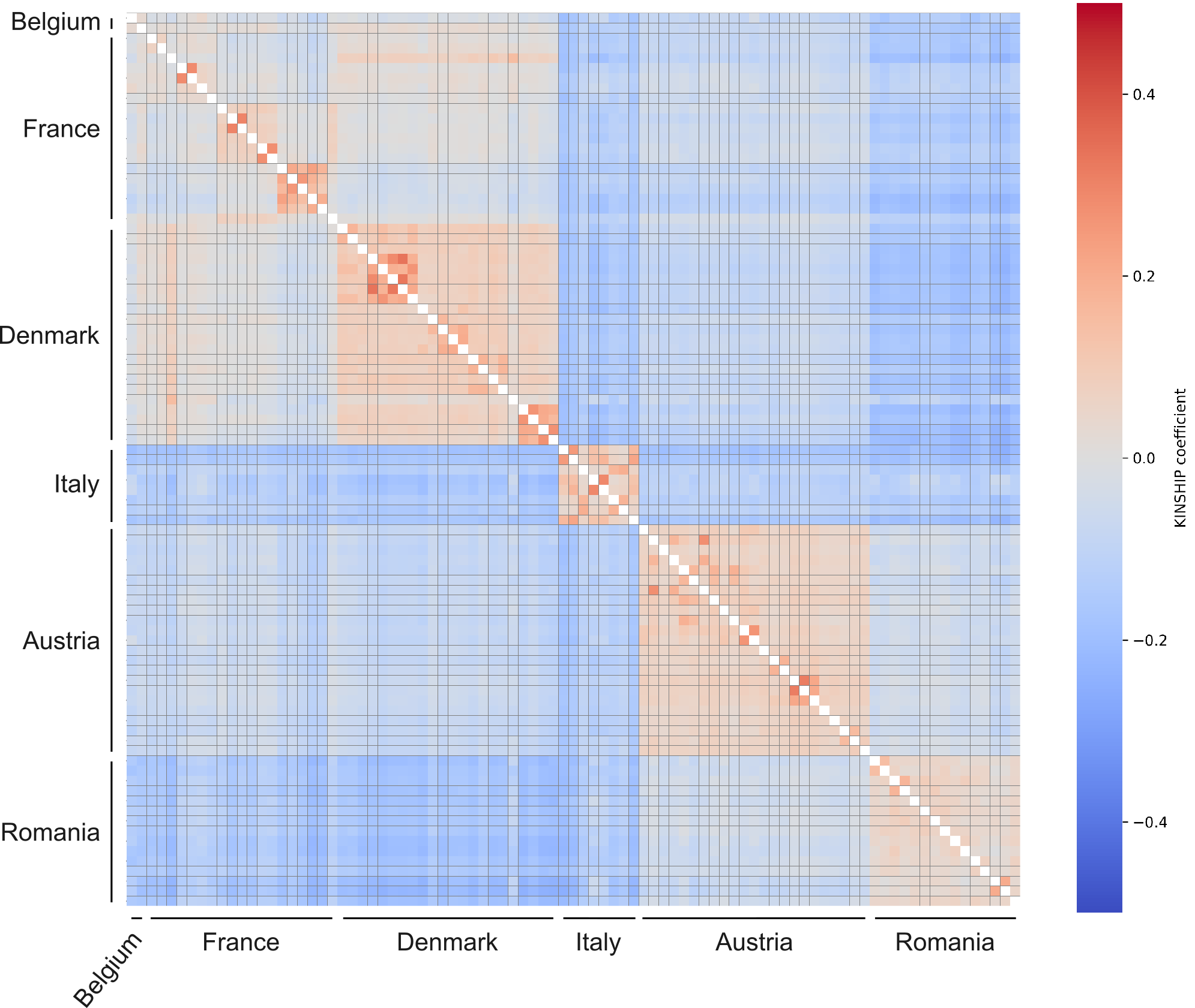

**Suppl. Fig. 10: Heatmap of pairwise kinship coefficients of pure wild and wild–wild admixed individuals.** Pairwise kinship coefficients ($\phi$) among the 89 PW and WW filtered individuals, illustrating patterns of relatedness grouped by species and country of origin (rather than inferred genetic clusters; see Fig. 2 for cluster assignments). Kinship was estimated using the relationship inference algorithm (KING-robust) implemented in PLINK 2 (1) based on the same unlinked, synonymous SNP dataset used for population structure analysis (54,050 SNPs, $n=89$; see Suppl. Fig. 19). Pairwise relationship classes follow standard KING kinship thresholds: duplicate or monozygotic twin pairs ($\phi>0.354$), first-degree relatives ($0.177<\phi\leq0.354$), second-degree relatives ($0.0884<\phi\leq0.177$), third-degree relatives ($0.0442<\phi\leq0.0884$), and unrelated pairs ($\phi<0.0442$).

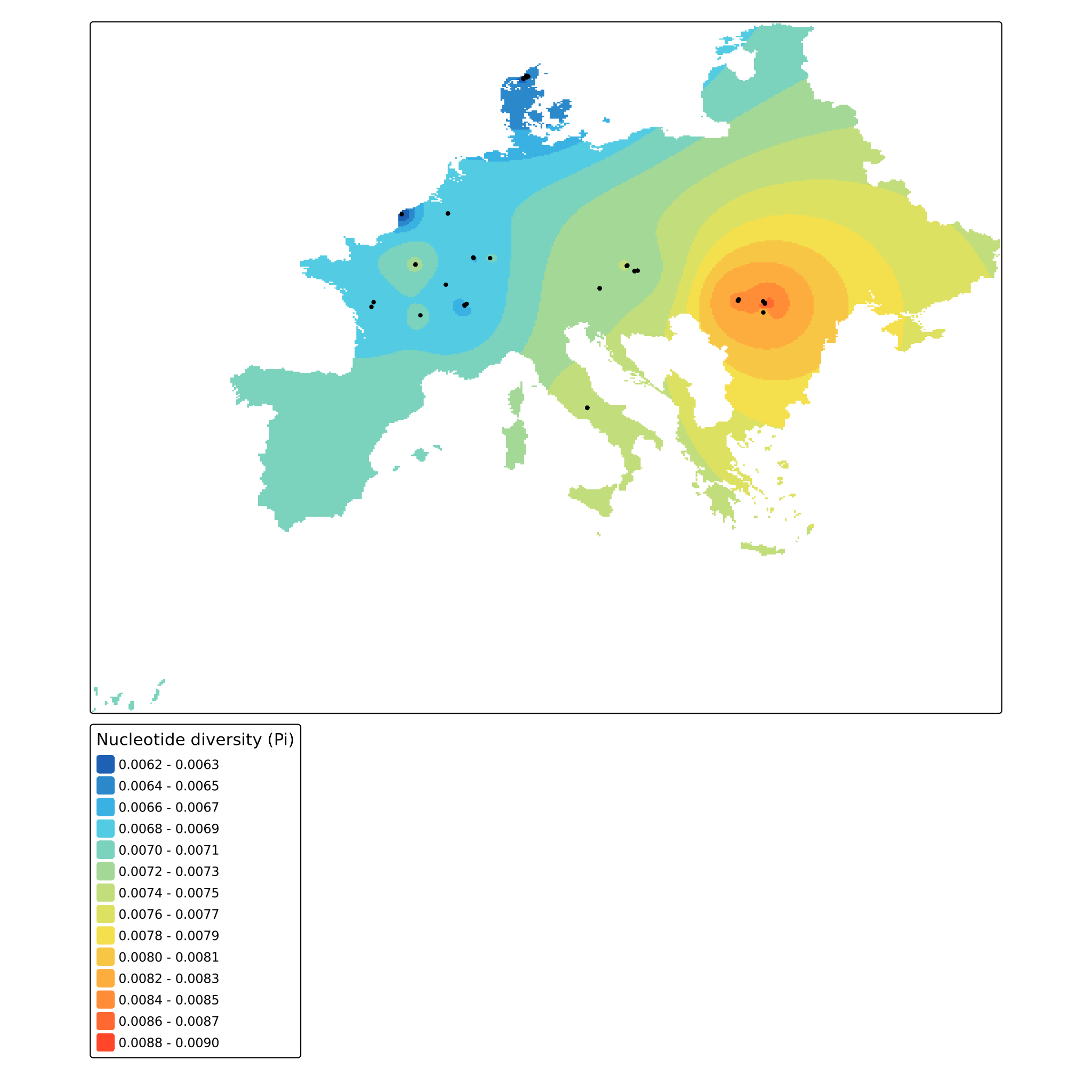

**Suppl. Fig. 11: Individual-level nucleotide diversity (**$\pi$**).** Distribution of individual-level nucleotide diversity ($\pi$) calculated using pixy across 89 wild individuals (excluding crop–wild hybrids). Estimates were computed from a filtered genome-wide dataset of 9,751,068 SNPs ($F_{missing}<0.20$, $MAF>0.05$).

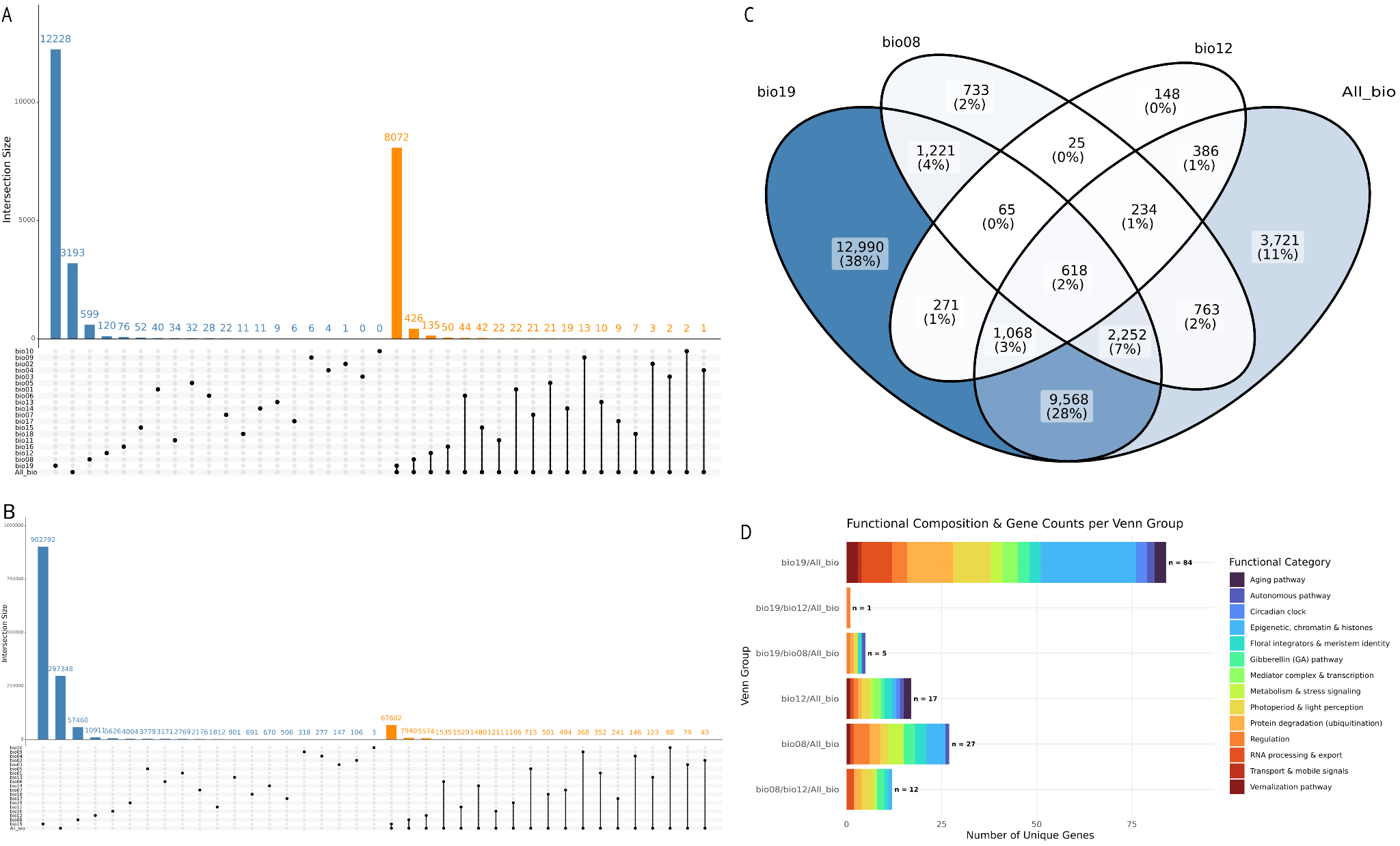

**Suppl. Fig. 12: Genomic distribution and functional profiling of candidate variants and genes identified via GEA (89 samples, 1,451,380 SNPs).** **(A)** UpSet plot displaying set intersections at the gene level across individual environmental variables and global datasets. **(B)** UpSet plot displaying set intersections at the SNP level across environmental variables. **(C)** Four-way Venn diagram highlighting overlapping gene sets between the three top bioclimatic variables (*bio19*, *bio08*, *bio12*) and all environmental variables (*All_bio*). **(D)** Distribution and composition of flowering time–related functional categories across the identified Venn intersection groups.

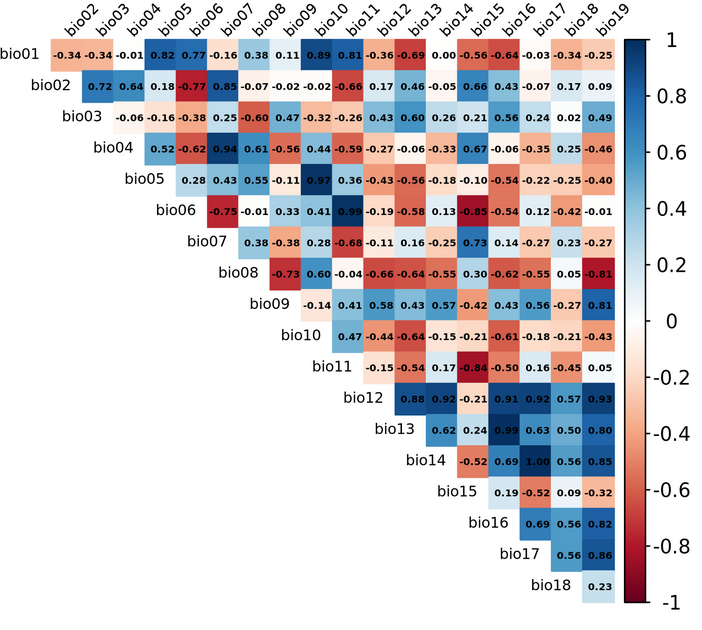

**Suppl. Fig. 13: Correlation matrix of bioclimatic variables.** Pairwise Pearson correlation matrix of the 19 bioclimatic variables extracted from the CHELSA climate database (1981–2010 baseline period, 30 arc-second resolution).

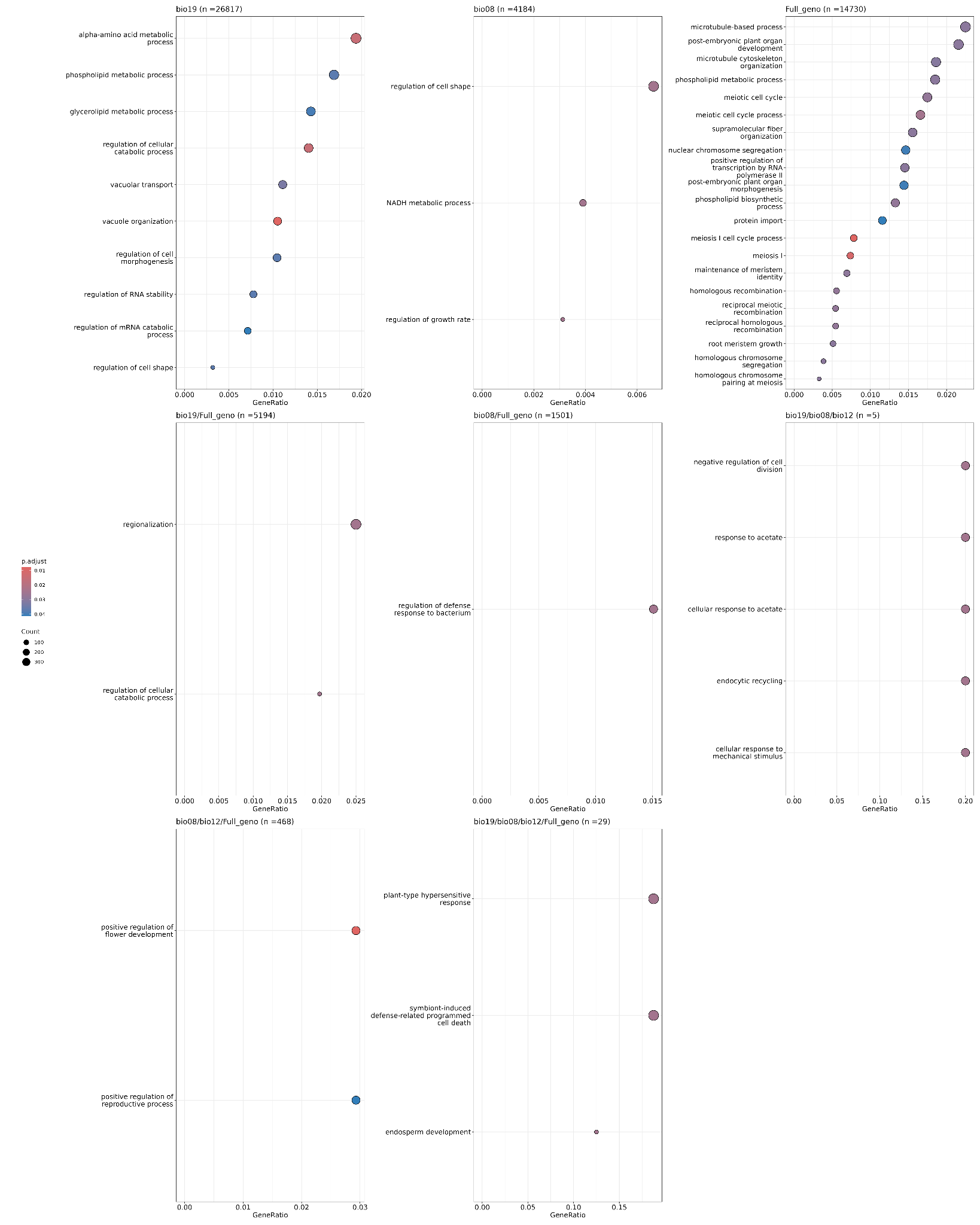
**Suppl. Fig. 14: Gene Ontology (GO) enrichment analysis across bioclimatic candidate gene sets.** Dot plots displaying significantly enriched Biological Process (BP) GO terms across distinct candidate gene sets, including bioclimatic-variable-specific sets (Bio8, Bio12, Bio19), the multi-variable model (All_Bio), and their overlapping intersections (see Suppl. Fig. 12). Enrichment analysis was performed using the clusterProfiler package against a background universe comprising all annotated genes from the GDDH13 reference genome, using functional annotations from (2). $P$-values were adjusted for multiple testing using the False Discovery Rate (FDR) method, with significantly overrepresented terms selected at adjusted $p<0.05$ and $q<0.05$ thresholds. Up to the top 50 enriched terms are shown per panel, with $n$ denoting the number of candidate genes in each set.

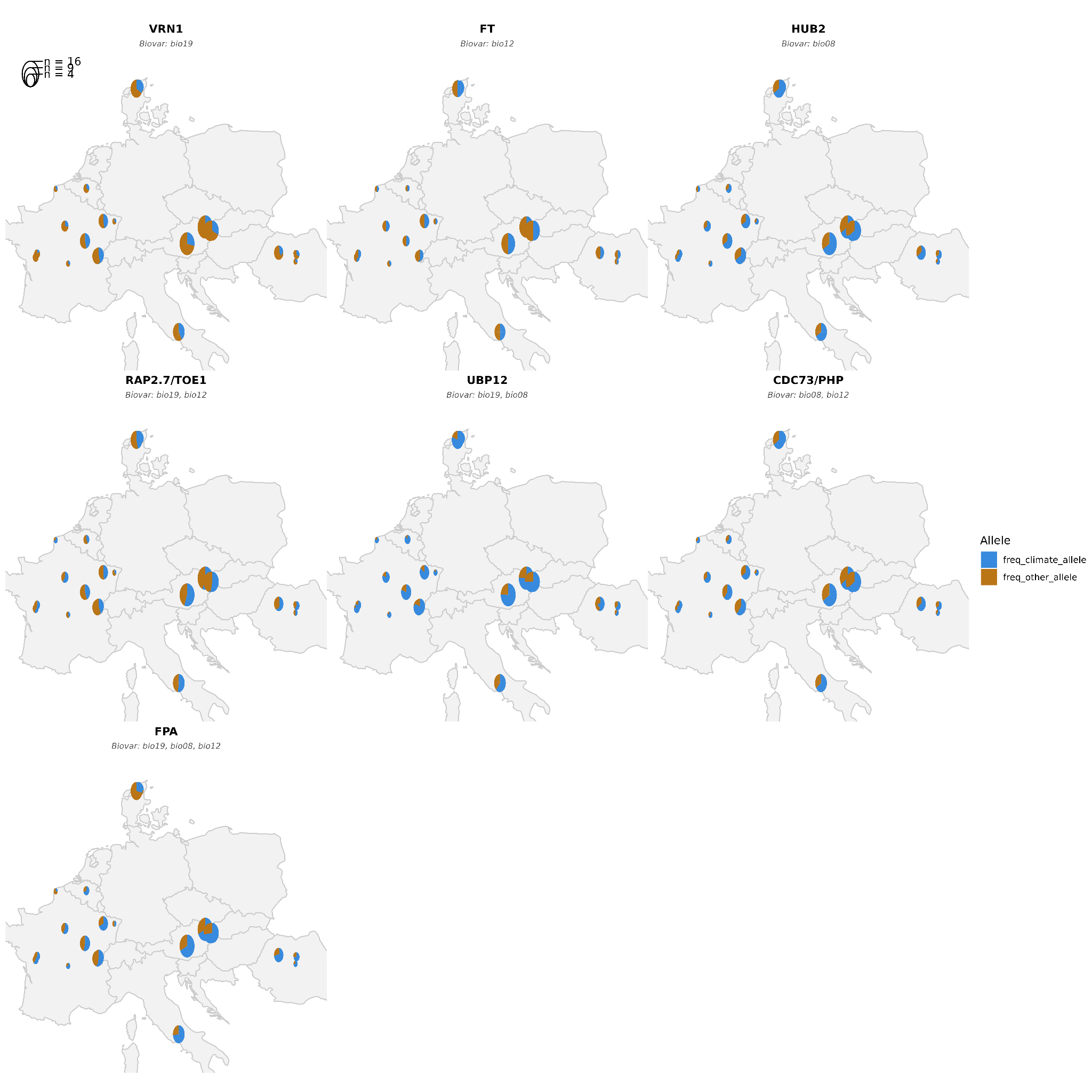

**Suppl. Fig. 15: Spatial distribution of climate-associated allele frequencies for top flowering-time candidate genes in European *Malus* populations.** Subpanels depict representative non-redundant candidate genes containing the highest density of GEA-identified SNPs associated with individual or combined bioclimatic variables (bio08: Mean temperature of wettest quarter; bio12: Annual precipitation; bio19: Precipitation of coldest quarter; Suppl. Table 9). Subtitles indicate the specific bioclimatic variable(s) associated with candidate loci within each gene region. Pie charts at each sampling location show the site-level mean frequency of the climate-associated allele (blue) relative to alternative alleles (brown). Pie chart area scales with sampling effort ($n$ individuals per site).

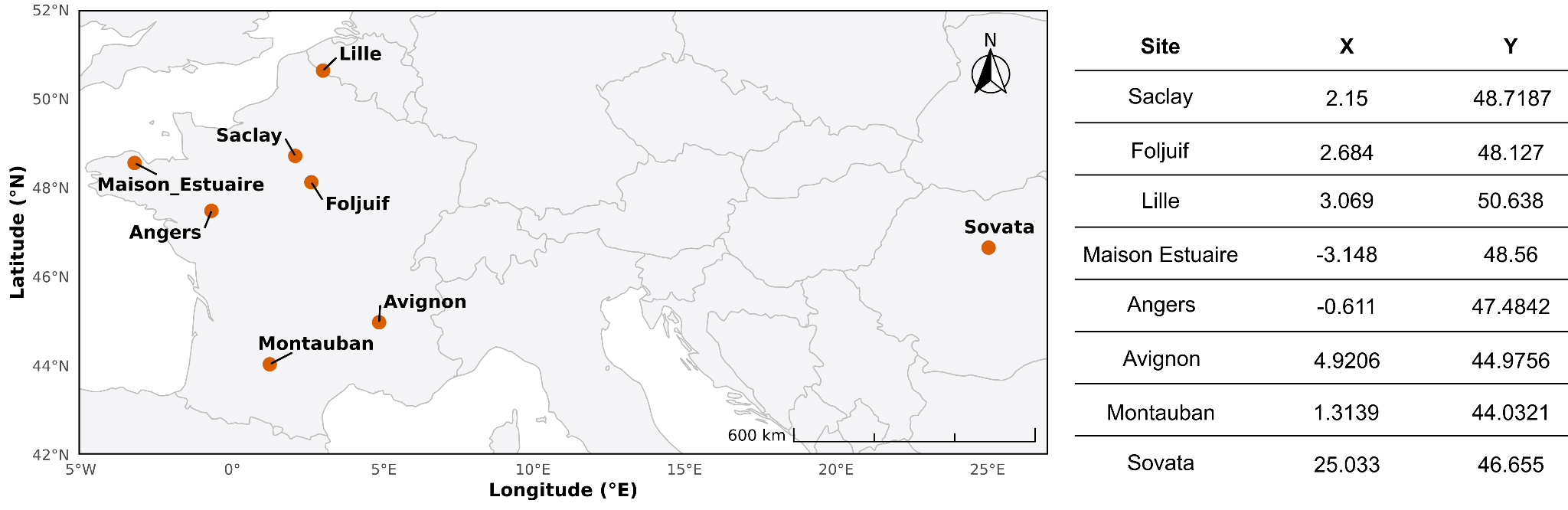

**Suppl. Fig. 16: Geographic distribution of the eight common garden orchards.** Map showing the locations of the experimental common garden sites established across Europe between 2021 and 2024 for phenotypic monitoring of *Malus sylvestris*: Saclay (France, n = 400 trees), Angers (France, n = 102), Foljuif (France, n = 104), Lille (France, n = 108), Maison-Estuaire (France, n = 103), Avignon (France, n = 105), Montauban (France, n = 100), and Sovata (Romania, n = 102). The Saclay orchard served as the primary validation site. Each site follows a fully randomized block design. As described in Materials and Methods, we reduced the Saclay sample to 100 individuals by randomly selecting trees to match the other orchards' sample sizes while ensuring representation of all genetic groups.

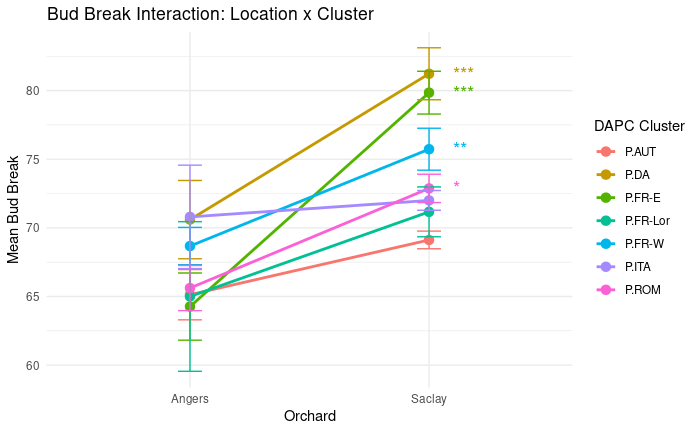

**Suppl. Fig. 17: Genotype-by-environment interaction for bud break timing across DAPC genetic clusters.** Reaction norms illustrating mean bud break dates (points connected by lines) across DAPC-derived genetic clusters evaluated at the Angers and Saclay experimental orchards ($n=86$ individuals per site). Error bars denote $\pm1$ standard error of the mean (SEM). Asterisks indicate statistically significant Location $\times$ DAPC cluster interaction terms derived from a linear mixed-effects model (see Materials and Methods; *p* < 0.05 ~ *, **p* < 0.01 ~ **, *p* < 0.001 ~ ***).

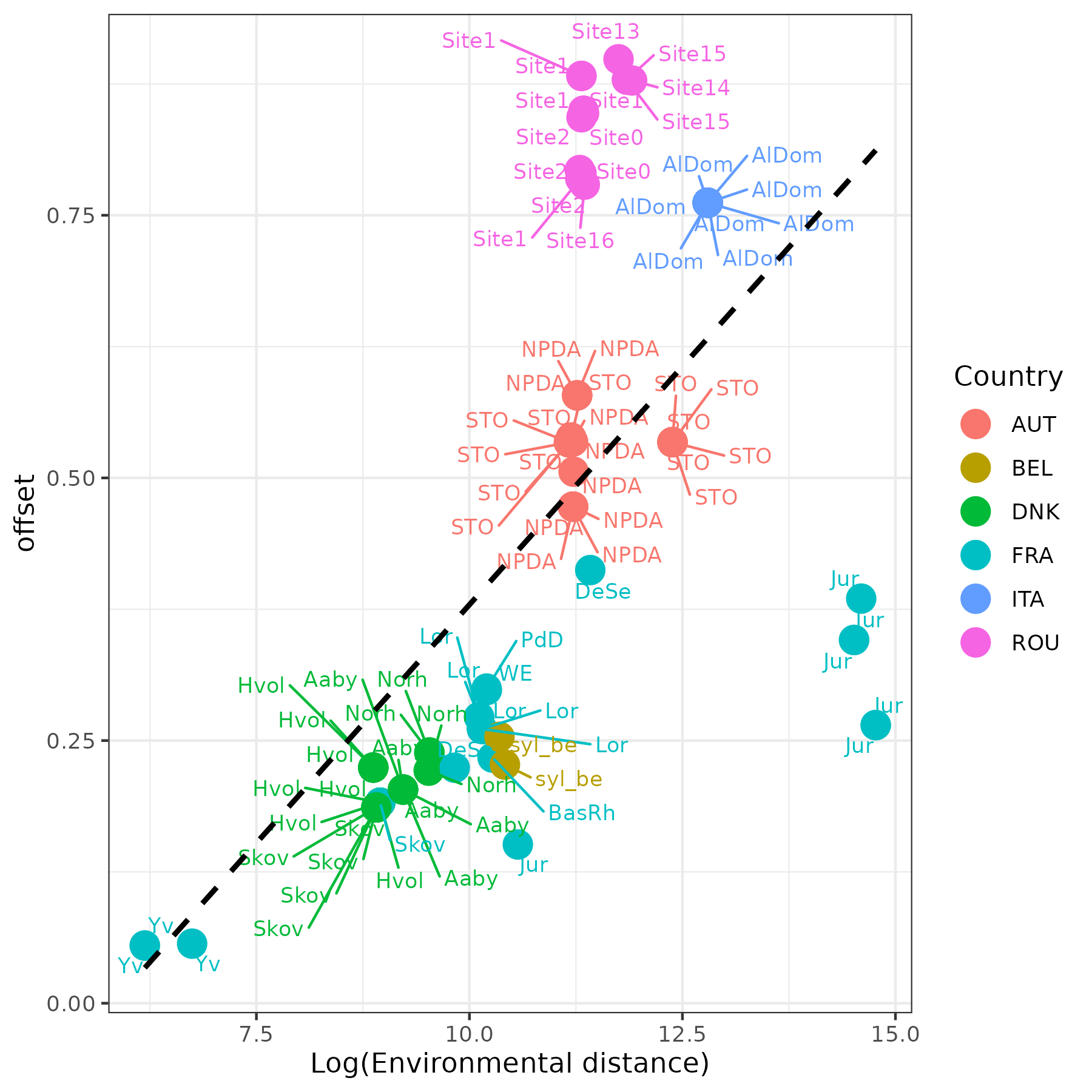

**Suppl. Fig. 18:** **Relationship between genomic offset and environmental distance.** Evaluation of predicted genetic offset for the Saclay orchard target site as a function of the log-transformed environmental distance between individual source environments and Saclay. The dashed black line indicates the overall correlation trend across both axes. Individuals are color-coded by country of origin (AUT: Austria, BEL: Belgium, DNK: Denmark, FRA: France, ITA: Italy, ROU: Romania), with specific sampling site labels indicated by text annotations.

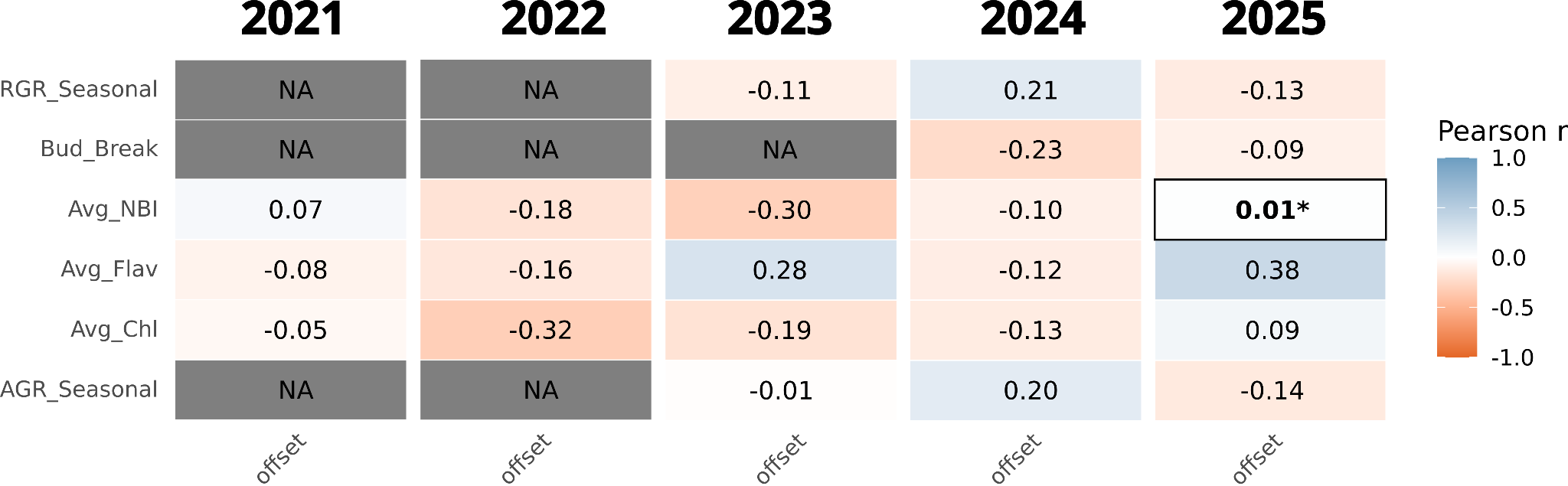

**Suppl. Fig. 19: Annual validation and trait x offset correlation.** Heatmap displaying Pearson correlation coefficients ($r$) between measured phenotypic traits (y-axis) and genomic offset (x-axis), across 5 years of measurement. Color intensity reflects the magnitude and direction of correlation, scaling from deep orange ($r=-1.0$, strong negative) to deep blue ($r=1.0$, strong positive), with white indicating no correlation ($r=0.0$). Evaluated traits include intra-annual relative and absolute growth rates (RGR, AGR), spring phenology (Bud Break), and leaf chemical signatures (Avg_NBI: Nitrogen Balance Index; Avg_Flav: Flavonoids; Avg_Chl: Chlorophyll). Statistically significant interaction terms (Genetic status group x offset) are indicated by asterisks (*). NA correlation reflects missing phenotypic measurement during the corresponding year.

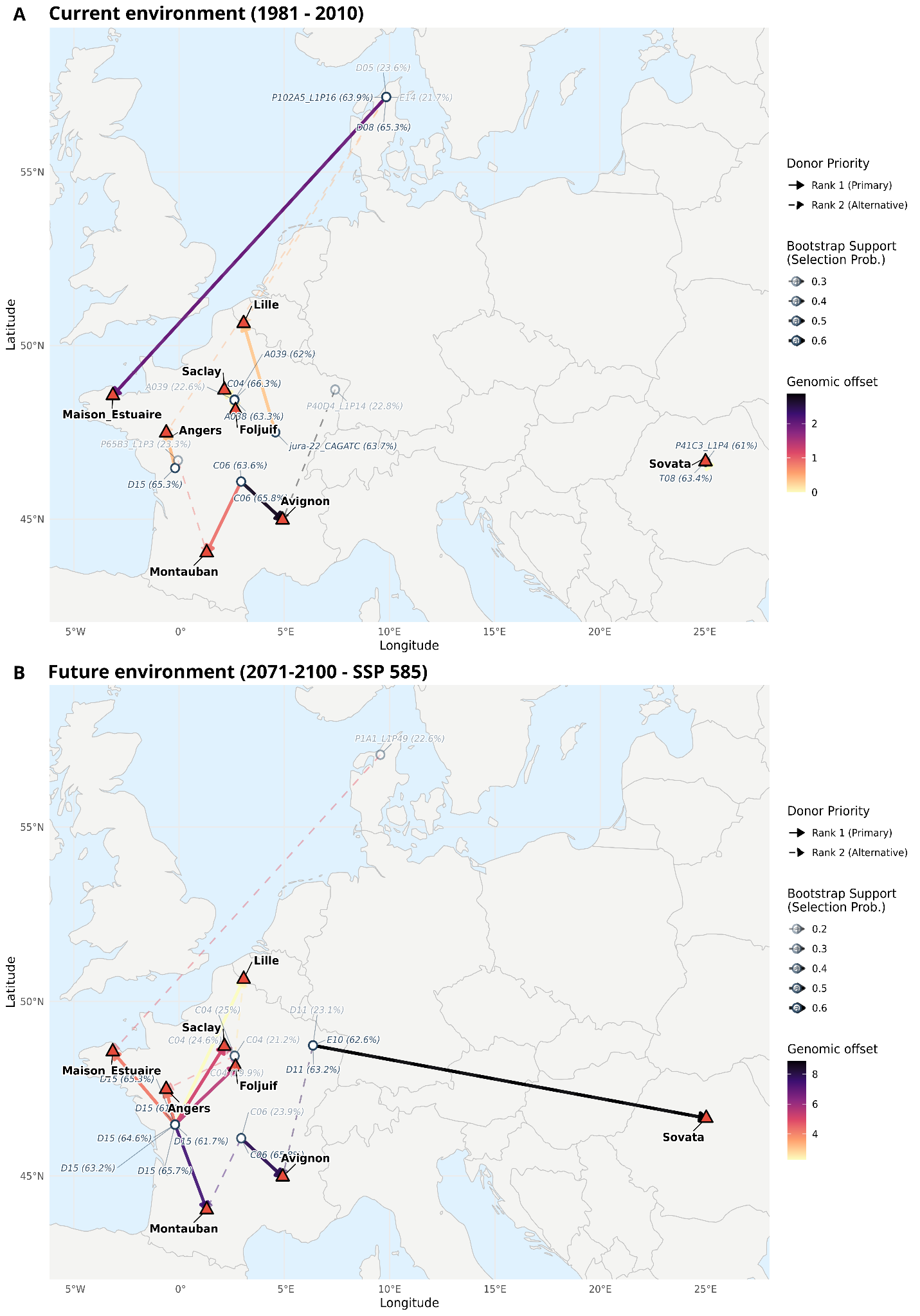

**Suppl. Fig. 20: Spatial donor–recipient translocation vectors and donor selection uncertainty under historical and future climate scenarios.** Arrows represent directional translocation vectors from donor populations (circles) to target destination orchards (triangles), indicating optimal genomic translocation targets rather than physical migratory pathways. Panels display translocations under (A) historical baseline climate conditions (1980–2010) and (B) future projections under the high-emissions SSP5-8.5 scenario (2071–2100). Solid arrows indicate the primary optimal donors (Rank 1) for each recipient site, while dashed arrows represent the top alternative donors (Rank 2). Arrow line width, line transparency, and donor point opacity scale continuously with donor selection probability derived from 1,000 bootstrap iterations, with numerical bootstrap support percentages explicitly reported in the donor labels alongside individual IDs. Vector colors indicate genomic offset values, ranging from bright colors (low offset) to dark colors (high offset). Note that color scales are independently rescaled for each panel—Panel A spans roughly 0–3.5 and Panel B roughly 2.5–7.5—so arrow colors should not be compared directly between panels. Complete donor rankings (Ranks 1–5), selection probabilities, and genomic offsets per orchard site are provided in Suppl. Table 20 (historical, 1980–2010) and Supp. Table 23 (SSP5-8.5, 2071-2100).

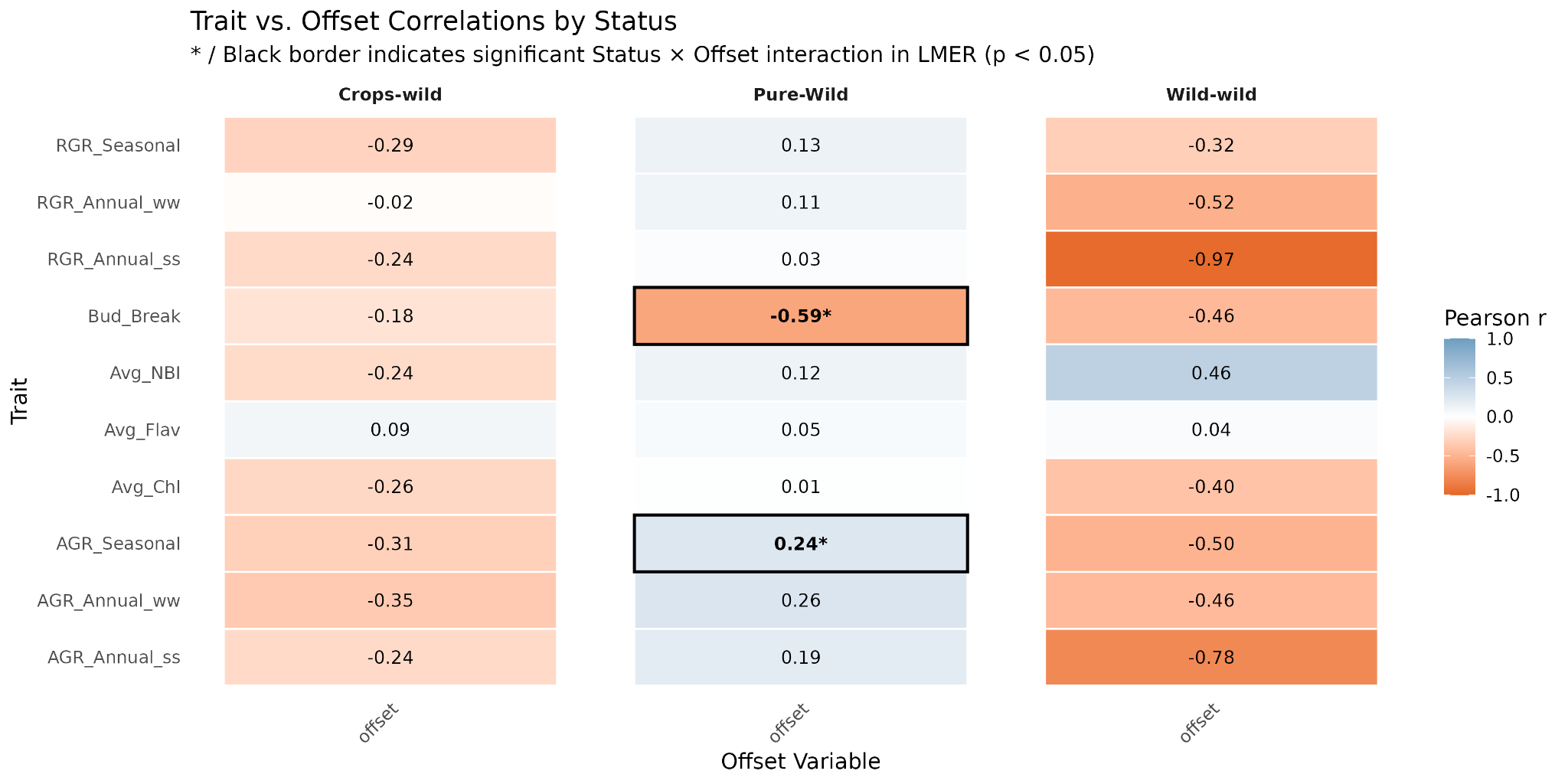

**Suppl. Fig. 21: Admixture history modulates phenotypic vulnerability to genomic maladaptation.** Heatmap displaying Pearson correlation coefficients ($r$) between measured phenotypic traits (y-axis) and genomic offset (x-axis), stratified across three genetic status groups: Crop–wild hybrids (CW, $n=46$), Pure wild (PW, $n=76$), and Wild–wild admixed (WW, $n=13$). Color intensity reflects the magnitude and direction of correlation, scaling from deep orange ($r=-1.0$, strong negative) to deep blue ($r=1.0$, strong positive), with white indicating no correlation ($r=0.0$). Evaluated traits include relative growth rates (RGR), absolute growth rates (AGR), spring phenology (Bud Break), and leaf chemical signatures (Avg_NBI: Nitrogen Balance Index; Avg_Flav: Flavonoids; Avg_Chl: Chlorophyll). Statistically significant interaction terms (Genetic status group x offset) are indicated by asterisks (*). Correlations lacking a significant interaction term (e.g., WW group values for annual RGR/AGR) are considered descriptive given the reduced sample size of the WW category ($n=13$).

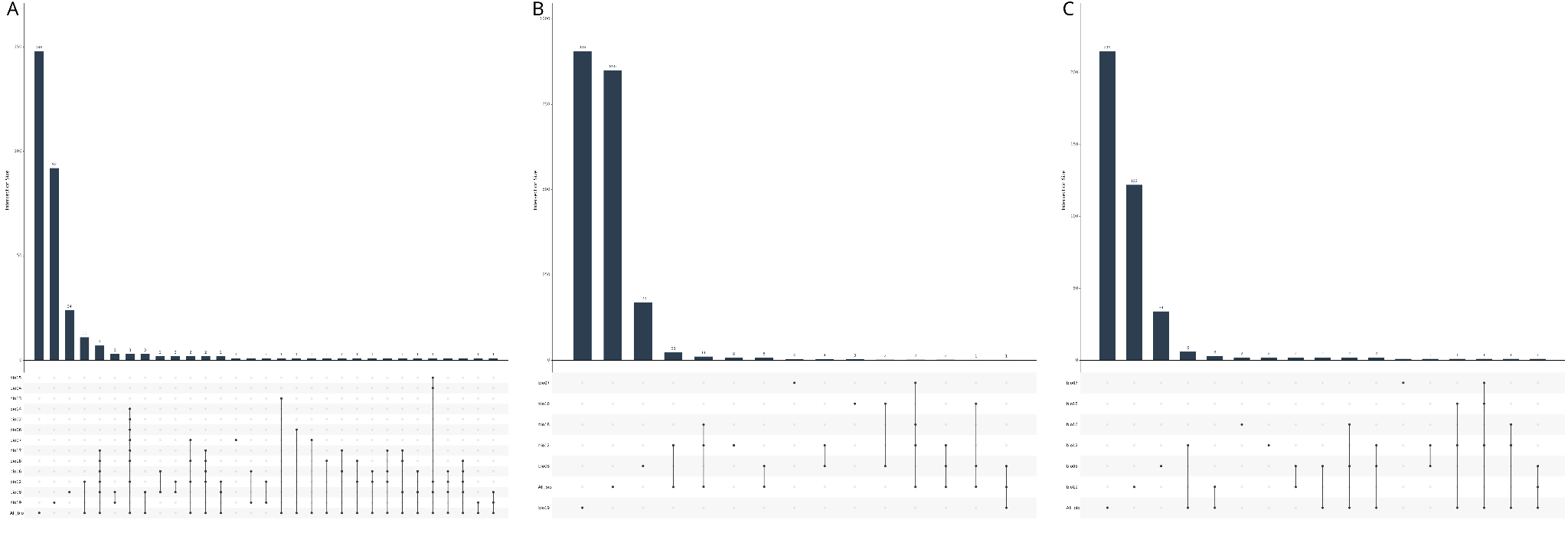
**Suppl. Fig. 22: Distribution and set intersections of climate-associated TIPs, SNPs, and candidate genes across bioclimatic variables.** UpSet plot showing set intersections and overlaps of co-associated transposable element insertion polymorphisms (TIPs; n = 319), single nucleotide polymorphisms (SNPs; n = 1,995), and candidate genes (n = 398) across individual bioclimatic variables and the integrated global model (All_Bio). Retained gene–TIP–SNP triplets meet specific spatial and environmental co-association criteria: TIPs are intergenic variants located within an intergenic span of ≤ 5 kb and situated specifically within the ≤ 2 kb upstream or downstream flanking regions of the candidate gene, while paired SNPs reside directly within the candidate gene transcript body (spanning CDS, exons, 5' UTR, 3' UTR, mRNA, or ncRNA) and share at least one bioclimatic variable association with the TIP. Set intersections reveal the highest counts in the multi-variable model (All_Bio), followed by Bio19 (precipitation of the coldest quarter) and Bio8 (mean temperature of the wettest quarter).

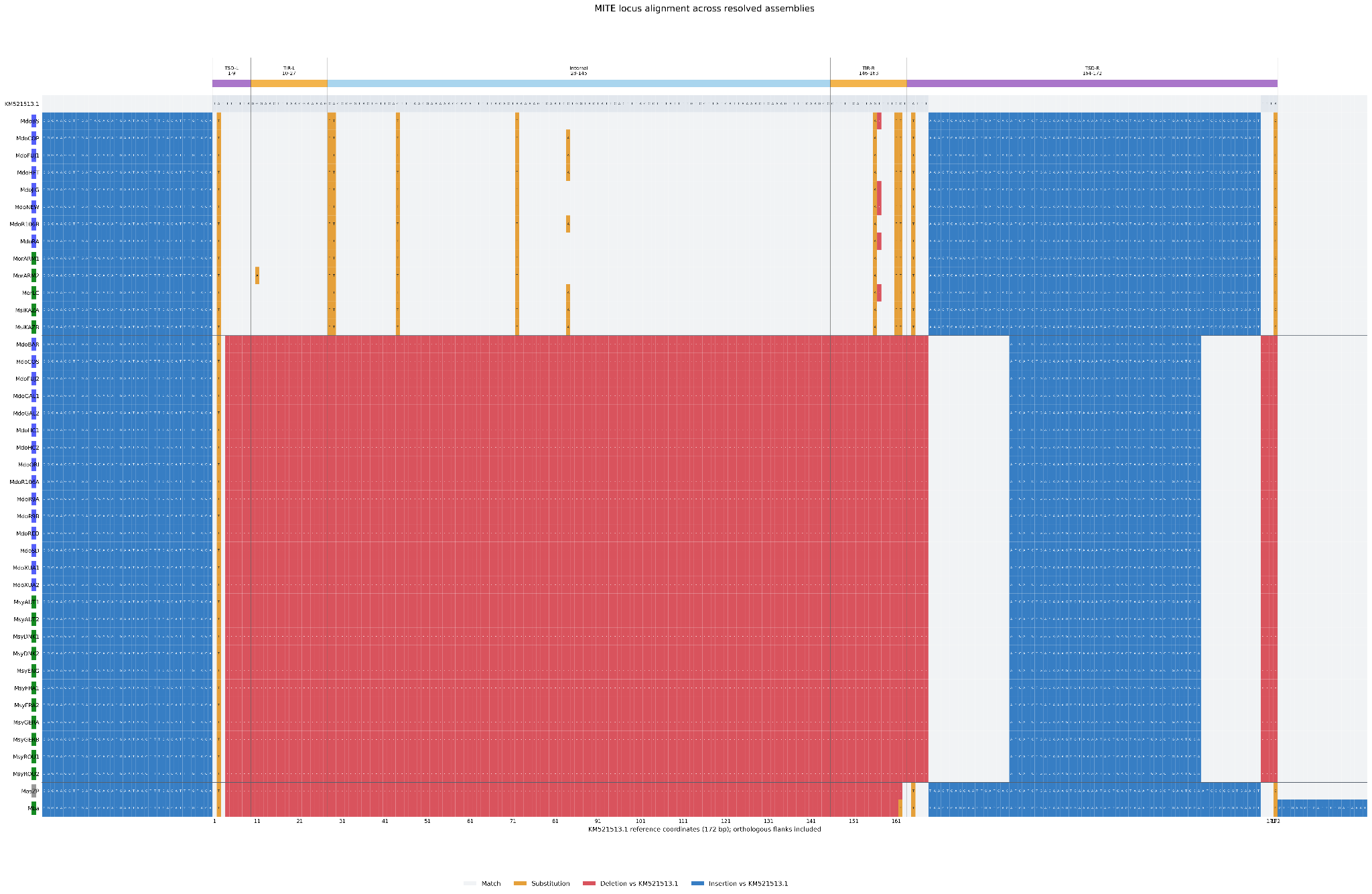

**Suppl. Fig. 23: Sequence alignment and structural features of the MdoRA.svim_asm.INS.2031 MITE insertion.** Alignment of MITE insertion sequences across 13 *Malus* pangenome assemblies against the 173-bp reference sequence (GenBank accession KM521513.1), highlighting nucleotide substitutions (orange), deletions (red), and insertions (blue). Target site duplications (TSDs; 9 bp) and terminal inverted repeats (TIRs; 18 bp) are indicated. Sample ID prefixes denote species identity: *Mdo* (*Malus domestica*), *Mor* (*Malus orientalis*), and *Msi* (*Malus sieversii*). This specific element (MdoRA.svim_asm.INS.2031; chr2:11,288,317) was selected for detailed sequence characterization because it represents the single overlapping gene-proximal transposon insertion polymorphism (TIP) identified between climate association (GEA) and phenotypic association (GWAS) framework analyses.

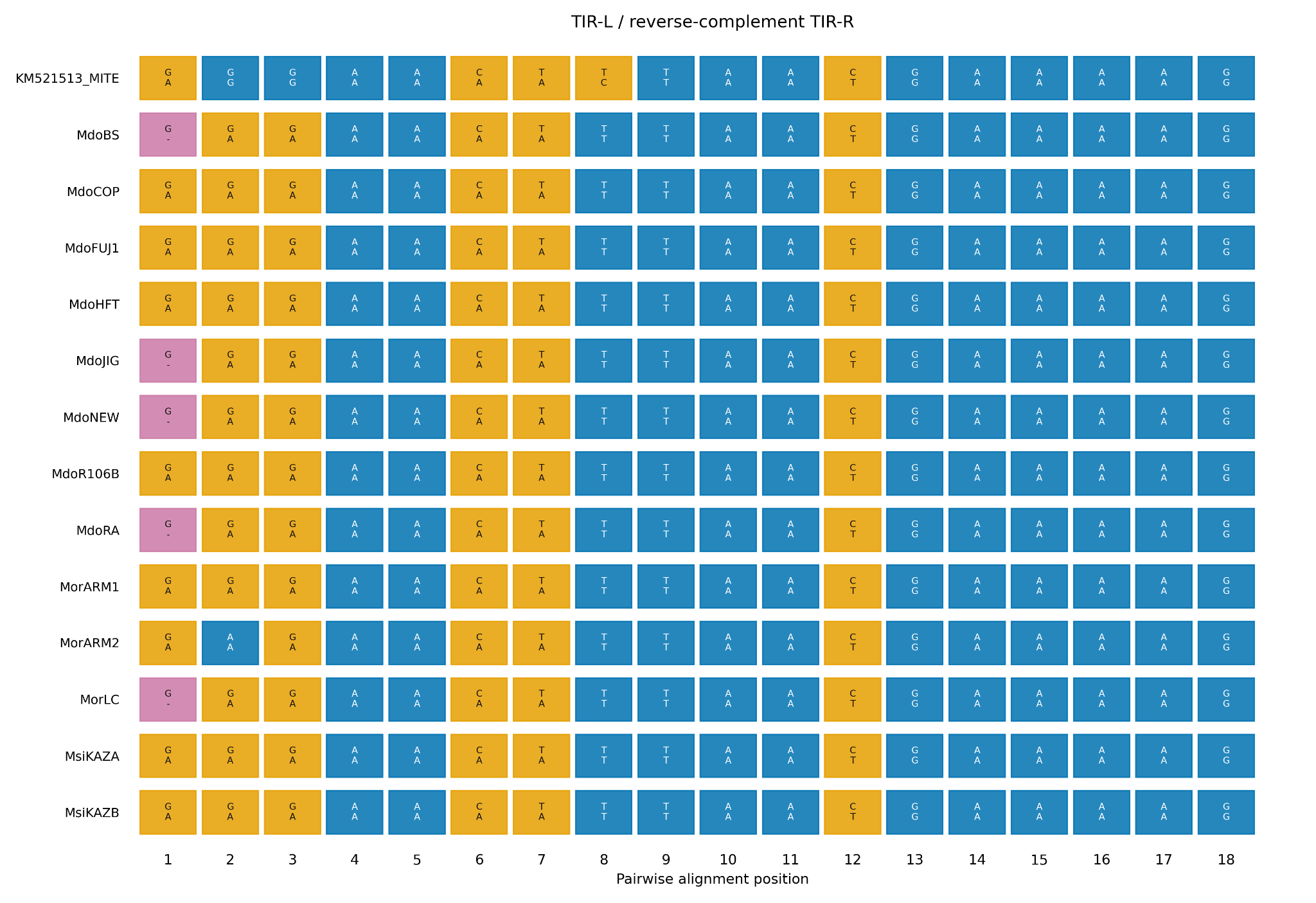

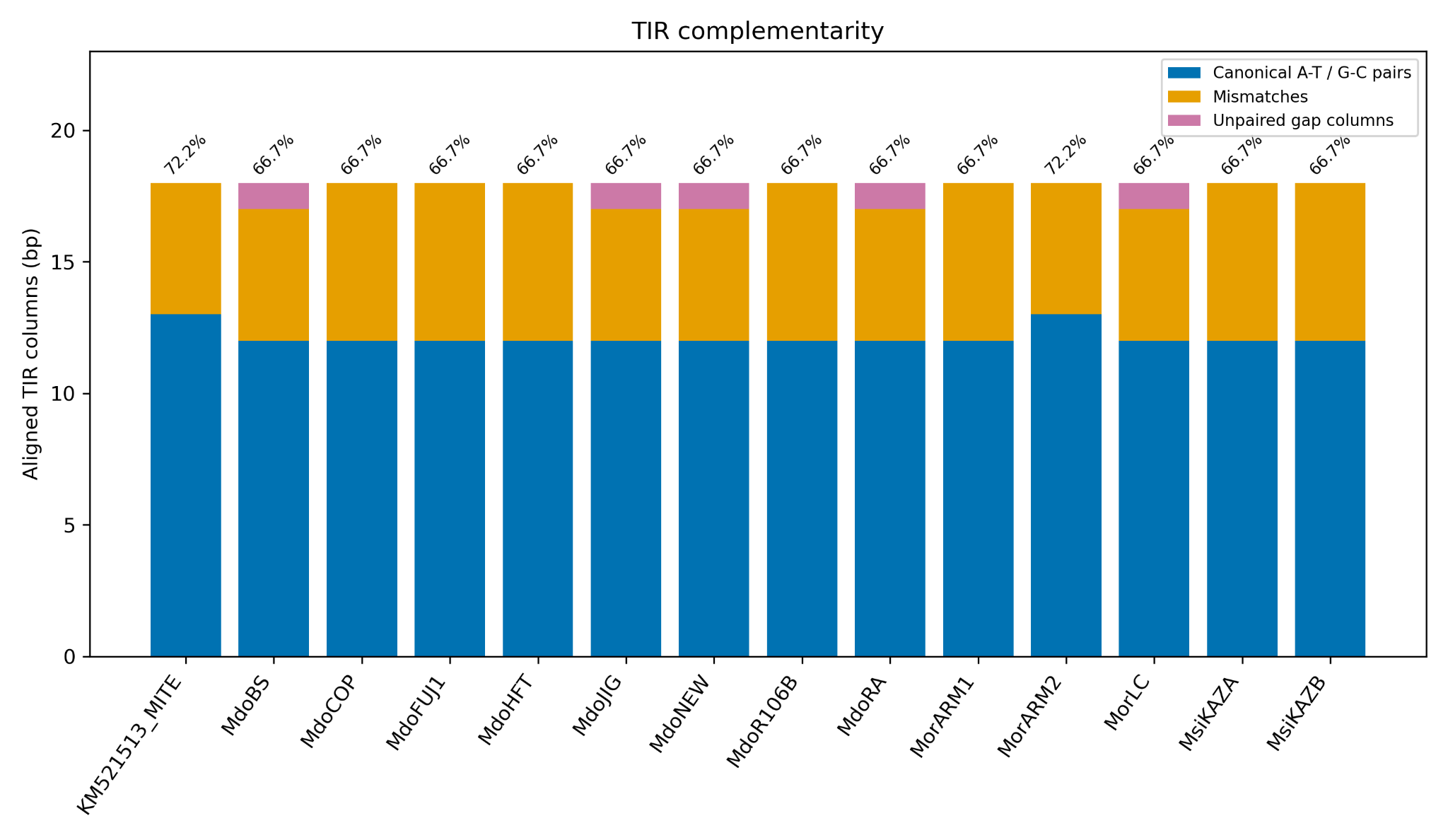

**Suppl. Fig. 24:** **Complementarity between the left TIR and the reverse complement of the right TIR in MITE-containing assemblies.** Canonical Watson–Crick pairs (blue), mismatches (orange), and gaps (purple) are shown at nucleotide resolution and summarized as stacked bars, with percentages indicating canonical pairing.

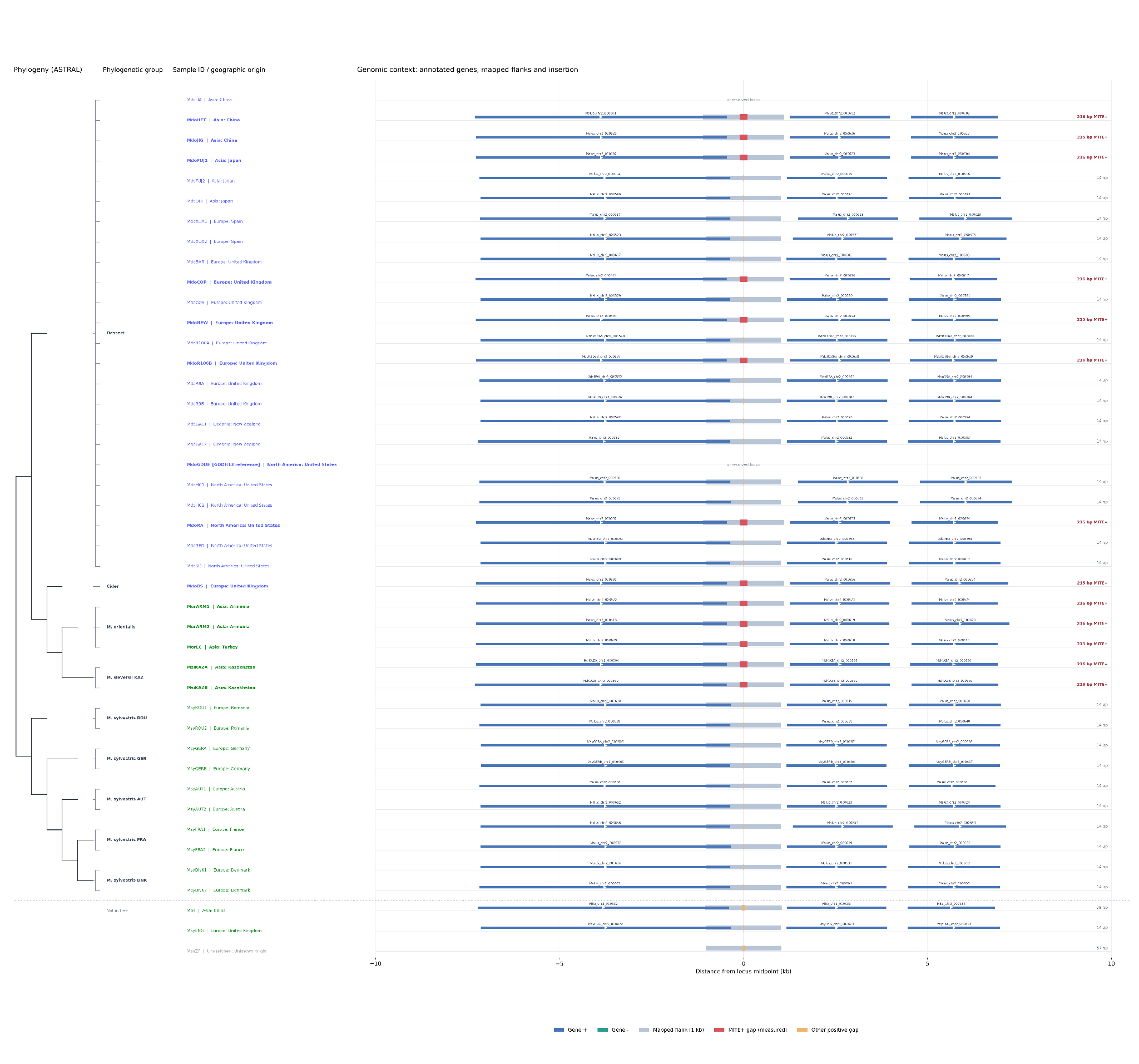

**Suppl. Fig. 25: Genomic context of the MITE locus across *Malus* assemblies, arranged according to an ASTRAL species tree.** Sample labels indicate geographic origin, with cultivated and wild accessions distinguished by color. Annotated genes (±10 kb), mapped flanks, and MITE insertions are displayed relative to the locus center. Assemblies absent from the ASTRAL tree are shown separately.

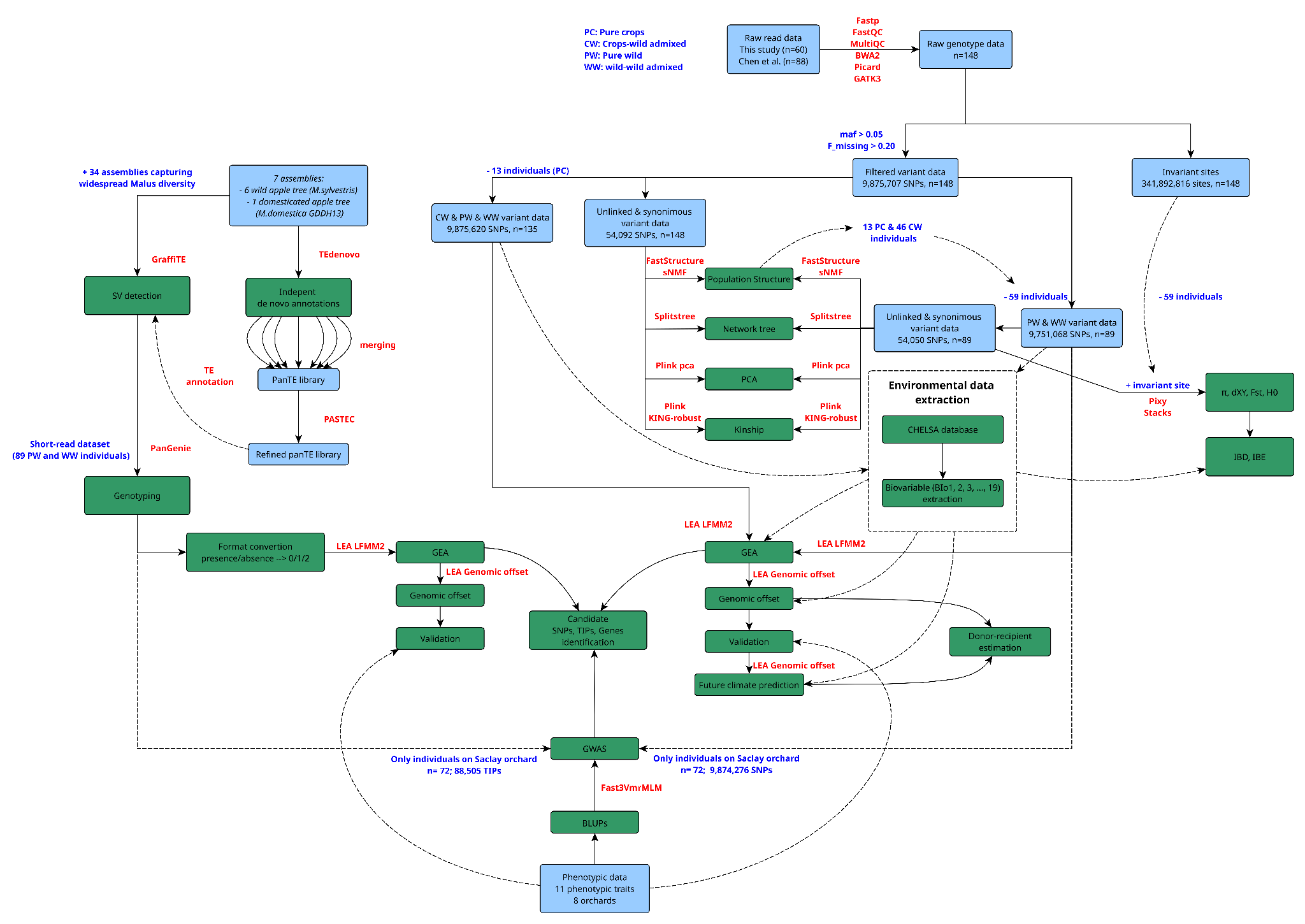

**Suppl. Fig. 26:** **Bioinformatics workflow showing the data origin, the number of individuals, single-nucleotide polymorphisms (SNPs), and transposon insertion polymorphisms at each step, and the methods used in this study.** The blue box indicates the SNPs and TIPs dataset and the filtering step. The green box indicates the different analyses done. Red text indicates the tools used, and blue indicates individual movement between datasets.

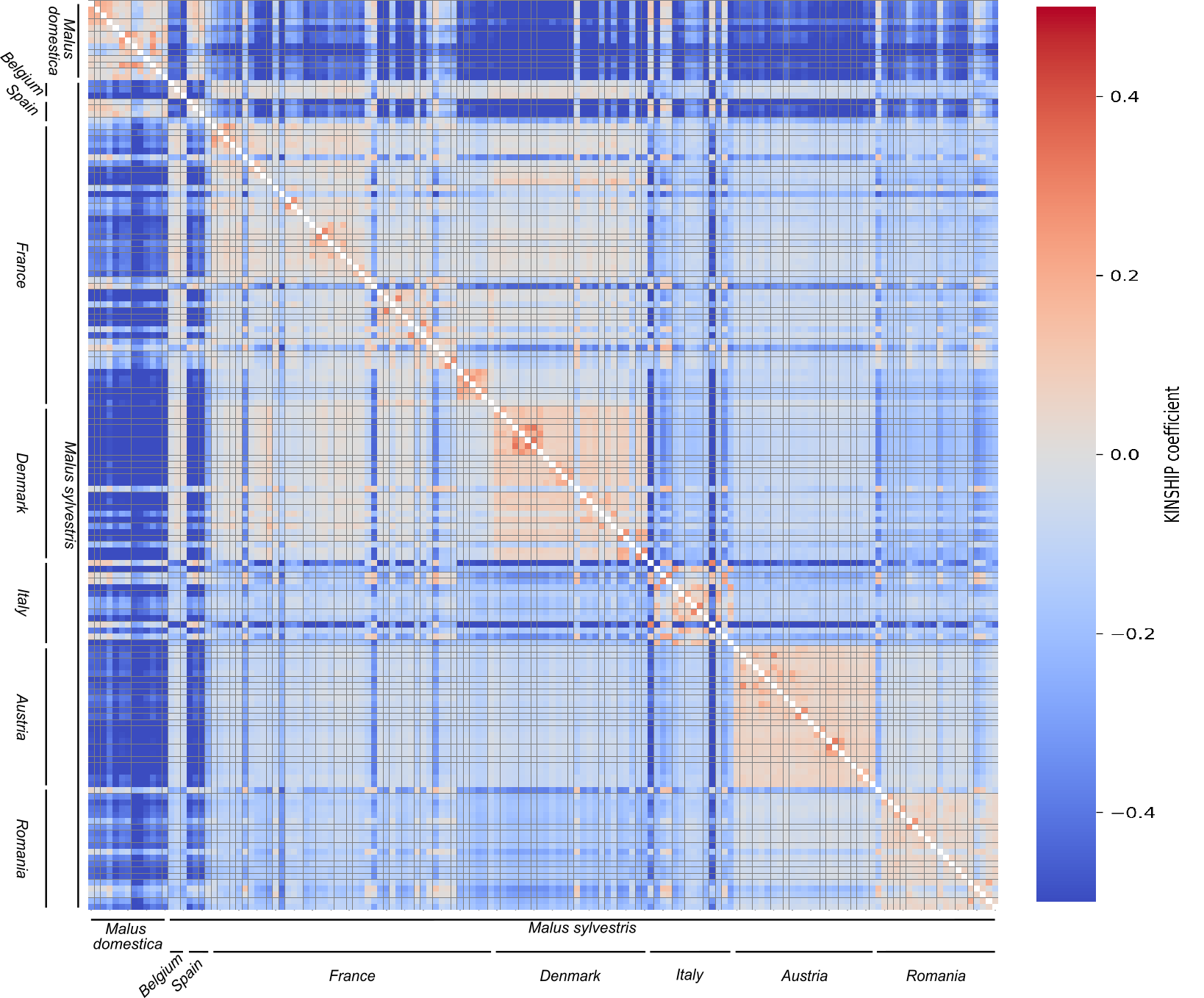

**Suppl. Fig. 27: Heatmap of pairwise kinship coefficients.** Pairwise kinship coefficients ($\phi$) among the 148 sampled individuals, illustrating patterns of relatedness grouped by species and country of origin (rather than inferred genetic clusters; see Suppl. Fig. 22 for cluster assignments). Kinship was estimated using the relationship inference algorithm (KING-robust) implemented in PLINK 2 (1) based on the same unlinked, synonymous SNP dataset used for population structure analysis (54,092 SNPs, $n=148$; see Suppl. Fig. 19). Pairwise relationship classes follow standard KING kinship thresholds: duplicate or monozygotic twin pairs ($\phi>0.354$), first-degree relatives ($0.177<\phi\leq0.354$), second-degree relatives ($0.0884<\phi\leq0.177$), third-degree relatives ($0.0442<\phi\leq0.0884$), and unrelated pairs ($\phi<0.0442$).

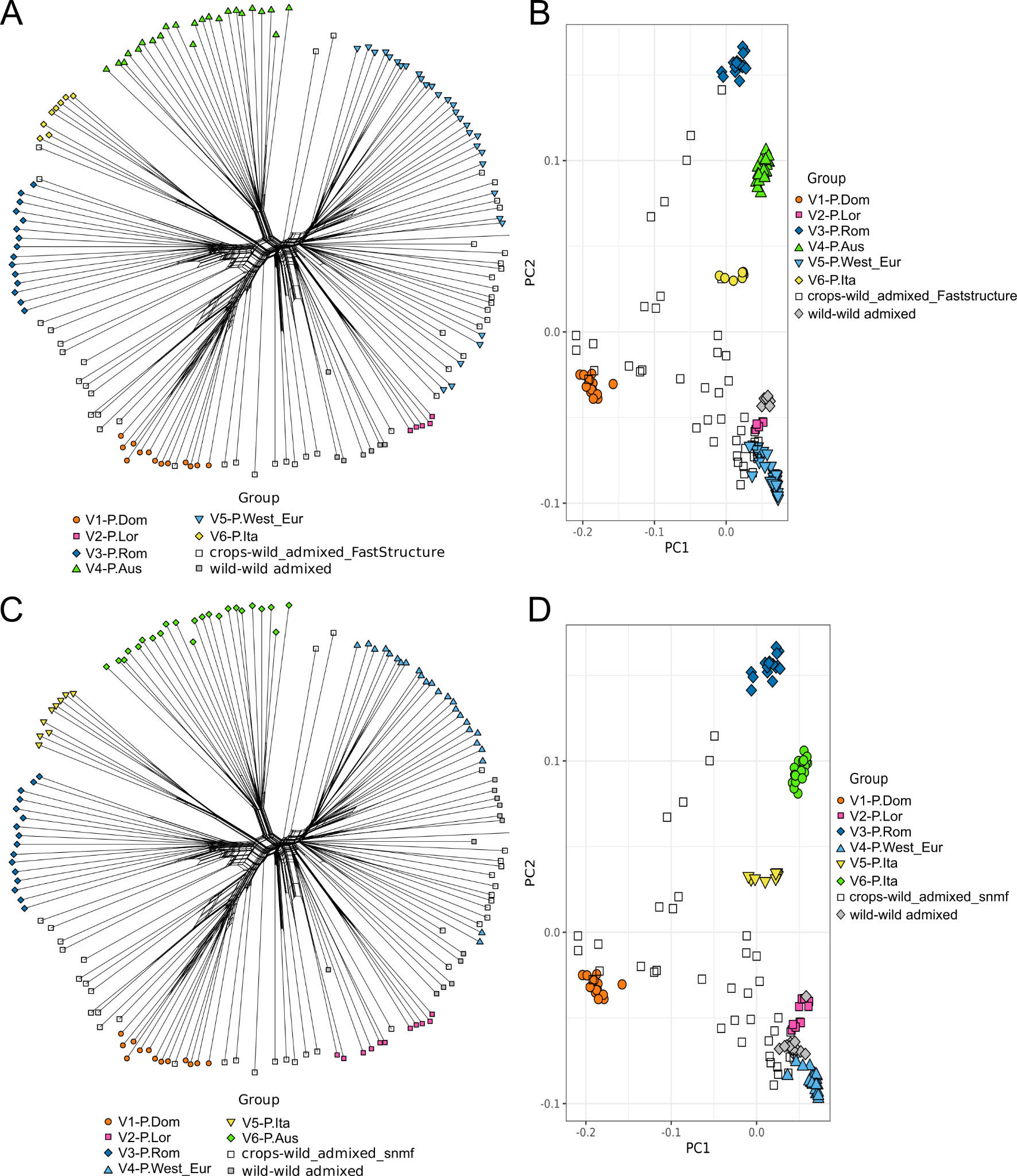

**Suppl. Fig. 28: SplitsTree and principal component analysis (PCA) for the two population structure inference methods applied to the 148 individuals.** **(A)** SplitsTree representation of genetic relationships among individuals, with clusters inferred by fastSTRUCTURE. **(B)** PCA based on fastSTRUCTURE cluster assignments. **(C)** SplitsTree representation showing clusters inferred by sNMF. **(D)** PCA based on sNMF cluster assignments. Cluster colors are as follows: P.Dom (orange), P.West_Eur (cyan), P.Lor (pink), P.Ita (yellow), P.Aus (green), and P.Rom (blue). Crop–wild admixed individuals identified by sNMF and fastSTRUCTURE are shown in white, and wild–wild admixed individuals in grey.

References

1.	Manichaikul A, Mychaleckyj JC, Rich SS, Daly K, Sale M, Chen WM. Robust relationship inference in genome-wide association studies. Bioinformatics. 15 nov 2010;26(22):2867‑73. doi:10.1093/bioinformatics/btq559

2.	[Dadole R, Chen X, Sandoval J, Venon A, Criado M, Delpouve N, et al. Conserved transcriptional plasticity, not local adaptation, dominates early climate responses across wild and cultivated apples [Internet]. 2026 [cité 19 oct 2026]. Disponible sur: http://biorxiv.org/lookup/doi/10.64898/2026.01.15.698562 doi:10.64898/2026.01.15.698562](https://www.zotero.org/google-docs/?v5DfZs)
